# Integrated multi-omics single cell atlas of the human retina

**DOI:** 10.1101/2023.11.07.566105

**Authors:** Jin Li, Jun Wang, Ignacio L Ibarra, Xuesen Cheng, Malte D Luecken, Jiaxiong Lu, Aboozar Monavarfeshani, Wenjun Yan, Yiqiao Zheng, Zhen Zuo, Samantha Lynn Zayas Colborn, Berenice Sarahi Cortez, Leah A Owen, Nicholas M Tran, Karthik Shekhar, Joshua R Sanes, J Timothy Stout, Shiming Chen, Yumei Li, Margaret M DeAngelis, Fabian J Theis, Rui Chen

## Abstract

Single-cell sequencing has revolutionized the scale and resolution of molecular profiling of tissues and organs. Here, we present an integrated multimodal reference atlas of the most accessible portion of the mammalian central nervous system, the retina. We compiled around 2.4 million cells from 55 donors, including 1.4 million unpublished data points, to create a comprehensive human retina cell atlas (HRCA) of transcriptome and chromatin accessibility, unveiling over 110 types. Engaging the retina community, we annotated each cluster, refined the Cell Ontology for the retina, identified distinct marker genes, and characterized cis-regulatory elements and gene regulatory networks (GRNs) for these cell types. Our analysis uncovered intriguing differences in transcriptome, chromatin, and GRNs across cell types. In addition, we modeled changes in gene expression and chromatin openness across gender and age. This integrated atlas also enabled the fine-mapping of GWAS and eQTL variants. Accessible through interactive browsers, this multimodal cross-donor and cross-lab HRCA, can facilitate a better understanding of retinal function and pathology.

## Introduction

The advent of high-throughput single-cell transcriptome technologies has greatly enhanced our exploration of cellular diversity. In particular, it enables the creation of comprehensive atlases for healthy tissues, which are crucial for investigating cellular function and disease mechanisms. In pursuit of these goals, the Human Cell Atlas project (HCA) has coordinated collaborative initiatives to catalog cell types throughout the entire human body ^1, 2^. Atlases released to date include the Human Lung Cell Atlas ^3^ and the Human Breast Cell Atlas^4^.

Within the HCA initiative, the Eye Biological Network aims to create a cell atlas for the human eye. Recent studies have generated atlases of the anterior and posterior segments of the human eye ^5, 6^. Other studies have generated retinal atlases from multiple species, including mouse, chick, macaque, and human ^7–15^. The goal of the work reported here is to augment previous datasets with additional donors, cells, and methods to generate the first version of a comprehensive cell atlas of the human retina. In the future, we plan to expand this effort to encompass the entire eye.

In addition to transcriptomic profiling, the advent of advanced technologies enables the exploration of individual cells in various modalities, such as the Assay for Transposase-Accessible Chromatin with sequencing (ATAC-seq) ^16^. Such large-scale multimodal datasets are crucial in the construction of reference cell atlases as they are essential for identifying rare cell types and understanding mechanisms previously restricted by individual datasets and single modality profiling. Additionally, examining the effects of donor traits on each cell type, e.g., aging, ancestry, and gender, requires a diverse and substantial set of donor samples. However, integrating extensive datasets is computationally challenging, especially with large and complex data ^17, 18^. Consequently, the convergence of substantial data resources, cross-donor investigations, and computational prowess represents an essential paradigm for advancing our comprehension of intricate biological systems and diseases.

This study created a comprehensive multi-omics human retina cell atlas (HRCA) through an integrated analysis of over 2 million snRNA-seq nuclei or cells and over 370,000 snATAC nuclei. The HRCA encompasses over 110 distinct retinal cell types, achieving nearly complete molecular characterization and comprehensive chromatin accessibility. The inclusion of a diverse set of donors revealed molecular changes during aging and between genders at a cellular resolution, shedding light on potential links to diseases. The chromatin profiles enabled an in-depth exploration of regulons and regulatory mechanisms governing cell classes, subclasses, and cell types in the human retina. Furthermore, this integrated atlas facilitated fine-mapping of causal variants, targeted genes, and regulatory mechanisms underlying GWAS and eQTL variants for retinal cell types. Overall, the HRCA provides a valuable resource for both basic and translational research on the retina.

## Results

### Single cell atlas of the human retina

To obtain a comprehensive atlas of cell types in the human retina, we integrated seven publicly available datasets ^7, 15, 19–23^ with newly generated unpublished data (Fig. 1A-B). The integrated dataset totals 2,070,663 single nuclei from 144 samples taken from 52 donors (Supplementary Table 1, 2 and 3). Recovered cells included astrocytes, amacrine cells (AC), bipolar cells (BC), cones, horizontal cells (HC), Müller glia cells (MG), microglia, retinal ganglion cells (RGC), retinal pigment epithelium (RPE), and rods. Annotation of the major classes was performed on individual samples by a coarse label prediction method (Methods). To accommodate the large number of cells, data integration for all cells was employed to facilitate lineage-specific annotations for BC, AC, and RGC, given their complex cell types. The major classes were consistently distributed, except for enriched AC and RGC in several donors from new samples where cell enrichments are performed to increase the proportion of highly heterogeneous classes (AC, BC, and RGC), enabling the annotation of rare cell types (Extended Data Fig. 1A-B and Supplementary Table 4).

**Figure 1.**
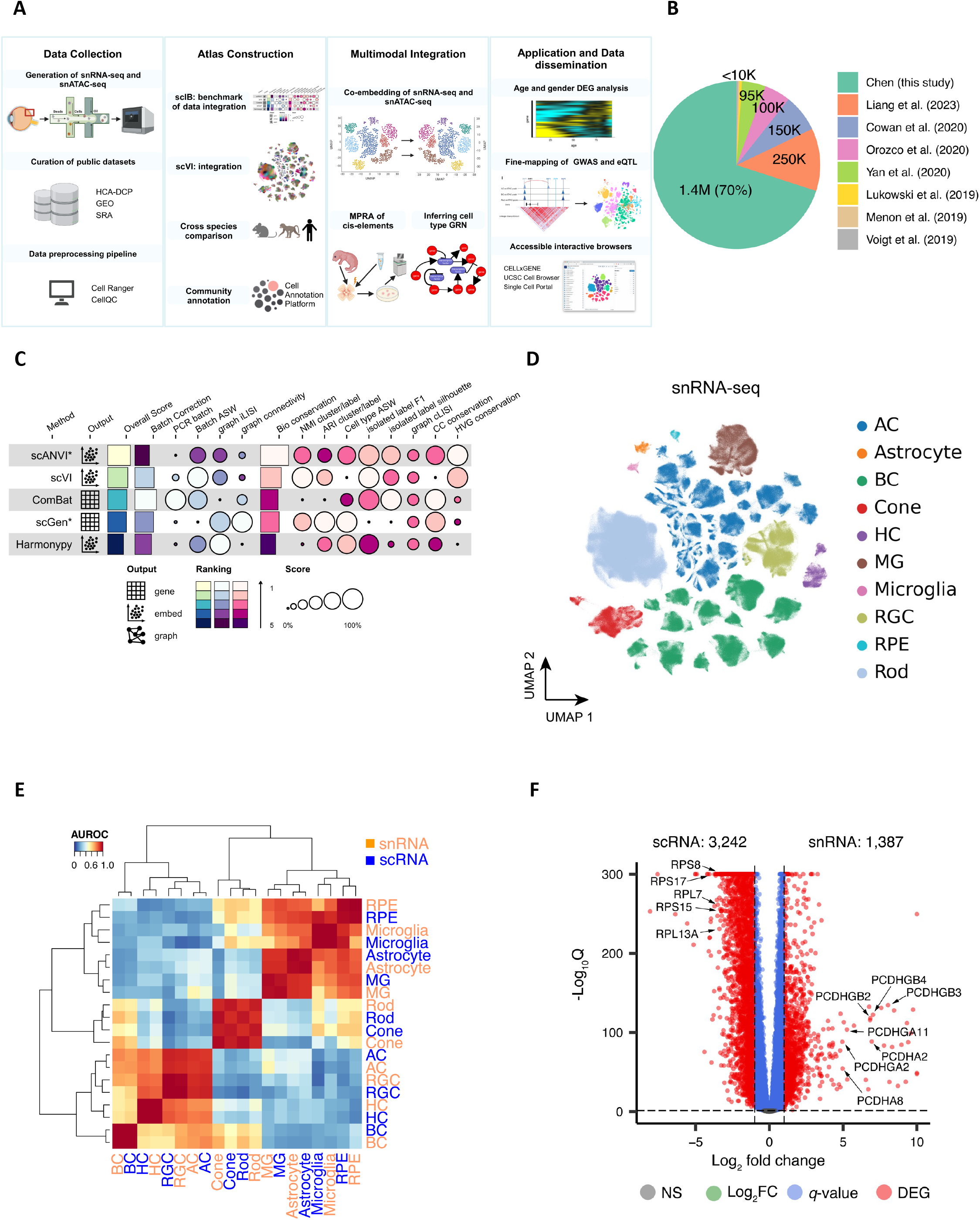
Overview of single cell atlas of the human retina. A. The integrated study for the atlas involves compiling public datasets and in-house generated data, integrating datasets, annotating cell clusters, utilizing chromatin profiles for multi-omics, and demonstrating the utility by applications. B. Collected retinal datasets comprising of both in-house newly generated and seven publicly available datasets. C. Five data integration algorithms are benchmarked for data harmonization. The algorithms are evaluated using 14 metrics, with the rows representing the algorithms and columns corresponding to the metrics. The algorithms are ranked based on their overall score. D. The atlas of snRNA-seq datasets is visualized in a UMAP plot at a major class resolution, with cells colored based on their major classes. E. Cell type similarities of major classes between snRNA-seq (in coral) and scRNA-seq (in blue). The color key is the average AUROC of self-projection for cell types. F. Volcano plot of genes over-expressed in snRNA-seq datasets (on the right) and scRNA-seq (on the left). The x-axis is log2 fold change, and the y-axis is –log10 *q*-value. Differentially expressed genes were identified under |log2 fold change|>1 and *q*- value<0.05 and are depicted as red dots. Selected gene symbols point to the DEGs, including seven genes encoding protocadherin proteins on the right: *PCDHGB2, PCDHGB3, PCDHGB4, PCDHGA2, PCDHA2, PCDHGA11, PCDHA8*; and five genes encoding ribosomal proteins on the left: *RPL7, RPL13A, RPS8, RPS15, RPS17*.

To facilitate the integrated analysis, an scIB approach ^17^ was utilized for benchmarking data integration algorithms, and scVI ^24^ was selected for the construction of the retinal atlas (Fig. 1C, Methods and Supplementary Note). Using scVI, we integrated the entire 2 million cells and embedded them in 2D using UMAP (Extended Data Fig. 1C). We compared the distribution of scRNA-seq and snRNA-seq within this UMAP and found significant differences between snRNA-seq and scRNA-seq transcriptomic signatures, precluding their alignment using scVI (Extended Data Fig. 1C-D). We also benchmarked the conservation of cell type variation when integrating both data types compared to maintaining separate scRNA-seq and snRNA-seq references (Methods and Supplementary Fig. 1). We observed that combining scRNA-seq and snRNA-seq modalities leads to a less accurate representation of cellular variation (Supplementary Fig. 1C). To compare the transcriptome differences, we visualized the 144 samples by averaging the expressions using pseudo-bulk analysis and confirmed that snRNA-seq and scRNA-seq yield distinct transcriptomes (Extended Data Fig. 1E), consistent with previous reports comparing these sequencing modalities ^25^. We therefore created two separate references for snRNA-seq (Fig. 1D) and scRNA-seq (Extended Fig. 1F), respectively. Both were verified by the expression of canonical marker genes for each cell class (Extended Fig. 1G).

To further investigate the transcriptomic differences between the snRNA-seq and scRNA-seq technologies, cell proportions of major classes were calculated and compared in fovea, macular and periphery regions (Supplementary Fig. 2, Extended Data Fig. 2A and Supplementary Note). The most significant differences in cell proportions observed is that scRNA-seq datasets have a higher proportion of MGs compared to snRNA-seq datasets. Cell clusters from these two technologies can be readily aligned as they share similar transcriptomic signatures of major classes (Fig. 1E). However, a large number of differentially expressed genes (DEGs) were identified between the two technologies (Methods and Supplemental Note). In total, 1,387 and 3,242 over-expressed genes were identified across all cell types in snRNA-seq and scRNA-seq datasets, respectively (|log2 fold change| > 1, *q*- value < 0.05) (Fig. 1F and Supplementary Table 5). These over-represented genes exhibited distinct yet biologically related enriched gene ontology (GO) biological processes (Extended Data Fig. 2B-E). For example, genes implicated in biological processes related to ribonucleoprotein complex or ribosome biogenesis, mitochondrial gene expression, and ATP synthesis were enriched in scRNA-seq datasets.

### Bipolar cells

Over 422,000 bipolar single nuclei included in the current atlas can be divided into 14 cell types based on marker genes ^7, 9^ (Fig. 2A). One significant difference from previous reports is that the giant bipolar (GB) and blue bipolar (BB) are separated into two distinct clusters, primarily due to a significant increase in the cell number (Fig. 2B). To facilitate the annotation of BC clusters, we conducted a cross-species analysis to align human BC clusters with mouse and macaque BC types, leveraging both single-cell transcriptomes and protein sequence embeddings with SATURN ^26^ (Fig. 2C-D). High concordance with one-to-one mapping was observed among the three species, consistent with the previous report ^7, 9^. Based on the co-embedding, the human cluster mapped with mouse cell type BC9 is annotated as the BB as BC9 has been reported to exclusively contact S-cones^9^, also known as “blue” cones in humans and macaques^12^, while the human cluster mapped with BC8 is annotated as GB. Despite high similarity between GB and BB, 341 genes highly expressed in GB cells, and 887 genes highly expressed in BB cells were identified (Extended Data Fig. 3D, Supplementary Table 6, Supplementary Fig. 3A-B, and Supplementary Note). Among them, *AGBL1* and *SORCS3* showed high specificity for the GB and BB cells, respectively (Fig. 2A, Extended Data Fig. 3B, and Supplementary Fig. 3C). Consistently, 14 BC corresponding clusters were also observed from the scRNA-seq dataset (Supplementary Table 1 and Extended Data Fig. 3A-C). Furthermore, DEGs in GB and BB, including *AGBL1* and SORCS3, were confirmed by the scRNA-seq (*p*-value<10^-6^), showing a 58% overlap in GB and a 12% overlap in BB (Fig. 2A and E, Extended Data Fig. 3B, Supplementary Fig. 3C, and Supplementary Table 7).

**Figure 2.**
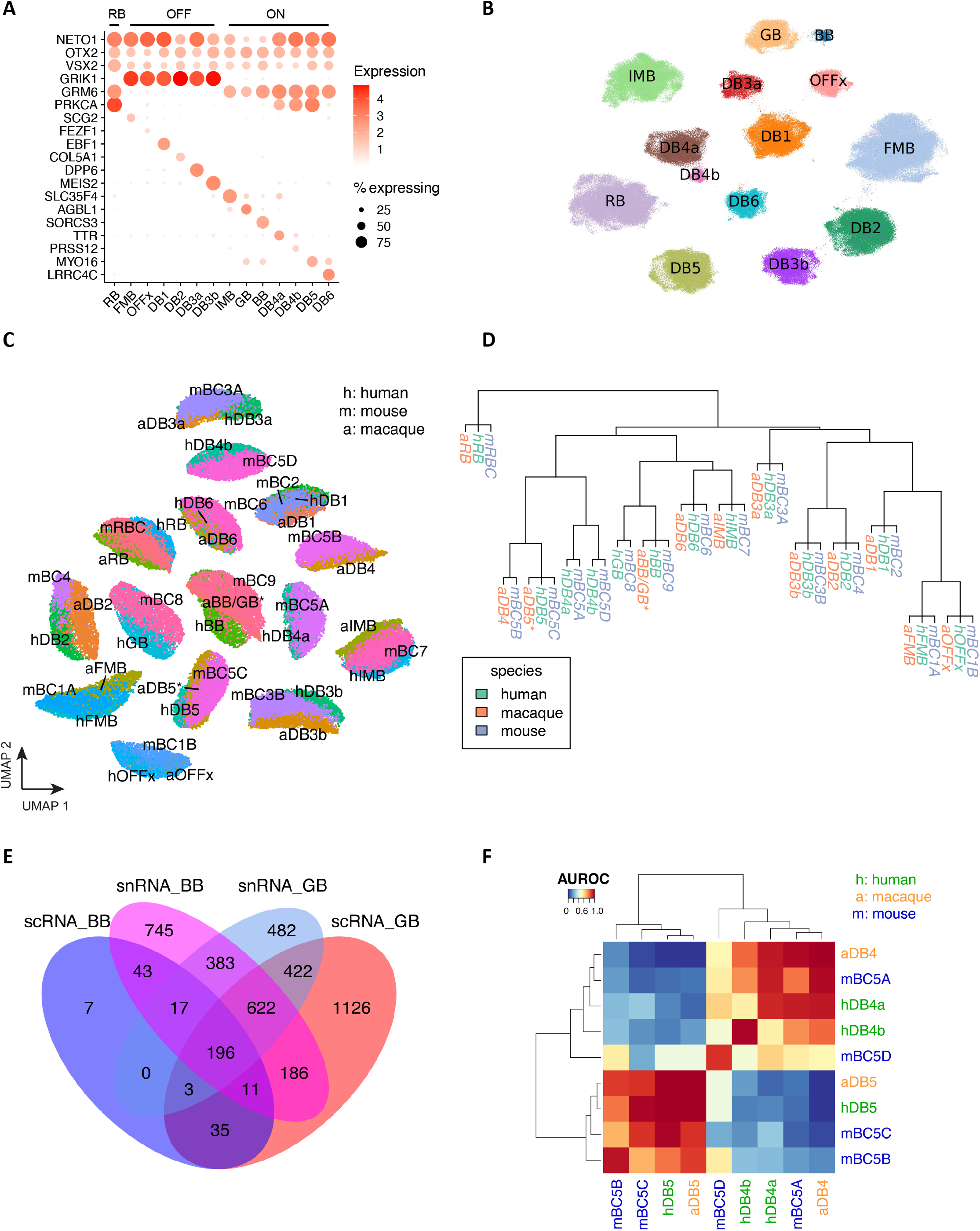
Bipolar cells. A. Distribution of marker genes for BC types. BC subclasses are in RB, OFF and ON. NETO1, OTX2, and VSX2 were used as BC pan-markers. GRIK1 and GRM6 were used as OFF and ON markers, respectively. Rows represent marker genes, and columns represent BC types. The names of BC types are extracted from macaque BC types. B. UMAP visualization of human BC cells. Cell clusters are colored by the annotated cell types. C. Co-embedding of human, mouse, and macaque BC cells. To differentiate between cell types from three species, prefixes were added to the names: “h” for human, “m” for mouse, and “a” for macaque. D. Hierarchical clustering of mouse BC cell types. Expanded leaf nodes are the correspondent cell types from human and macaque BC cell types. E. The overlap between the top-ranked genes of human GB and BB is examined using snRNA-seq and scRNA-seq datasets. Fisher’s exact test was used to calculate the significance of the overlap of top ranked genes in GB (*p-* value=7.5×10^-293^) and BB (*p-*value=1.7×10^-131^) between snRNA-seq and scRNA-seq. F. Cell type similarities among mouse BC5A, BC5B, BC5C, and BC5D, and mapped types in humans and macaques.

In mice, four BC5 types have been identified: BC5A, BC5B, BC5C, and BC5D ^12^. However, how these four closely related BC types correlate with BCs in primates has not been fully resolved. Previously, only BC5A in mice exhibited a confident mapping to DB4 in macaques ^9^. As shown in Fig. 2F, two human BC types, DB4a and DB4b, are closely related to BC5A in mice and DB4 in macaques, while BC5B and BC5C in mice appeared most similar to human and macaque DB5. However, the mouse BC5D appears to be an outlier without closely related BC type in primate. To distinguish the BC types, we identified a set of 55 gene markers that shows robust performance (Extended Data Fig. 3E, Supplementary Table 8 and Supplementary Note).

### Amacrine and retinal ganglion cells

A total of 73 AC types was identified among over 380,000 AC nuclei (Fig. 3A, Extended Data Fig. 4A-B, and Supplementary Table 9), nearly doubling the number of types in a previous study ^7^. Two AC pan-markers, *PAX6* and *TFAP2B*, were confirmed to be highly expressed in these 73 types (Extended Data Fig. 4A). By utilizing makers for GABAergic ACs (the GABA-synthetic enzymes *GAD1* and *GAD2*) ^15^ and Glycinergic ACs (the glycine transporter *SLC6A9*), we identified 55 GABAergic AC types, accounting for ∼65% of ACs, and 11 Glycinergic AC types, accounting for ∼23% of ACs. Seven clusters expressed both markers, classifying them as the “Both” AC types, as previously described in mice ^14^. Based on expression of additional previously characterized markers ^9, 15, 27^, 14 of the 73 AC clusters could be annotated as known AC types (Extended Data Fig. 4C-D, Supplementary Fig. 4A and Supplementary Note). For example, two clusters (HAC10, HAC26) were annotated as Starburst AC (SAC) by *CHAT* and ON-SAC/OFF-SAC by *MEGF10* and *TENM3*, respectively. A set of gene markers to distinguish these 73 AC clusters are identified (Fig. 3B and Supplementary Table 8). To further annotate AC types, a cross-mapping approach was utilized to map the identified AC types with external sources with an existing labeling from public datasets and other species such as macaques and mice (Extended Data Fig. 5A-C, Supplementary Table 9, Supplementary Fig. 5A-C, and Supplementary Note). As expected, high concordance between snRNA-seq and scRNA-seq is observed: 92% (23/25) scRNA-seq clusters can be mapped to this dataset ^15^. Similarly, 94% (32/34) macaque AC types ^9^ mapped to the human dataset. In contrast, only 83% (52/63) mouse AC types mapped to humans ^14^, including four non-GABAergic non-Glycinergic (nGnG) types in mice ^14, 15^ to human clusters (3 Glycinergic and 1 GABAergic) (Supplementary Table 9). Eight human clusters (5 GABAergic and 3 Both) do not have a clear correspondence to previously annotated types. All of these clusters appear to be rare cell types, with the most abundant of them comprising only 0.18% of the AC population (670 nuclei).

**Figure 3.**
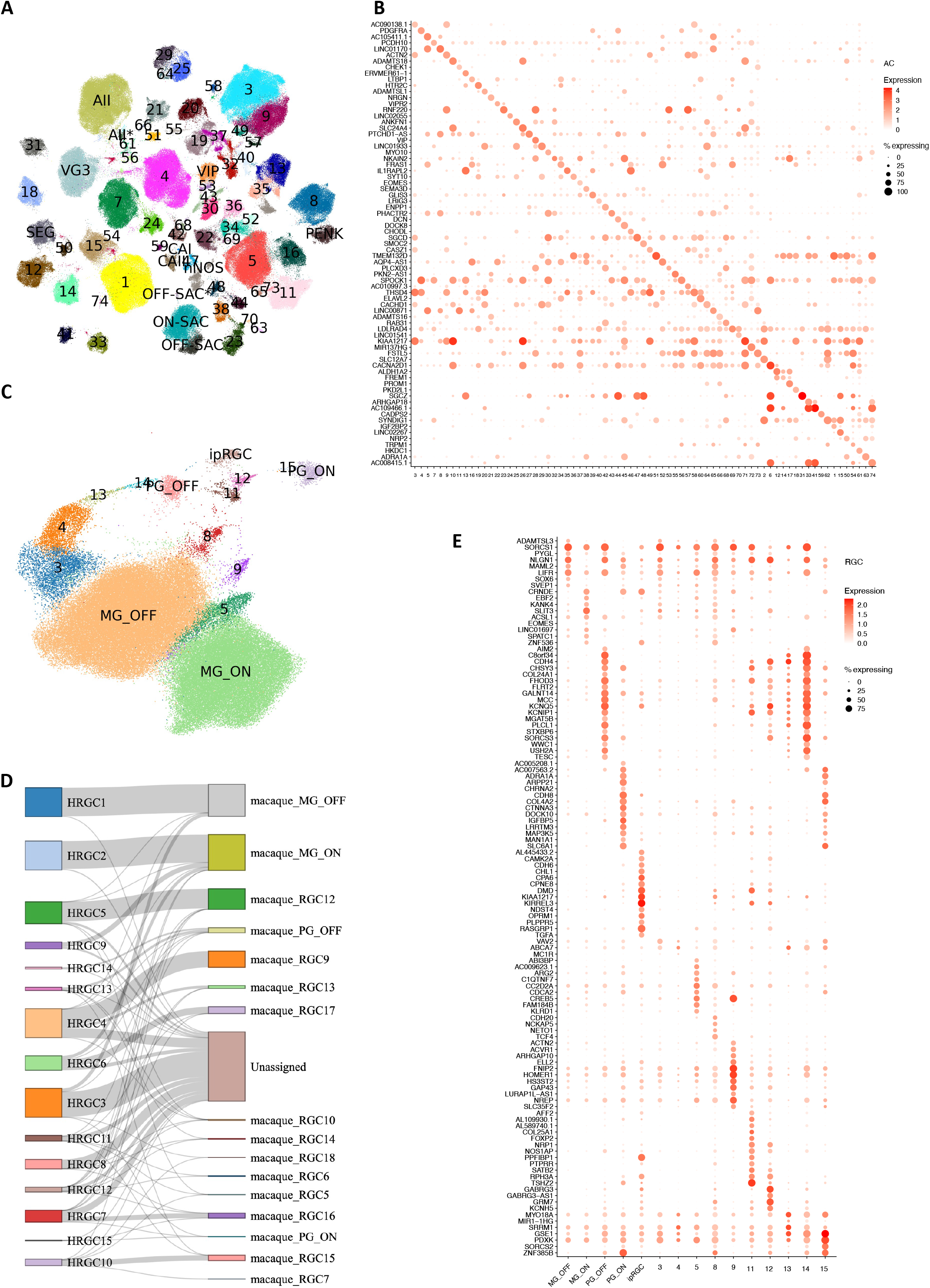
Amacrine cells and retinal ganglion cells. A. UMAP visualization of the identified 73 AC cell clusters. Cluster IDs are placed on top of clusters, and cells are colored by the cluster IDs, where 14 clusters have annotated types. B. Dot plot of predicted markers for AC cell types. C. UMAP visualization of RGC cell types with labels on top of cells. D. Sankey diagram illustrating RGC types alignment between humans (left column) and macaques (right column). E. Dot plot of predicted markers for RGC cell types.

We identified 15 RGC clusters are identified from over 99,000 RGC nuclei included in the atlas (Fig. 3C and Supplementary Table 9). Utilizing previously characterized markers from macaque, five clusters can be annotated (Extended Data Fig. 6A), OFF midget RGC (MG_OFF) by *TBR1* (HRGC1), ON midget RGC (MG_ON) by *TPBG* (HRGC2), OFF parasol RGC (PG_OFF) by *FABP4* (HRGC6), ON parasol RGC (PG_ON) by *CHRNA2* (HRGC7), and an intrinsically photosensitive RGC (ipRGC) by *OPN4* (HRGC10). Consistent with previous findings, the distribution of RGC types in human is highly skewed, with midgets accounting for 87.9% of all RGCs. Parasol RGCs, which accounts for 1.8%, are relatively low compared to previous reports ^9, 15^ due to experimental enrichments (Extended Data Fig. 6B). Cross-species comparisons among humans, macaques and mice reveal that RGC types are highly divergent (Fig. 3D and Extended Data Fig. 6C-D, Supplementary Fig. 6A-C, Supplementary Table 9, and Supplementary Note). As primate RGC types (approximately 18 types) ^28^ are significantly less diverse compared to mouse RGCs (45 molecularly distinct types) ^13^, making it challenging to perform cell cluster mapping between humans and mice (Supplementary Table 9 and Extended Data Fig. 6D). Lastly, a set of novel markers for RGC clusters are identified using the binary classification approach (Fig. 3E and Supplementary Table 8).

### HRCA: chromatin accessibility landscape

To decipher the gene regulatory programs for retinal cell types, 372,967 snATAC nuclei from 52 samples of 26 donors were profiled (Supplementary Table 10 and 11). These nuclei were classified into six neuronal and three glial classes (Fig. 4A-B). Expression of genome-wide genes including canonical marker genes was highly correlated with local chromatin accessibility and inferred gene activity in all cell classes (Fig. 4C, Extended Data Fig. 7A).

**Figure 4.**
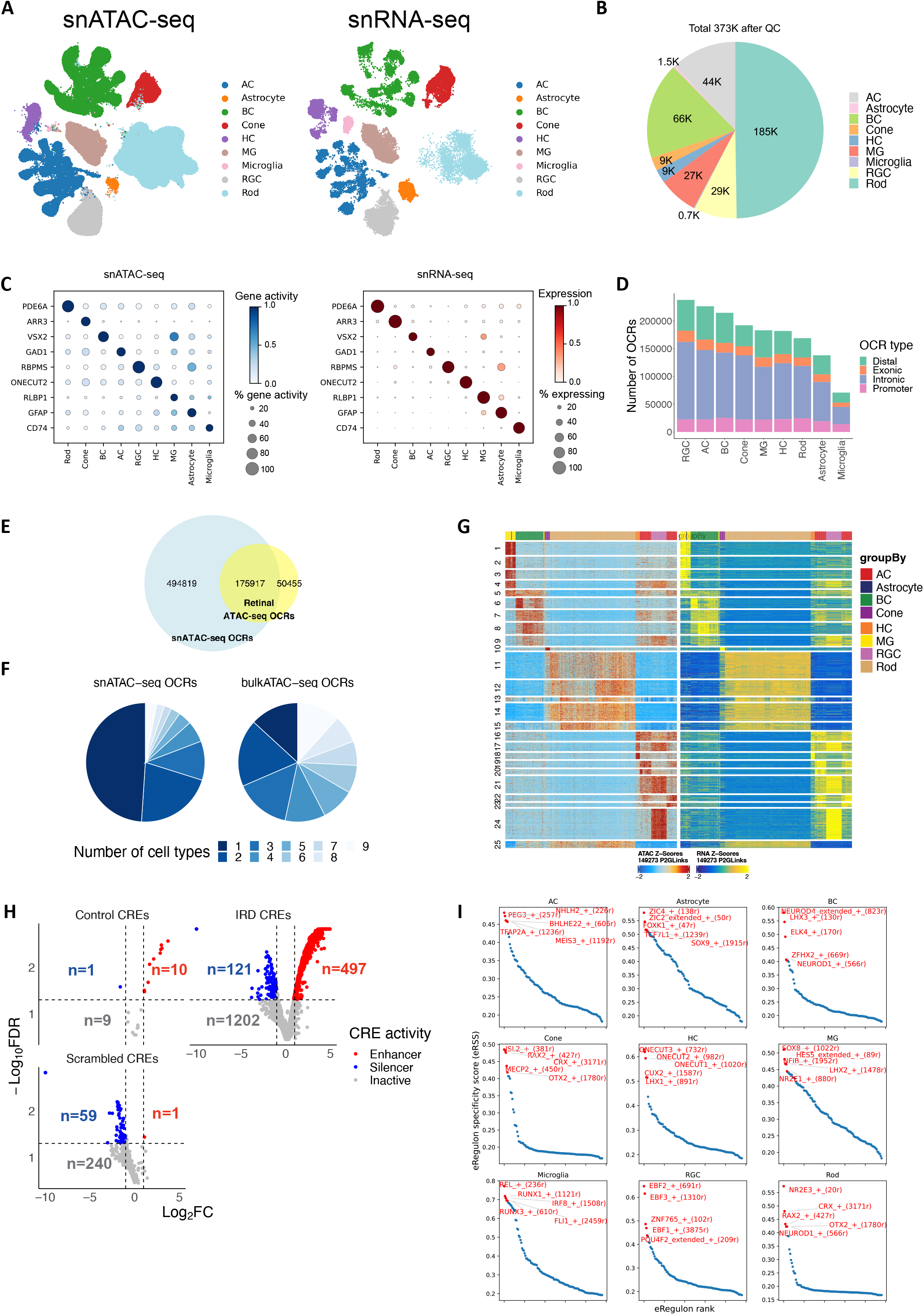
A high resolution snATAC-seq cell atlas of the human retina. A. Uniform Manifold Approximation and Projection (UMAP) of co-embedded cells from snRNA-seq and snATAC-seq showing cells are clustered into major retinal cell classes. B. Pie chart showing the cell proportion distribution of major retinal cell classes in this study. C. Dot plot showing marker gene expression measured by snRNA-seq and marker gene activity score derived from snATAC-seq are specific in the corresponding cell class. D. Bar plot showing the number of open chromatin regions (OCRs) identified in each major cell class. E. The Venn Diagram showing the overlapped OCRs detected by retinal snATAC-seq and bulk ATAC-seq. F. Pie chart showing cell type specificity of OCRs identified from retinal snATAC-seq (left) and bulk ATAC-seq (right). The color codes the number of cell types where the OCRs were observed. G. Heatmap showing chromatin accessibility (left) and gene expression (right) of 149,273 significantly linked CRE-gene pairs identified by the correlation between gene expression and OCR accessibility. Rows represented CRE-gene pairs grouped in clusters by correlations. H. Volcano plot showing the log_2_^*FC*^ value (comparison between activity of each tested sequence and the activity of a basal CRX promoter, X axis) and the −log_10_^*FDR*^ value (Y axis) of each tested sequence by MPRAs (IRD CREs n=1,820, control CREs with a variety of activities n=20, Scrambled CREs n=300). Each dot corresponds to a tested sequence, colored by the activity of the sequence. I. Scatter plot showing the eRegulon specificity score for each transcription factor (TF) and the corresponding regulon across major retinal cell classes. The top five TF and eRegulon are highlighted in red.

Based on this dataset, 670,736 open chromatin regions (OCRs) were identified, with 70,909 to 237,748 OCRs per cell class (Fig. 4D, Supplementary Table 12). To evaluate the quality of these OCRs, we compared them with the OCRs detected by retinal bulk ATAC-seq. The snATAC-seq OCRs captured most (77.7%) of OCRs detected by bulk ATAC-seq. More importantly, many cell class-specific OCRs absent from bulk ATAC-seq analysis were present in the snATAC dataset, resulting in a three-fold increase in the total number of OCRs (Fig. 4E-F). Although many OCRs are shared among multiple cell classes, 4.14% to 24.4% (9,361 to 24,338) of the OCRs per cell class showed differential accessibility depending on cell classes, suggesting potential roles in cell class-specific gene regulation; we refer to these OCRs as differentially accessible regions (DARs) (Extended Data Fig. 7B-C). By calculating the correlation between gene expression or promoter accessibility and chromatin accessibility of surrounding OCRs (-/+250kb), 162,481 linked OCR-gene pairs were identified (Fig. 4G). These linked OCRs are candidate cis regulatory elements (CREs) and the linked genes are likely to be the targets of the CREs. To further validate these putative CREs, particularly those potentially associated with human disease, we conducted massively parallel reporter assays (MPRAs) ^29, 30^ on 1,820 CREs that were linked to inherited retinal disease (IRD) genes, utilizing the mouse retina as an *ex vivo* system (Methods). Confirming the gene regulation activity of the identified CREs, 27.3% and 6.6% of the CREs displayed strong enhancer and silencer activities, respectively (Fig. 4H, Extended Data Fig. 7D, Supplementary Table 12). In addition, we identified transcription factors (TFs) for major classes by integrating snRNA-seq and snATAC-seq data using SCENIC+ ^31^ (Fig. 4I, Supplementary Table 13). A significant portion of the identified TFs have been implicated in specification of individual retinal cell classes, such as *OTX2* and *CRX* for photoreceptor cells, *NR2E3* for rods, *RAX2* for cones, *NEUROD4* for BCs, *ONECUT1* and *ONECUT2* for HCs, *TFAP2A* for ACs, and *NFIB* and *LHX2* for MGs ^32–38^. Many novel TFs were also identified (Supplementary Table 13).

To annotate cell types within classes, we co-embedded snATAC-seq and snRNA-seq data with GLUE and used a logistic regression model to predict the cell type of snATAC-seq cells based on snRNA-seq annotation ^39^ (Methods). For example, 14 BC types corresponding to the 14 cell types by snRNA-seq were identified (Extended Data Fig. 8A-B). Consistently, two snATAC-seq cell clusters were identified for GB/BB and predicted as GB and BB, respectively. Local chromatin accessibility of the marker genes of GB and BB, *UTRN* and *SORCS3* (identified by snRNA-seq, Extended Data Figure 3D-E) also showed high specificity in the corresponding snATAC-seq cell clusters, suggesting gene regulation of *UTRN* and *SORCS3* indeed differ in GB and BB (Extended Data Fig. 8C-D). Similarly, cell types in other heterogenous cell classes, i.e., AC, HC, cone and RGC, as well as non-neuronal cell classes, MG, astrocyte, and microglia cells were distinguished (Extended Data Fig. 8E-H, Supplementary Fig. 7A-B and Supplementary Note).

### The HRCA enables uncovering cell-type-specific gene regulatory circuits

To investigate gene regulatory programs governing individual cell types or groups of types (subclasses) within classes, we performed further SCENIC+ ^31^ analysis (Methods). The identified regulons show high specificity in distinguishing subclasses within the corresponding major cell class, with a maximum regulon specificity score (RSS) ^31^ > 0.8 (Extended Fig. 9A-D, Supplementary Table 13, Supplementary Note). Interestingly, these subclass specific regulons are distinct from the regulons that distinguish their respective major cell class. Some subclasses in different classes share the same TFs. For example, *ISL1* specifically regulates ON-BCs within the BC class and the HC1 type within the HC class. Similarly, *NFIX* is specific for ON-BCs within the BC class and Glycinergic-ACs within the AC class. These findings suggest that cell identity is established through a multiple-layered, hierarchical regulation involving combinations of TFs, with individual TFs playing context-dependent roles.

Using regulons of individual BC types as an example, a set of high-quality regulons that exhibit strong correlation between expression level of TFs and chromatin accessibility of TF target regions across BC types were identified (Pearson correlation rho > 0.70 or <-0.75, Fig. 5A, Supplementary Table 13). It appears that each cell type is under the control of a combination of activators and repressors. For example, *ISL1* and *SMAD9* are activators, while *MEF2C* serves as both activator and repressor for RBC. Importantly, we identified the regulons potentially governing BB and GB, two closely related BC types discerned in this study. Specifically, *ELK4* and *SALL4* appear as activators for BB and GB, *DMBX1* as both activator and repressor for GB, and *PBX1* as both activator and repressor for BB (Fig. 5A, Supplementary Fig. 8A-B). This aligns with the DEG analysis, where PBX1 showed significantly higher expression in BB compared to GB (log2FC=-1.43, *p*-adjust= 5.68 × 10^-55^, Supplementary Table 6).

**Figure 5.**
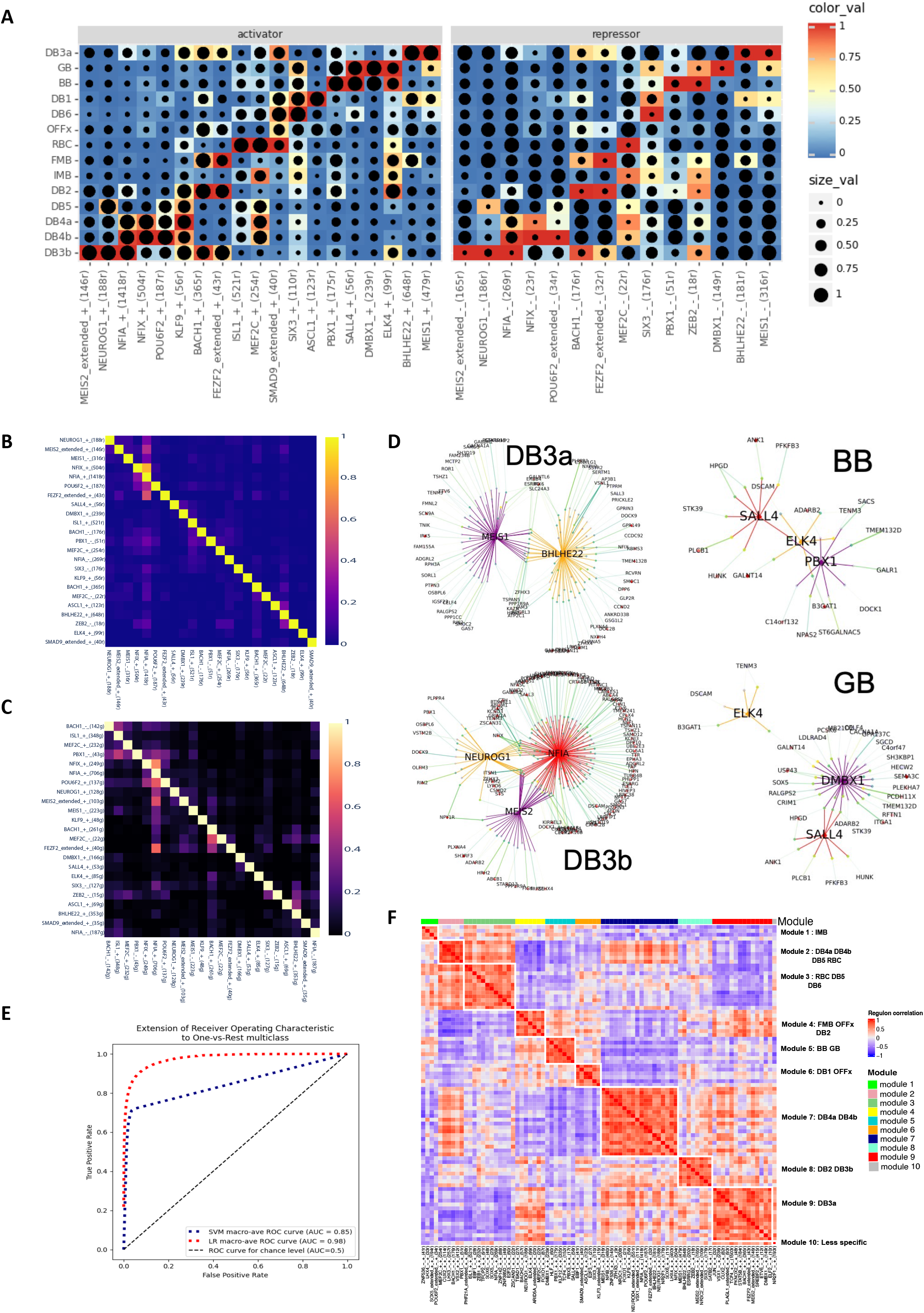
Regulon of the human bipolar cell types. A. Heatmap showing the identified regulons where the gene expression level (color scale) of transcription factors and the enrichment (dot size) of TF motifs in the snATAC peaks are highly correlated. The rows represent BC cell types, and the columns represent the identified regulons. B. Jaccard heatmap showing the intersection of target regions of the identified TFs. Each cell in the heatmap represents the Jaccard index of target regions between a pair of TFs. C. Jaccard heatmap showing the intersection of target genes of the identified TFs. Each cell in the heatmap represents the Jaccard index of target genes between a pair of TFs. D. Network plot showing the regulons and interactions between them in DB3a, DB3b, BB and GB. Each regulon includes the TF, target regions and target genes. E. ROC-AUC of logistic regression model and SVM model to predict BC cell type based on the accessibility of target regions of identified TFs. F. Heatmap showing the correlation in target-regions-based AUC of the identified regulons.

It is worth noting that the cell type regulons show reduced cell type specificity compared to those at the cell class and subclass levels, with a maximum RSS lower than 0.5 (Extended Data Fig. 9E, Supplementary Table 13). Indeed, we observed potential TF cooperativity, exhibited as overlap of the target regions and target genes among these TFs (Fig. 5B-C, Supplementary Note). For example, a subset of *NFIA* target regions and target genes overlap with those of *MEIS2* and *NEUROG1*, while their target regions are highly accessible and their gene expression level are high in DB3b. Interestingly, *NFIA* target regions and target genes also show overlap with those of *NFIX* and *POU6F2*, while the accessibility of their target regions and their gene expression level are high in DB4b. (Figure 5A-C). Thus, as is the case for classes, the same TF can collaborate with different TFs in distinct types. Consistently, regulon network analysis revealed interconnections among these regulons, demonstrated by the mutual or directional regulation among TFs and their regulation of the shared target regions and target genes (Fig. 5D).

To further evaluate the identified TFs, we utilized chromatin accessibility of the target regions of these TFs to predict the cell type via a logistic regression model and a Support Vector Machine (SVM) model (Methods). The logistic regression model achieved a high ROC-AUC value of 0.98 (Figure 5E, Methods), supporting our findings. We also calculated the correlation of regulons based on the regulon activity, which was measured by target region AUC values associated with cell type identities, resulting in 10 regulon modules (Methods, Fig. 5F, Supplementary Table 13). Most of these regulon modules have higher AUC values for specific subsets of BC types, particularly those that are more similar in transcriptome profiles (Extended Data Fig. 9F and Fig. 2D, Supplementary Fig. 8C). In summary, these observations suggest that each cell type is defined by a unique TF combination code, established through precise modulation of both TF expression and the chromatin state of their target regions in each type.

### Differential gene expression associated with age and sex

Differences in retinal functions and disease risks have been associated with individual traits such as age and sex ^40, 41^. We sought molecular correlates of these differences in a set of 135 samples from 57 donors (39 male and 18 female) aged 10 to 91 years (Methods and Supplementary Note), including 24 newly profiled samples from 14 young adult donors (Supplementary Table 14, Extended Data Fig. 10A). We identified 465 to 2,693 genes per cell class with age-dependent expression, utilizing a linear mixed effect model (LMM) (*q*-value < 0.05, Fig. 6A-B and Extended Data Fig. 10B, Supplementary Table 15, Methods). Notably, surges of gene expression changes were observed around the ages of 30, 60, and 80 across major classes, revealed by a sliding window analysis (Fig. 6C, Supplementary Table 16, Methods). Although the dynamic patterns of gene expression changes were similar across classes, many DEGs (on average 37.6% per cell class) were specific to single classes (Fig. 6B, Extended Data Fig. 10C). Gene set enrichment analysis of the age-dependent DEGs pinpointed several pathways activated across cell types (Fig. 6D-E, Supplementary Table 17, FDR < 0.1). They include complement and coagulation cascades, steroid hormone biosynthesis, adaptive immune response, and regulation of calcium ion import (Fig. 6D-E, Supplementary Table 17). Complement pathways have been shown to play important roles in the pathogenesis of age-related macular degeneration (AMD) ^42–47^, and alterations in steroid hormone homeostasis have been linked to glaucoma ^48, 49^. In contrast, the common suppressed pathways included ribosome, cytoplasmic translation, mitochondrial gene expression, and ribonucleoprotein complex assembly, aligning with findings in a fly aging study ^50^ (Fig. 6D, Extended Data Fig. 10D). Suppression of oxidative phosphorylation, protein folding and modification process, ATP metabolic process, and several pathways involved in multiple neurodegeneration diseases were observed in RGC (Fig. 6D-E). These results highlight age-related changes in gene expression that may contribute to age-dependent incidence of major retinal diseases.

**Figure 6.**
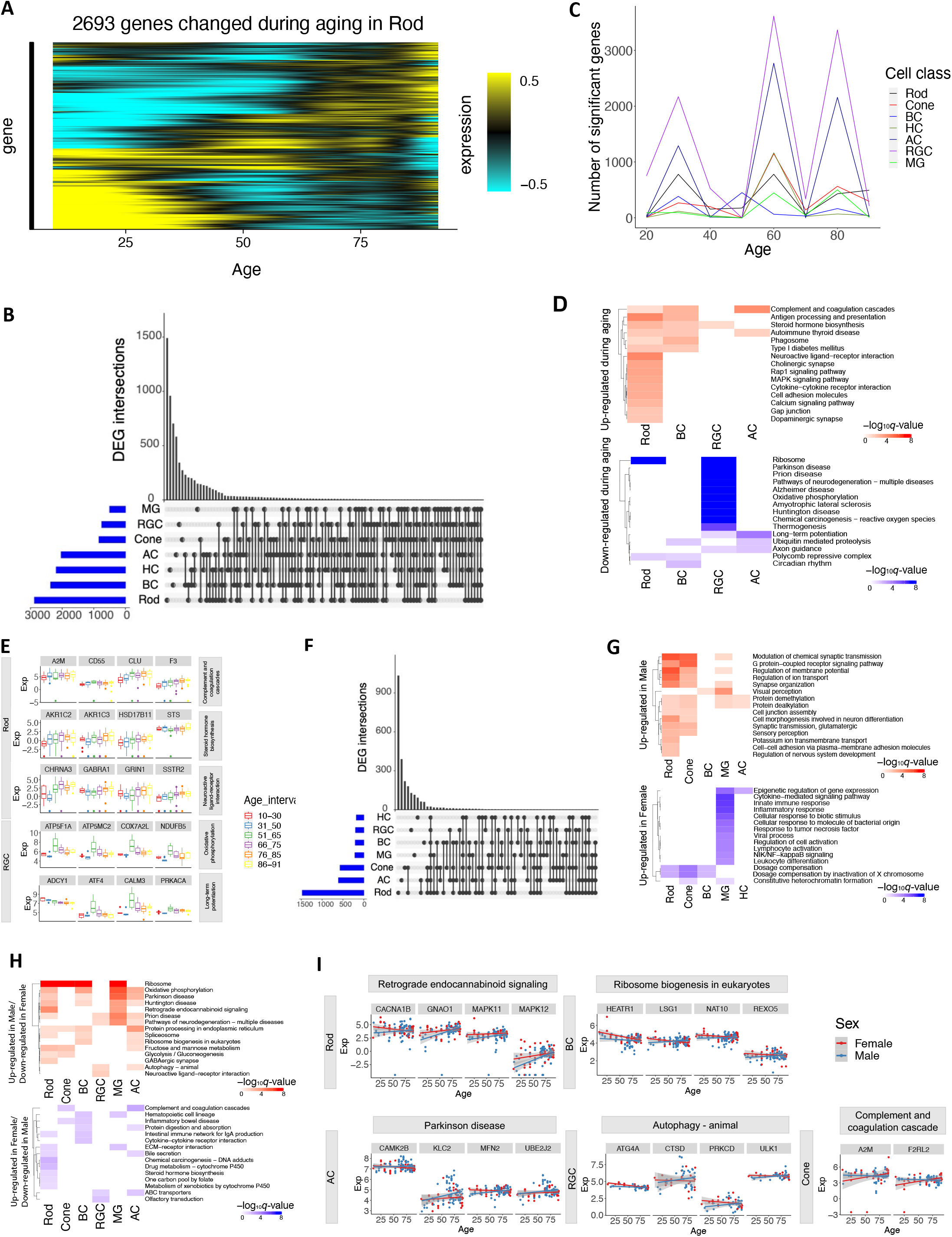
Differential gene expression associated with sex and age. A. Heatmap showing gene expression level of differentially expressed genes (DEGs) during aging in Rod identified with linear mixed effect model (LMM). B. UpSet plot showing the number of cell type specific and common DEGs across major retinal cell classes. C. The number of DEGs identified through sliding window analysis at each age stage. D. The selected KEGG pathways significantly enriched (FDR <0.1) of DEGs during aging identified by LMM across retinal cell classes. E. The examples of DEG during aging involved in the enriched KEGG pathways. F. The number of DEGs between male and female across major retinal cell classes. G. The selected GO terms significantly enriched (FDR < 0.1) of DEGs between male and female across retinal cell classes. H. The selected KEGG pathways significantly enriched (FDR < 0.1) of DEGs with gender dependent aging effect. I. The examples of DEGs with gender dependent aging effect involved in the enriched KEGG pathways.

We also observed transcriptomic differences between males and females across cell classes (Supplementary Table 18, Supplementary Note). The majority (87.7%) of DEGs (*q*-value < 0.05, |log2FC| ≥0.5) were identified on the autosomes while the remaining (12.3%) were on the X or Y chromosomes. Similar to the DEGs associated with aging, many DEGs between males and females (average 53.6% per cell class) are cell class specific (Fig. 6F) and enriched of both cell type specific and shared GO terms (FDR < 0.1, Fig. 6G, Extended Data Fig. 10E, Supplementary Table 17, Supplementary Note). For example, immune-related genes such as those involved in cytokine-mediated signaling pathways, viral processes, and innate immune responses are up-regulated in females specifically in MG (Fig. 6G, Extended Data Fig. 10E). This finding aligns with the sexual dimorphism observed in the mammalian immune system, where females have higher levels of immune responsiveness than males ^51, 52 53, 54^.

Finally, expression of some genes exhibits sex-dependent aging changes driven by sex-age interaction. (Supplementary Table 17 and 19, FDR < 0.1). For examples, genes involved in complement and coagulation cascades, e.g., *A2M* and *F2RL2*, show more significant activation during aging in females compared to males in cones and ACs (Fig. 6H-I). This result aligns with the previous studies suggesting *F2RL2*’s role in progression to advanced macular disease with neovascularization ^55^ and higher prevalence of neovascular age-related macular degeneration in females than males ^56^. Conversely, genes involved in autophagy exhibit more significant up-regulation over aging in males compared to females in RGCs and ACs (e.g., *ATG4A*, *CTSD*, *PRKCD*, *ULK1* in RGC, Fig. 6H-I). Interestingly, autophagy has been found to play a crucial role in glaucoma ^57, 58^, which is more prevalent in males than females ^59, 60^.

### Leveraging the HRCA to study GWAS and eQTL loci

The HRCA provided a unique opportunity to prioritize candidate causal variants, genes, and affected cell types underlying GWAS traits in a multimodal way. To demonstrate this utility, we first identified enriched cell classes associated with GWAS traits based on cell class specific OCRs and gene expression using LDSC ^61^ and MAGMA ^62^, respectively (Fig. 7A, Supplementary Fig. 9A, q-value < 0.05). Consistent results are obtained from both snRNA-seq and snATAC-seq datasets. We observed significant enrichment of age-related macular degeneration (AMD)-associated loci in Retinal Pigment Epithelium (RPE) and Microglia ^63^. Loci linked to the thickness of the outer segment (OST), inner segment (IST), and outer nuclear layer (ONL) exhibited enrichment in rods, cones, and MGs ^64^. Loci associated with traits related to open-angle glaucoma were enriched in MGs and Astrocytes ^65–67^. Refractive error and myopia loci showed enrichment across most retinal cell classes ^68^. As a negative control, bone mineral density loci did not display enrichment in any of the retinal cell classes^69^.

**Figure 7.**
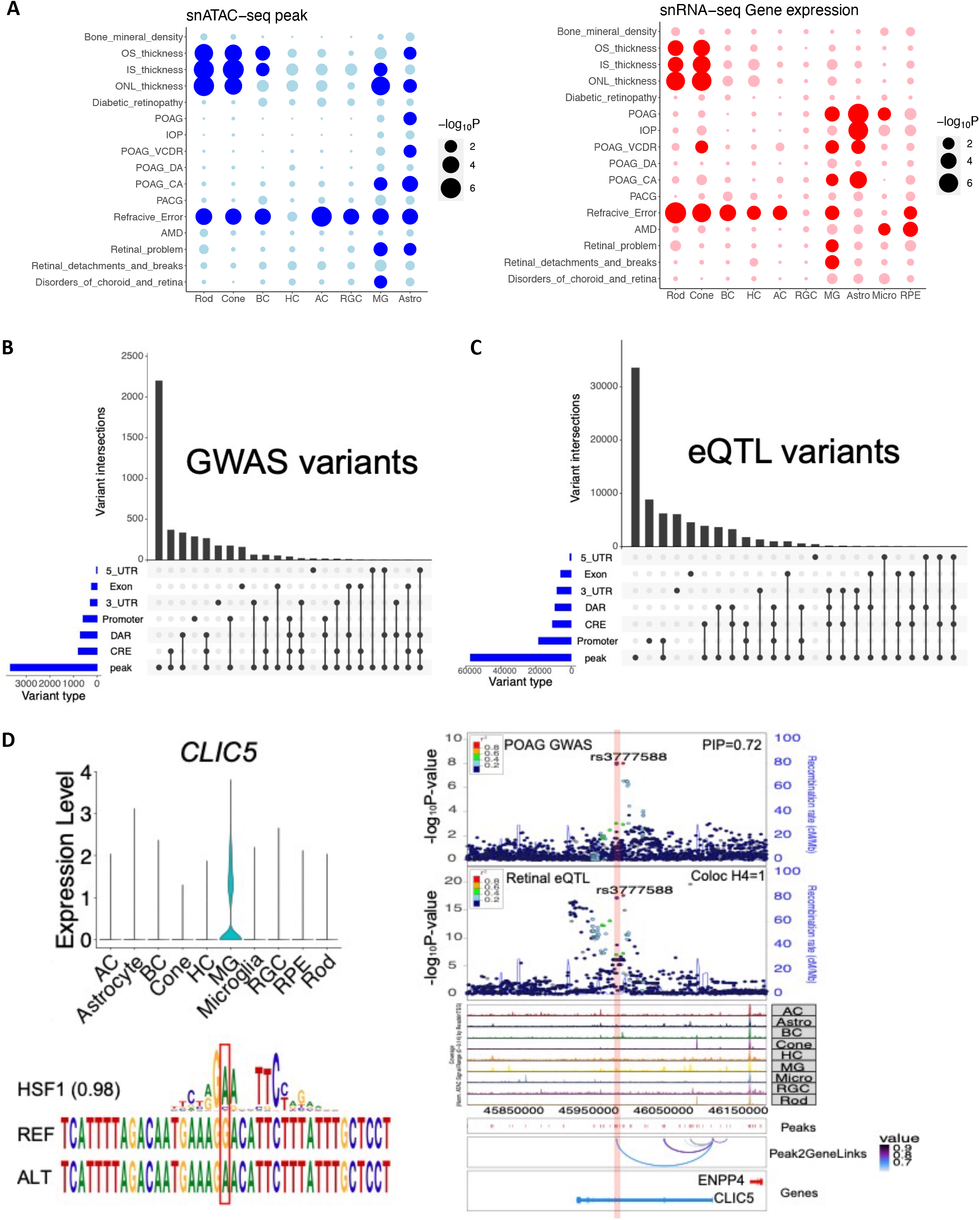
Leveraging multi-omics data to study GWAS and eQTL loci. A. Cell class enrichment of GWAS loci based on chromatin accessibility with LDSC (left) and gene expression with MAGMA (right). Rows represent enriched GWAS traits, and columns represent retinal cell classes. The highlight dot indicates the enrichment q-value < 0.05. B. Categorization of fine-mapped GWAS variants located in various genomic regions. Categories include peak (i.e., open chromatin regions), linked cis-regulatory elements (CREs), differentially accessible regions (DARs), promoter, exon, 5_UTR and 3_UTR of gene annotation. C. Categorization of fine-mapped eQTL variants located in various genomic regions. D. Visualization of fine-mapped loci in *CLIC5* region.

To further identify candidate causal variants, target genes, and affected cell types for GWAS loci, we performed fine-mapping of GWAS loci associated with seven retinal GWAS traits: OST ^64^, IST ^64^, ONL ^64^, POAG ^65^, AMD ^68^, refraction error/myopia ^68^, and diabetic retinopathy ^70^. Based on summary statistics and linkage disequilibrium of genome-wide variants analyzed in previous GWAS studies, we identified 18,959 variants that fell within the 95% credible sets of these GWAS loci (Fig. 7B and Supplementary Table 20). Notably, a substantial proportion (19.4%, n=3,673) of the variants were found within OCRs (i.e., snATAC-seq peaks). Additionally, small subsets of variants were mapped in regions where target genes could be inferred: 4.2% (796) were within linked CREs, 3.1% (592) within promoter regions, and 2.9% (553) within exonic, 3’ UTR and/or 5’ UTR regions, resulting in 1,784 variants linked to 691 potential target genes (Table 1). By cross-referencing these GWAS variant-gene pairs with eQTL-eGene pairs identified in bulk retina tissue, we found that 130 GWAS genes were eGenes of the GWAS variants, reinforcing the validity of our findings. Furthermore, a significant proportion of the identified target genes are marker genes of disease relevant cell classes, known genes linked to complex diseases or inherited retinal diseases (Table 1). Specifically, we uncovered well-known AMD related genes such as *APOE*, *C2*, and *C3*. In the case of POAG, our findings included *EFEMP1* ^71^, which has been linked to familial juvenile-onset open-angle glaucoma, as well as *TMCO1* and *SIX6*, known to be associated with POAG ^72^. For diabetic retinopathy, *ABCF1* was identified as a regulator of RPE cell phagocytosis ^73^ and as one of the proteomic biomarkers of retinal inflammation in diabetic retinopathy ^74^. For target genes linked to retinal layer thickness, we pinpointed *ATOH7*, *PAX6*, *VSX2*, and *RAX*, all of which have been implicated in retinogenesis ^75, 76^. Additionally, we identified genes like *MKKS*, *FSCN2*, *PDE6G*, *PRPH2*, *RDH5*, *RHO*, *SAG*, *RP1L1*, and *RLBP1*, known to be associated with inherited retinal diseases. Similarly, we fine-mapped retinal eQTLs using a comparable method (Fig. 7C). A significant portion of eQTL variants was also found within OCRs, while eQTLs exhibited greater enrichment in promoter regions than GWAS variants (two-sided binomial test, *p* = 4.94 × 10^-324^, Supplementary Table 21). Moreover, these fine-mapped variants provided candidates to study regulatory mechanism of GWAS loci (Supplementary Note). As an example, one POAG variant (rs3777588) was fine-mapped (posterior inclusion probability [PIP]=0.72) to a LCRE of *CLIC5* (Fig. 7D), a region specifically open in MG. Consistently, *CLIC5* is highly expressed in MG among retinal cell classes. Furthermore, the GWAS signal was colocalized with retinal eQTL signal of *CLIC5* through this variant (H4=1.00 and Methods). Notably, this variant was also predicted to strength the binding of the transcription factor *HSF1*.

**Table 1.**
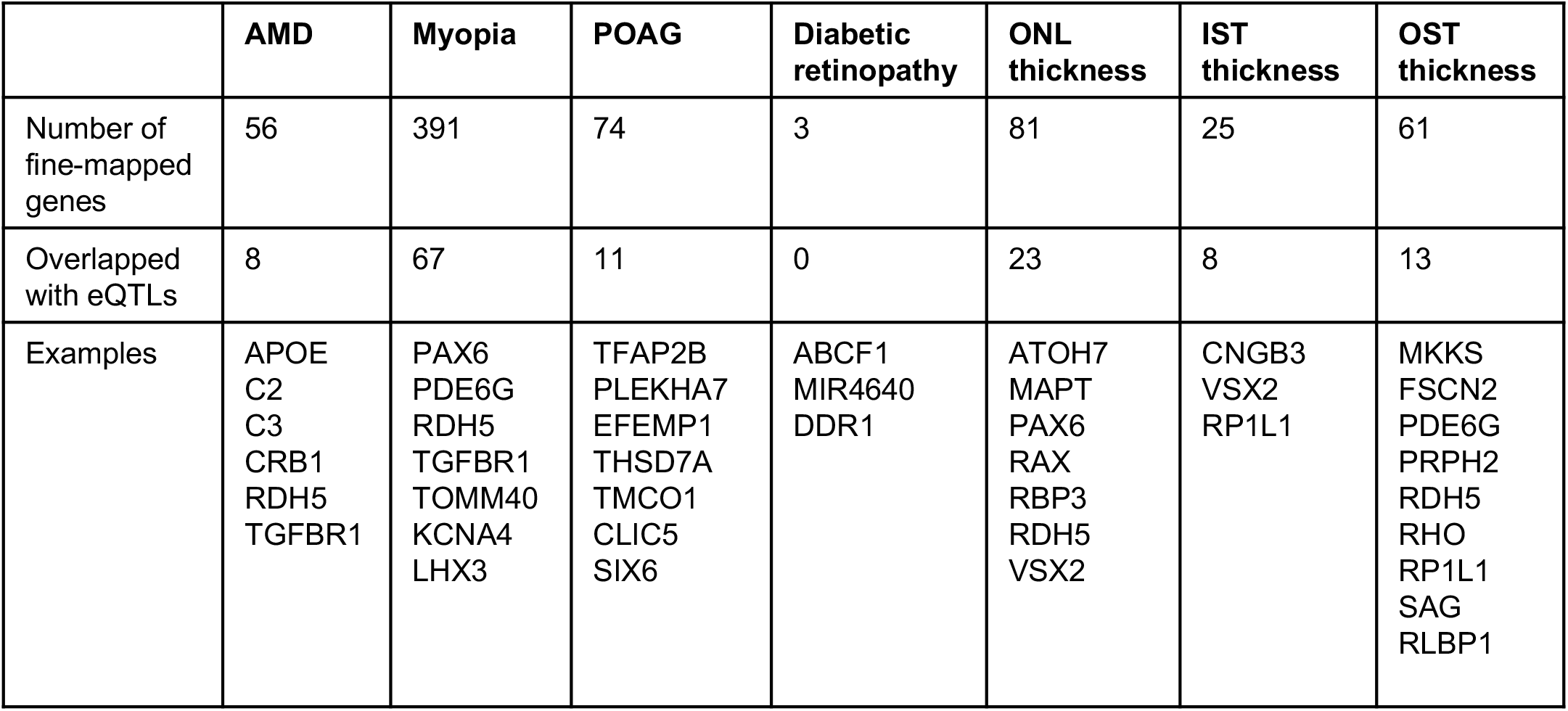
Summary of fine-mapped GWAS loci associated with the seven GWAS traits.

## Discussion

In this study, we introduced HRCA version 1, an integrated multi-omics single-cell atlas of the human retina, which marks the first multi-omics reference atlas in the HCA framework ^1, 2^. The HRCA provides a comprehensive view of the transcriptomic and chromatin profiles of retinal cells, comprising data from more than 2 million sn-/sc-RNA-seq cells and over 370,000 snATAC-seq cells. Our cross-donor and cross-lab atlas provides a model for future HCA atlases. The HRCA is accessible for the community through numerous interactive platforms, including CELLxGENE ^77^, UCSC Cell Browser ^78^, and Single Cell Portal ^79^, and can therefore serve as a common reference for advancing research on human eye health and diseases.

Given the large number of cells profiled, coupled with targeted cell enrichment, the HRCA is nearly saturated for retinal cell types. The integrated analysis of over 2 million single cell/nuclei, including 1.4 million unpublished data points, revealed over 110 cell types in the human retina, nearly doubling the number reported in previous studies ^7^. For example, the HRCA separates two rare and closely related BC types, GB and BB, which co-clustered in previous analyses ^7, 9, 15^. Cross-species comparisons among humans, macaques, and mice augment those reported previously ^7, 9, 15^, especially with additional species ^8^, improving cell type annotation and providing guidance for translational studies in rodents of human vision disorders. Further annotation of this atlas by experts from the community will be used to update the HRCA.

The HRCA also provides a comprehensive gene regulatory landscape of the human retina at single-cell resolution, uncovering over 670K open chromatin regions, and revealing potential CREs in individual cell type contexts. These results enable the identification of GRNs defining cellular identities at the class, subclass, and cell type levels, revealing a multiple-layered, hierarchical regulation principle involving combinations of TFs. Hundreds of CREs linked to IRD genes were validated through a high-throughput functional assay in an *ex vivo* mouse model system. However, a high proportion of inactive sequences were observed in validation, which may result from a combination of limited experimental sensitivity, divergent human-mouse CRE activity, and inactive or false enhancers. Silencers in scrambled CRE sequences could result from retained motif content but low motif diversity ^80^.

Intriguingly, the HRCA also enabled the discovery of dynamic patterns of transcriptome during aging, where DEG surging patterns were consistent across cell types, but the individual genes were mostly differentially expressed in only one or two cell classes. A subset of aging-related DEGs is overlapped with GWAS genes of aging-related diseases, e.g., *C3* in Rod and *VEGFA* In Cone, and aging-related biological pathways include some known to be associated with age-related diseases, such as age-related macular degeneration. Similarly, we detected cell type specific transcriptomic and pathway difference between sexes beyond sex chromosomes, including immune response-related dimorphisms in autosomal genes expression. Interestingly, certain genes show sex-specific aging patterns, which may shed light on gender differences in certain age-related diseases.

Finally, the HRCA facilitated a comprehensive functional annotation of disease-related variants, and exploration of the regulatory mechanisms of causal variants. By combining HRCA with fine-mapping, we identified potential causal variants, target genes, and the acting cell types associated with GWAS and eQTL loci, providing testable hypotheses about the action mode of GWAS variants. Additionally, we offer utilities designed to automate the annotation of cell types for new samples using scArches ^81^ (Supplementary Fig. 10 and Supplementary Note). In summary, the HRCA represents a comprehensive reference of the human retina and facilitates future analysis across cell types, individuals, and diseases for the human eye.

## Methods

### Human retina sample collection

Tissues not described in previous publications were obtained from 28 individuals within 6 hours post-mortem from the Utah Lions Eye Bank. Detailed donor information can be found in Supplementary Table 2. The procedure for dissecting the eyes followed the established protocol ^82^. Macular samples were collected using disposable biopsy punches measuring 6 mm in diameter. Subsequently, the retinal tissues were flash-frozen in liquid nitrogen and stored at −80 °C until nuclei isolation. Only healthy donors with no recorded medical history of retinal diseases were included in this study. Post-mortem phenotyping using OCT was conducted to confirm the absence of disease phenotypes, such as drusen or atrophy, as described in the previous study ^7^. Institutional approval for the patient tissue donation consent was obtained from the University of Utah, adhering to the tenets of the Declaration of Helsinki. Each tissue was de-identified in accordance with HIPAA Privacy Rules.

### Nuclei isolation and sorting

The frozen retinal tissues were resuspended and triturated in a freshly prepared, pre-chilled RNase-free lysis buffer (10 mM Tris-HCl, 10 mM NaCl, 3 mM MgCl2, 0.02% NP40) with a Wheaton™ Dounce Tissue Grinder to obtain nuclei. To enrich the retinal ganglion cell nuclei, isolated macular retinal nuclei were stained with a mouse anti-NeuN monoclonal antibody (1:5000, Alexa Flour 488 Conjugate MAB377X, Millipore, Billerica, Massachusetts, United States) in staining buffer (1% BSA in PBS, 0.2U/μl RNAse inhibitor) for 30 minutes at 4°C. After centrifugation at 500g 4°C for 5 minutes, nuclei were resuspended in staining buffer and filtered with a 40μm Flowmi Cell Strainer. DAPI (4′,6-diamidino-2-phenylindole, 10 μg/ml) was added before fluorescent cytometry sorting.

The stained nuclei were sorted with a BD (Becton Dickinson, San Jose, CA) Aria II flow sorter (70μm nozzle). Gating was performed based on flow analysis of events and strengths of DAPI (450-nm/40-nm-band pass barrier filter) and FITC (530-nm/30-nm-band pass filter) signals. The sorting rate was 50 events per second based on side scatter (threshold value > 200). The nuclei group with strongest 5% FITC signal was collected for RGC enrichment, specifically, while all DAPI-positive nuclei were collected for general retinal nuclei study.

For single nuclei ATAC profiling, nuclei were isolated in fresh-made pre-chilled lysis buffer (10 mM Tris-HCl, 10 mM NaCl, 3 mM MgCl2, 0.02% NP40, 1%BSA) with a Wheaton™ Dounce Tissue Grinder until no tissue pieces were visible. After being washed at 500g, 4C for 5min twice in a pre-coated 5ml round bottom Falcon tube (wash buffer: 10 mM Tris-HCl, 10 mM NaCl, 3 mM MgCl2, 1%BSA; coating buffer: 10 mM Tris-HCl, 10 mM NaCl, 3 mM MgCl2, 4%BSA; Falcon tube Cat. NO. 352054), the nuclei were resuspended in 1X diluted nuclei buffer (10X PN-2000153, PN-2000207) with a final concentration of 3000-5000 nuclei/ul.

### Single-nuclei RNA and ATAC sequencing

All single-nuclei RNA and single-nuclei ATAC sequencing was conducted at the Single Cell Genomics Core at Baylor College of Medicine in this study. The library preparation and sequencing of single-nuclei cDNA were carried out following the manufacturer’s protocols (https://www.10xgenomics.com). To obtain single cell GEMS (Gel Beads-In-Emulsions) for the reaction, single-nuclei suspension was loaded onto a Chromium controller. The library for single nuclei RNA-seq was prepared with the Chromium Next GEM Single Cell 3’ Kit v3.1 (10x Genomics), while the library of single nuclei ATAC-seq was prepared with the Chromium Next GEM Single Cell ATAC Library and Gel Bead Kit v1.1 (10x Genomics). The constructed libraries were subsequently sequenced on an Illumina Novaseq 6000 (https://www.illumina.com).

### Data preprocessing of unpublished and public datasets

Raw sequencing reads were first downloaded for all the curated public datasets. Along with unpublished generated datasets, data samples were processed using the same versions of software and databases by a quality control pipeline (https://github.com/lijinbio/cellqc). Raw sequencing reads were first analyzed using 10x Genomics Cell Ranger (version 7.0.1) ^83^ utilizing the hg38 genome reference obtained from 10x Genomics (https://cf.10xgenomics.com/supp/cell-exp/refdata-gex-GRCh38-2020-A.tar.gz). The resulting feature count matrices were retained for downstream quality control. Cell Ranger implemented EmptyDrops to filter empty droplets in experiments based on significant deviations from a background model of low-count cells ^84^. To further eliminate potential empty droplets from the filtered feature count matrices by Cell Ranger, dropkick was utilized to construct dataset-specific training labels by applying a logistic regression for real cells, with a threshold based on the total number of transcript counts in cells ^85^. The real cells retained were those identified by both EmptyDrops and dropkick, and they were preserved for downstream analysis. To correct for the background transcript measurements derived from ambient RNAs that are not endogenous to cells, SoupX was used to estimate a global contamination fraction across cells and to correct gene expression profiles by subtracting the contaminations ^86^. To exclude potential multiplets, DoubletFinder simulated artificial doublets and ranked real cells based on the proportion of artificial neighbors ^87^. Cells predicted to be multiplets with high proportions of artificial neighbors were ruled out. Following cell filtering criteria of ≥ 300 features, ≥ 500 transcript counts, and ≤10% (or ≤ 5% for snRNA) of reads mapped to mitochondrial genes, the retained cells constituted the clean cells for downstream analysis.

To annotate major retinal cell classes, a pre-trained multi-class classifier was applied using scPred to predict a type for each cell ^88^. The training data was constructed in-house by collecting cells with ten major annotated cell classes, including amacrine cells (AC), bipolar cells (BC), horizontal cells (HC), retinal ganglion cells (RGC), retinal pigment epithelium (RPE), astrocytes, muller glia (MG), microglia, rods and cones. Raw gene expression counts were initially log-normalized and scaled using Seurat. The scaled matrix was decomposed through principal component analysis. The principal component embeddings were the features utilized for training binary-SVM classifiers (one-versus-all) for cell types. During prediction, the raw counts matrix of test data was also initially log-normalized and scaled using Seurat ^89, 90^. The scaled data were then projected into the principal component coordinate basis established by the training data. The projected principal components served as features for prediction against the trained classifiers. Positive cell types were predicted based on classification probabilities ≥ 0.9, and doublets were identified if cells were classified into multiple types.

### Integration benchmarking of single cell and single nuclei RNA-seq sequencing

An integration benchmarking of retina datasets was conducted based on previous work, such as scIB ^17^ and the the Human Lung Cell Atlas v1 ^3^. Briefly, cells from each donor and sample were independently annotated using one of nine major class cell types using scPred, and then these datasets were concatenated as a single input object, with annotations for batches, cell types, and technologies (sc or sn). We tested two levels of feature selection, 1,000 and 3,000 highly variable genes (HVGs), we only tested raw counts without rescaling based on previous insights.

To allow batch correction comparisons between single-cell and single-nuclei datasets, we performed three integration pipelines: one with only single-cell RNA-seq datasets (sc), one with only single-nuclei RNA-seq datasets (sn), and one with both dataset types combined (sn+sc). This allowed measuring the integration quality of cells based on matched cells from the combined technologies, with respect to each technology alone.

Due to scaling limitations while running methods for the largest single-cell datasets, (more than two million cells), we limited our tests to Python methods with a scalable implementation. Empirically, methods were discarded if output was not generated in 48 h as a single task, with 150GB of memory, 4 CPUs, and one GPU if required. Based on these criteria, we were able to generate batch-corrected objects for 7 methods using 1,000 HVGs, including scANVI, scVI, scGen, scanorama, BBKNN, Harmony (harmonpy), and combat. When using 3,000 HVGs and sn-datasets, scanorama and BBKNN were discarded. When benchmarking sn+sc datasets, scGen and Combat were discarded due to running times.

The calculation of some metrics requires a non-linear time with respect to the number of cells, and this makes their computing expensive for the largest datasets. As an improvement during the metrics calculation step, we incorporated into our pipeline a metrics approach to allow fixed subsamples of the full object, with custom percentage sub-samples set up as 3, 5, and 6 percent. This allows measuring integration quality with a sample representative of the full object, and in a shorter computational time, while recovering best methods with a lower computational effort.

### Integration of single cell and single nuclei transcriptome data

From the benchmark results, scVI^24^ outperformed all the label-agnostic methods in our benchmark results. Therefore, scVI was selected for integrating the transcriptome data. On the entire 2 million cells, the major cell classes are well integrated, but the subclass clusters within the major classes are mixed. For example, many clusters of the AC class are intermixed with clusters of the BC class (Extended Data Fig. 1C). We compared the cell distribution of snRNA-seq and scRNA-seq and found that many cell clusters overlap between the two technologies, while a few do not (Extended Data Fig. 1D). Therefore, separate integrations for single-nuclei and single-cell samples were conducted to account for the differences in dissociation technologies. For integrating data specific to BC, AC, and RGC types, only subsets of cell type-specific cells for subclass integration were retained. To capture the nuanced similarities between cell clusters, the top 10,000 highly variable genes was calculated using the “sampleID” as the batch key with the Scanpy Python package ^91^. The “sampleID” was also used as the batch variable in the scVI modeling. In scVI, two hidden layers for encoder and decoder neural networks and a 30-dimensional latent space were calculated to represent cells after removing sample batches. The number of epochs was adjusted based on the total number of cells in the subclass integration and a minimum of 20 epochs was used for the variational autoencoder training. The trained latent representation was used to measure the distance among cells. These distances were used to calculate the cell clustering using the Leiden algorithm ^92^. To facilitate the inspection of integrated cell clusters, 2D visualization was generated using UMAP ^93^. To determine the optimal resolution for the Leiden clustering, a range of resolution values were evaluated and manually examined by the resulting cell clusters using a UMAP plot. To assess and mitigate potential over-clustering, the self-projection accuracy of the clustering was computed using the SCCAF Python package ^94^. Furthermore, a two-level clustering method was used to capture the cellular diversities of BC, AC, and RGC when performing subclass clustering. Various resolutions were tested for clustering, and the first-level resolution was selected to achieve initial clustering without over-clustering, as confirmed by UMAP visualization. In the second-level clustering, various resolutions were also tested to refine any under-clustering and achieve optimal clustering without over-clustering on UMAP. Ultimately, the two-level clustering approach determined the number of clusters in the atlases.

### Comparison between snRNA-seq and scRNA-seq

To evaluate the differences between snRNA-seq and scRNA-seq, the cell proportions of major cell classes were computed in each sample using both technologies. The samples were categorized into fovea, macular, and periphery tissue regions for both approaches. To address any potential cell proportion bias arising from experimental enrichment in a subset of snRNA-seq samples, only samples without enrichment were included. Subsequently, bar plots were generated to compare the cell proportions of major classes across tissue regions for the two technologies.

To examine the cell type similarities of major classes between the two technologies, raw counts of the complete cells were first aggregated into pseudo-bulk for each major class across samples. The resulting pseudo-bulk measurement has three metadata columns: the “sampleID,” which represents unique sample IDs in the atlas; “dataset,” indicating whether the sample is from “snRNA” or “scRNA” technologies; and the “majorclass,” which denotes the annotated major class cell types. Utilizing the pseudo-bulk count matrix, cell type similarities were calculated using the MetaNeighbor R package ^95^. Specifically, highly variable genes were detected using the “variableGenes()” function with “dataset” as the source of samples, and the mean AUROC matrix was calculated for “dataset” and “majorclass” using the “MetaNeighborUS()” function with the calculated variable genes.

To identify differentially expressed genes in two technologies, the DESeq2 R package^96^ was applied to the aggregated pseudo-bulk count matrix. To account for major class cell type information during the statistical test, the design formula used “∼ majorclass + dataset”. The Wald test was employed to calculate *p*-values of gene expression differences between the two technologies. The contrast used in the “results()” function was “contrast=c(‘dataset’, ‘snRNA’, ‘scRNA’)” to derive differentially expressed genes after regressing out major classes by “majorclass”. To enhance the statistical power, genes with average expressions less than 10 among pseudo-bulk samples were excluded from the analysis. For calculating adjusted q-values from the p-values, we employed the Benjamini-Hochberg procedure ^97^. Subsequently, differentially expressed genes were identified under |log2 fold change|>1 and *q*-value<0.05. Enriched Gene Ontology (GO) terms were identified using the “enrichGO()” function of the clusterProfiler R package ^98^ on the differentially expressed genes. To investigate gene expression changes among major class cell types between the two technologies, the count matrix was subsetted per major class and subjected to differential gene expression analysis using the design formula “∼ dataset” in a similar manner. To explore the shared differentially expressed genes across major classes, an UpSetR image was produced using the “upset()” function from the UpSetR R package ^99^.

### Cross-species analysis

To conduct cross-species analysis, the SATURN algorithm^26^ was utilized to compare human, mouse, and macaque cell clusters and cell types. The human cell clusters were identified from clean cells, while the mouse reference was generated from an integrated analysis of collected mouse samples available at the data portal of Baylor College of Medicine (https://mouseatlas.research.bcm.edu/). Raw single cell measurements and cell labeling for the macaque reference were obtained from the GEO repository (accession GSE118546)^9^. To ensure accurate alignment of cell clusters, we randomly sampled up to 2,000 cells per cell cluster and cell type. Protein embeddings for human, mouse and macaque are retrieved from the respective SATURN repositories. To capture nuanced similarities among cell clusters, SATURN feature aggregation employs a set of 5,000 macrogenes. Additionally, during pre-training, “sampleID”s are utilized as non-species batch keys to effectively reduce batch effects caused by samples. The trained 256-dimensional latent representations were utilized to compute cell dissimilarities and generate UMAP for visualizations.

### Differential gene expression analysis for bipolar cells

The DESeq2 R package^96^ was utilized to identify genes that were highly expressed in specific cell types, e.g., GB and BB cell types. First, a pseudo-bulk measurement was calculated by summing the gene expressions of single cells within each cell type for each sample, excluding samples with less than 2,000 cells. The pseudo-bulk datasets were then used in a paired test, incorporating sample information in the design formula “∼ sampleID + celltype”. Lowly expressed genes with an average expression less than 10 were filtered out to improve computation speed and statistical power. A Wald test was used to calculate *p*-values for differential testing, comparing gene expression changes between BB and GB by contrasting the “celltype” factor using the DESeq2 package’s “results()” function. The adjusted *q*-value was calculated from *p-*values using the Benjamini-Hochberg procedure ^97^. The EnhancedVolcano R package^100^ was used to visualize the distribution of log2 fold change and *q*-values. Differentially expressed genes were identified based on criteria of |log2 fold change|>1 and *q*-value<0.05. Enriched Gene Ontology (GO) terms were identified using the “enrichGO()” function of the clusterProfiler R package^98^ on the changed genes.

To identify the top-ranked genes in GB and BB between the snRNA-seq and scRNA-seq datasets, we normalized and transformed raw count matrices from the two technologies using the “normalized_total()”and “log1p()” functions within the Scanpy Python package ^91^. To expedite the computation, 10,000 highly variable genes were calculated using the “seurat” flavor with the batch key set as the “sampleID”. Subsequently, the highly variable genes were tested for top-ranked genes via the Wilcoxon test. Top-ranked genes were identified by *q-* value < 0.05. To visualize the overlapped genes, a venn diagram was generated using the “venn.diagram()” function from the VennDiagram ^101^ R package. Fisher’s exact test was used to calculate the significance of the overlap of top ranked genes between GB and BB in snRNA-seq and scRNA-seq, with 10,000 genes as the background for gene expression.

To evaluate the cell type similarities between DB4a, DB4b, and DB5 in humans and their corresponding mapped cell types in mice and macaques, gene symbols of the raw count matrices of mouse and macaque data were converted into human orthologs using the MGI ^102^ and HGNC ^103^ databases. Utilizing human gene symbols and orthologs, cell type similarities were computed in a manner similar to the comparison of cell types between snRNA-seq and scRNA-seq datasets utilizing the MetaNeighbor R package ^95^.

### Marker identification by binary classification analysis

To identify novel markers for BC, AC, and RGC types, a binary classification approach was applied to detect 2- or 3-marker combinations for each type^13^. To mitigate classification bias resulting from unbalanced cell type abundances, up to 2000 cells were randomly sampled for BC types, and up to 500 cells were sampled for AC and RGC clusters. First, the raw counts were normalized, and the top 50 ranked genes were calculated for each cell type using the Scanpy package ^91^. Support vector classifiers were then trained by considering combinations of the top-ranked genes for each cell type. The “SVC()” function with “kernel=rbf” was employed from the scikit-learn Python package ^104^. Combinations of markers were ranked based on several classification metrics, including precision, recall, F1 score, and AUROC.

### Annotation of snATAC-seq cells and co-embedding of snATAC-seq and snRNA-seq cells

To annotate cell types for snATAC-seq, the low-quality cells and doublets were first filtered out, and the retained cells were clustered with ArchR ^105^ (minTSS=4, minFrags=1000, filterRatio=1). By integrating with snRNA-seq data, six major neuron cell classes and a mixed non-neuron cell class were identified through ArchR. Then peaks were called by MACS2 ^106^ through ArchR and cell by peak fragment count matrices were generated for each of the major cell classes and across major cell classes via Seurat ^89^ and Signac ^107^. The co-embedding of snRNA-seq and snATAC-seq was performed with the GLUE algorithm^39^. Specifically, to integrate major cell class annotation, all snATAC-seq cells were co-embedded with the down-sampled snRNA-seq cells by scGlue under the supervised mode, since major cell classes from both snATAC-seq and snRNA-seq were already annotated. However, to identify cell types per major class, the snATAC-seq cells were co-embedded with the snRNA-seq cells for a major class by scGlue under the unsupervised mode. A logistic regression model and an SVM model were then trained using the GLUE embedding and annotation of snRNA-seq cells to predict the cell types of snATAC-seq cells using the scikit-learn python package. The ROC-AUC of the logistic regression model was consistently higher than that of SVM model, so the logistic regression model was used to annotate snATAC-seq cells. The peaks were called by MACS2 through ArchR for snATAC-seq cell classes and types. Differentially accessible regions (DARs) and linked CREs were identified across cell classes and types using ArchR. The linked CREs were the union set of peak-gene pairs identified through the correlation of accessibility between snATAC-seq peaks (-/+ 250kb surrounding TSS) and promoters (co-accessibility), as well as the correlation between gene expression and the accessibility of snATAC-seq peaks (-/+ 250kb surrounding TSS).

### Identification of regulon of retinal cell types

Regulons were identified for each of major cell classes, subclasses, and cell types respectively utilizing SCENIC+ ^31^. Since SCENIC+ is memory-demanding, up to 1,000, 2,000, or 4,000 cells per cell type (depending on specific cell class/subclass/type) were down-sampled for snATAC-seq cells and snRNA-seq cells respectively. The down-sampled cell by gene matrices and cell by peak matrices were then submitted to SCENIC+. Transcription factors (TF), target regions of TFs, and target genes of TFs were also identified across cell types. The transcription factors that showed significant correlation between gene expression and chromatin accessibility of the target regions across cell types were further selected as candidate TFs. From these TFs, eRegulon Specificity Score (RSS) was also computed for the TFs that were identified as activators in the corresponding cell type. Furthermore, the TFs that displayed a significant correlation between the accessibility of target regions and the expression level of target genes were identified. Subsequently, TF modules displaying a significant correlation in the region-based AUC between TFs were identified.

### Massively parallel reporter assays

We developed a MPRA library, which contains the sequences of 1,820 CRE candidates linked to inherited retinal disease genes identified in the rod cells, along with 20 control cis-regulatory elements (CREs) with a variety of activity that have been previously validated ^80^, and negative controls (i.e., 300 scrambled sequences, and a basal promoter without CRE). Each CRE or control sequence was labeled with three unique barcodes, and 25 barcodes were assigned to the basal promoter. Oligonucleotides (oligos) were synthesized as follows: 5’ priming sequence /EcoRI site/Library sequence (224-bp)/SpeI site/C/SphI site/Barcode sequence (9-bp)/NotI site/3’ priming sequence. These oligomers were ordered from TWIST BIOSCIENCE (South San Francisco, CA) and cloned upstream of a photoreceptor-specific Crx promoter, which drives the expression of a DsRed reporter gene. The resulting plasmid library was then electroporated into three retinal explants of C57BL/6J mice at postnatal day 0 (P0) in four replicates. On Day 8, DNA and RNA were extracted from the cultured explants and next-generation sequencing was conducted. The activity of each CRE was calculated based on the ratio of RNA/DNA read counts and was normalized to the activity of the basal Crx promoter. The bioinformatics analysis of the MPRA result followed the previously published pipeline^80^.

### Differential gene expression analysis during aging and between genders

We conducted two types of differential gene expression analysis during aging. First, for each cell class, raw read counts were aggregated per gene per sample. Only the samples containing at least 100 cells in the corresponding cell class were considered. Additionally, the samples that had < 0.75 correlation in read counts with > 65% of samples were considered as outliers and were not included in subsequent analysis. Genes with low expression in the corresponding cell classes were filtered out, resulting in about 18,003 genes retained per cell class for further analysis. Based on the filtered genes and samples, the genes significantly correlated with aging and different between sexes were identified using a mixed linear effect model via edgeR ^108^ and variancePartition ^109^ R packages. The formula we applied were: ∼ age + sex + race + tissue + seq+ (1|batch) for age and sex effect, and ∼ age + sex + race + tissue + seq+ (age:sex) + (1|batch) for the interaction between age and sex. Log2 fold change and *p*-values were extracted for all genes for the covariate of interest, i.e., age, sex, and interaction between age and sex. In addition, a sliding window analysis was conducted over aging, and DEGs between two adjacent time windows were identified per cell class utilizing the DEswan R package ^110^. The read counts of the filtered genes were normalized based on the library size of each sample per cell class via the edgeR R package. The sliding window analysis was conducted over aging, considering batch and sex as covariates at the age: 20, 30, 40, 50, 60, 70, 80, and 90, with the bucket size = 20 years. In all time windows (10-year interval) except three windows in RGC, there are more than three samples per cell class, ensuring statistical robustness. Enriched pathways and GO terms were identified through gene set enrichment analysis of the differentially expressed genes utilizing the clusterProfiler R package ^98^. The significance cutoff for enriched gene sets was set at FDR < 0.1.

### Cell type enrichment underlying GWAS locus

Cell class enrichment underlying GWAS loci was identified based on both chromatin accessibility and gene expression. For chromatin accessibility, the heritability of GWAS traits were partitioned into cell class specific snATAC-seq peaks using stratified LD score regression via LDSC ^61^. Initially, GWAS SNPs that overlapped with HapMap3 SNPs were annotated based on whether they were in OCRs in each cell class. Subsequently, LD-scores of these SNPs within 1 cM windows were calculated based on the 1000 Genome data. The LD-scores of these SNPs were integrated with those from the baseline model, which included non-cell type specific annotation (downloaded from https://alkesgroup.broadinstitute.org/LDSCORE/). Finally, the heritability in the annotated genomic regions was estimated and compared with the baseline model to determine if regions in each cell class were enriched with the heritability of the corresponding GWAS trait. For gene expression, the linear positive correlation between cell class specificity of gene expression and gene-level genetic association with GWAS studies were assessed by using the MAGMA.Celltype R package ^62^. GWAS summary statistics were formatted with the “MungeSumstats” R package ^111^ based on SNPs in - 35kb/+10kb of each gene and 1000 genome “eur” population. snRNA-seq expression data was formatted with the “EWCE” R package ^112^. Linear enrichment was detected using the MAGMA.Celltype R package. To correct for multiple testing, the Benjamini-Hochberg method was applied to the enrichment *p*-value based on chromatin accessibility and gene expression respectively, considering the number of cell types and GWAS studies tested.

### Fine-mapping of GWAS and eQTL variants

GWAS loci were fine-mapped based on the summary statistics of GWAS studies. For each GWAS study, the SNPs with * < 5 × 10^-8^ and present in 1000 genome (phase 3) European population were considered and were categorized into the LD blocks identified by a previous study. Within each LD block, the posterior inclusion probability (PIP) of each SNP and credible set of SNPs were calculated using the susieR package (L=10) ^113^. Similarly, eQTL variants were fine-mapped based on the summary statistics of bulk retinal eQTLs. The colocalization analysis of GWAS signal and bulk eQTL signal was conducted using the coloc R package ^114^. The motif disrupt effect of SNPs was predicted by the motifbreakR R package^115^.

### Query to reference mapping using scArches

The HRCA cell type labeling enables automated cell type annotation using scArches^81^. We trained query-to-reference models using scArches, using default parameters as recommended in their core tutorials. Models were trained during 20 epochs for scVI, scANVI, and label transfer on sc and sn cells from the healthy reference and using batch information during the integration benchmark. Additional cell type sub-annotations were used, based on clustering and marker-based selection per major classes. Only healthy donors were considered to generate reference models.

To test the cell mapping and uncertainty estimations in new samples, we used age-related macular degeneration samples (AMD) related to 17 donors. As validation of the label transfer accuracy, we pre-annotated one of the disease samples using scPred, obtaining 98% agreement in labels. Label uncertainties per major class mapped on AMD donors were analyzed as a single-variable distribution, and we defined a percentile threshold of 97.5% to label cells as high- or low-uncertainty based on this value. Selection of visualization of marker genes across categories was done on each cell type, between both uncertainty categories, using Scanpy ^91^. Overlap between selected marker genes AMD-related genes was inspected using the ontology term Macular Degeneration (DOID:4448) from the DISEASES database ^116^.

## Data availability

The landing page of the HRCA data resources is accessible at https://rchenlab.github.io/resources/human-atlas.html. Raw sequencing data files, processed Cell Ranger data files, and sample metadata information files of the HRCA have been deposited in the HCA DCP. Additionally, raw and normalized count matrices, cell type annotations, and multi-omics embeddings are also publicly available through the CELLxGENE collection (https://cellxgene.cziscience.com/collections/4c6eaf5c-6d57-4c76-b1e9-60df8c655f1e). The HRCA is also accessible at the UCSC Cell Browser (https://cells-test.gi.ucsc.edu/?ds=retina-atlas+rna-seq+chen) and the Single Cell Portal.

## Code availability

All code used for the HRCA project can be found in the HRCA reproducibility GitHub repository (https://github.com/RCHENLAB/HRCA_reproducibility). The pipeline to process the unpublished and collected public datasets is accessible at https://github.com/lijinbio/cellqc. Scripts related to the benchmark study, integration pipeline, and label transfer using scArches are available at https://github.com/theislab/HRCA-reproducibility.

## Acknowledgements

This project was funded by the Chan Zuckerberg Initiative (CZI) 2019-002425 and 2021-239847 to R.C., and CZI 2021-237885 to T.S. Additionally, next-generation sequencing (NGS) was performed on instrument supported by the National Institutes of Health (NIH) shared instrument grant S10OD023469 to R.C. This work was also funded by NIH EY028633 and U01 MH105960 and CZI award CZF-2019-002459 to J.R.S., and NIH/NEI EY012543 (to SC) and EY002687 (to WU-DOVS). This publication is part of the Human Cell Atlas (http://www.humancellatlas.org/publications/). We would also like to acknowledge the feedback and discussions from the members of the HCA Eye Biological Network.

## Author contributions

J.L., J.W., and R.C. conceptualized and designed the study. R.C. supervised the work.

M.M.D. and J.T.S. collected samples. X.C. and Y.L. generated snRNA-seq and snATAC-seq data in this study. I.L.I. performed the benchmark study for data integration of RNA-seq datasets in this study and label transfer analyses. J.L. and J.W. compiled dataset collection for public snRNA-seq/scRNA-seq and snATAC-seq datasets. J.L. performed data integration for RNA-seq datasets. J.W. performed data integration for ATAC-seq datasets. J.L. performed atlas construction, annotation, and data dissemination of the atlas. J.W. conducted multi-omics analysis, gene expression across covariates, and genetic variants analysis. I.L.I., M.D.L., and F.J.T. provided input to various analysis methods. N.M.T. and K.S. provided input for various annotations. A.M., W.Y. and J.R.S. collected, analyzed and provided processed data from an unpublished dataset. Y.Z. and S.C. advised and performed massively parallel reporter assays in mouse retina. J.L. and J.W. wrote the first draft of the manuscript. All authors edited the manuscript and contributed to critical revisions of the manuscript.

## Extended Data Figure legends

**Extended Data Figure 1.**
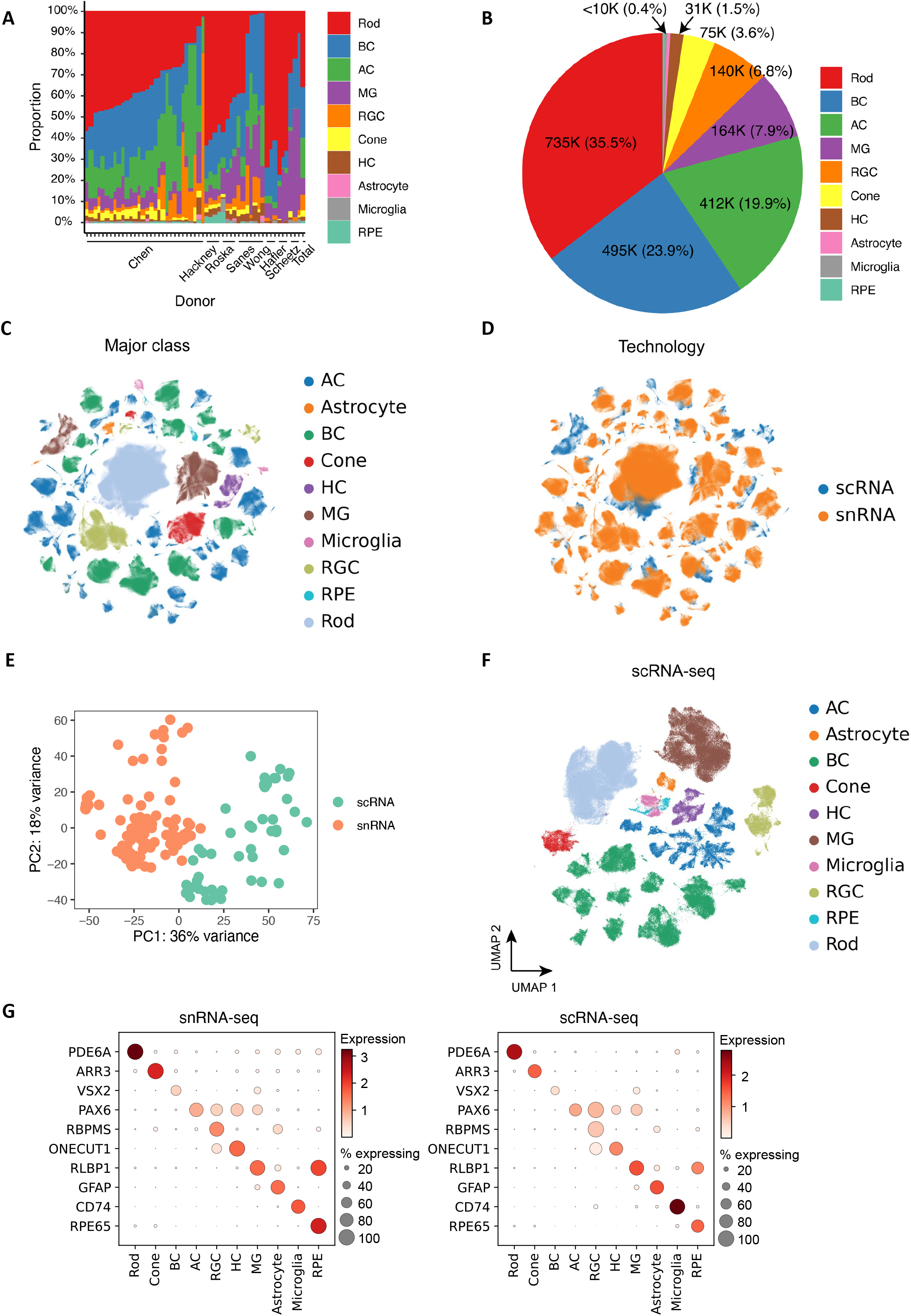
Overview of the HRCA. A. Cell proportion distribution of major classes among donors. The x-axis corresponds to each donor, and the y-axis is the cell proportion of major classes. The last bar is the cell proportion across total cells. B. A pie chart illustrating the number of cells for major classes and their proportions. C. Integration of datasets from snRNA-seq and scRNA-seq datasets. The cells are colored by major classes. D. The atlas is colored by the two technologies: snRNA-seq (in coral) and scRNA-seq (in blue). E. The distribution of transcriptomic data for 152 samples obtained from snRNA-seq and scRNA-seq technologies. Each sample is colored by the technology used. F. The atlas of scRNA-seq data, with major classes represented using different colors. G. Dot plots illustrating the distribution of expression levels of marker genes for major cell classes in snRNA-seq (on the left) and scRNA-seq data (on the right).

**Extended Data Figure 2.**
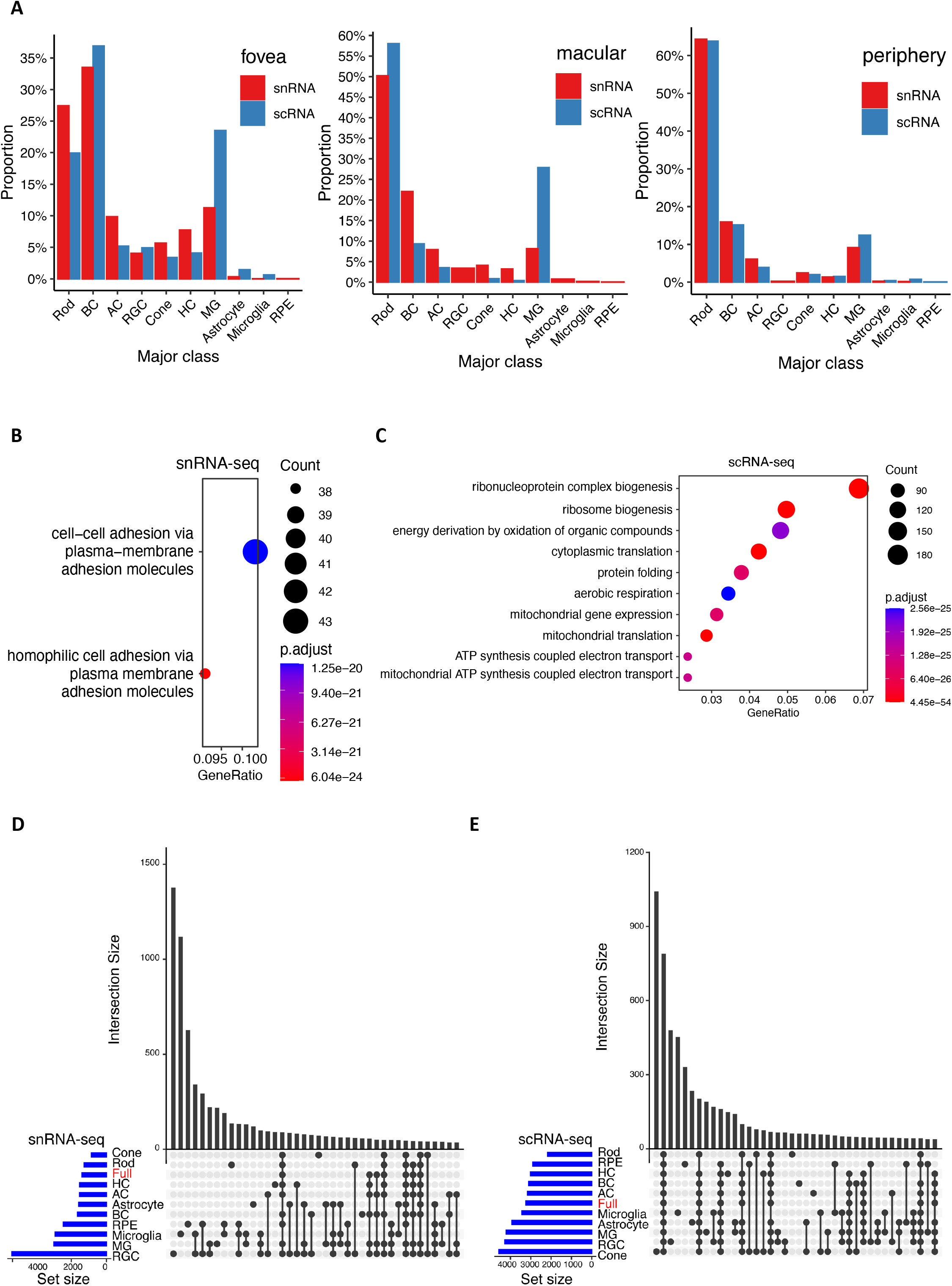
Comparison between single-nuclei and single-cell technologies. A. Cell proportion of major class of samples between snRNA-seq and scRNA-seq in fovea, macular, and periphery tissue regions. The red bar represents cell proportions of major classes in snRNA-seq samples, and the blue bar represents cell proportions of scRNA-seq samples. B. Enriched GO BPs of 1,387 over-expressed genes in snRNA-seq data. C. Enriched GO BPs of 3,242 over-expressed genes in scRNA-seq data. D. Shared genes over-expressed in snRNA-seq data among major cell classes. The “Full” (in red) is genes over-expressed in snRNA-seq data regardless of cell classes. E. Shared of genes derived from scRNA-seq data.

**Extended Data Figure 3.**
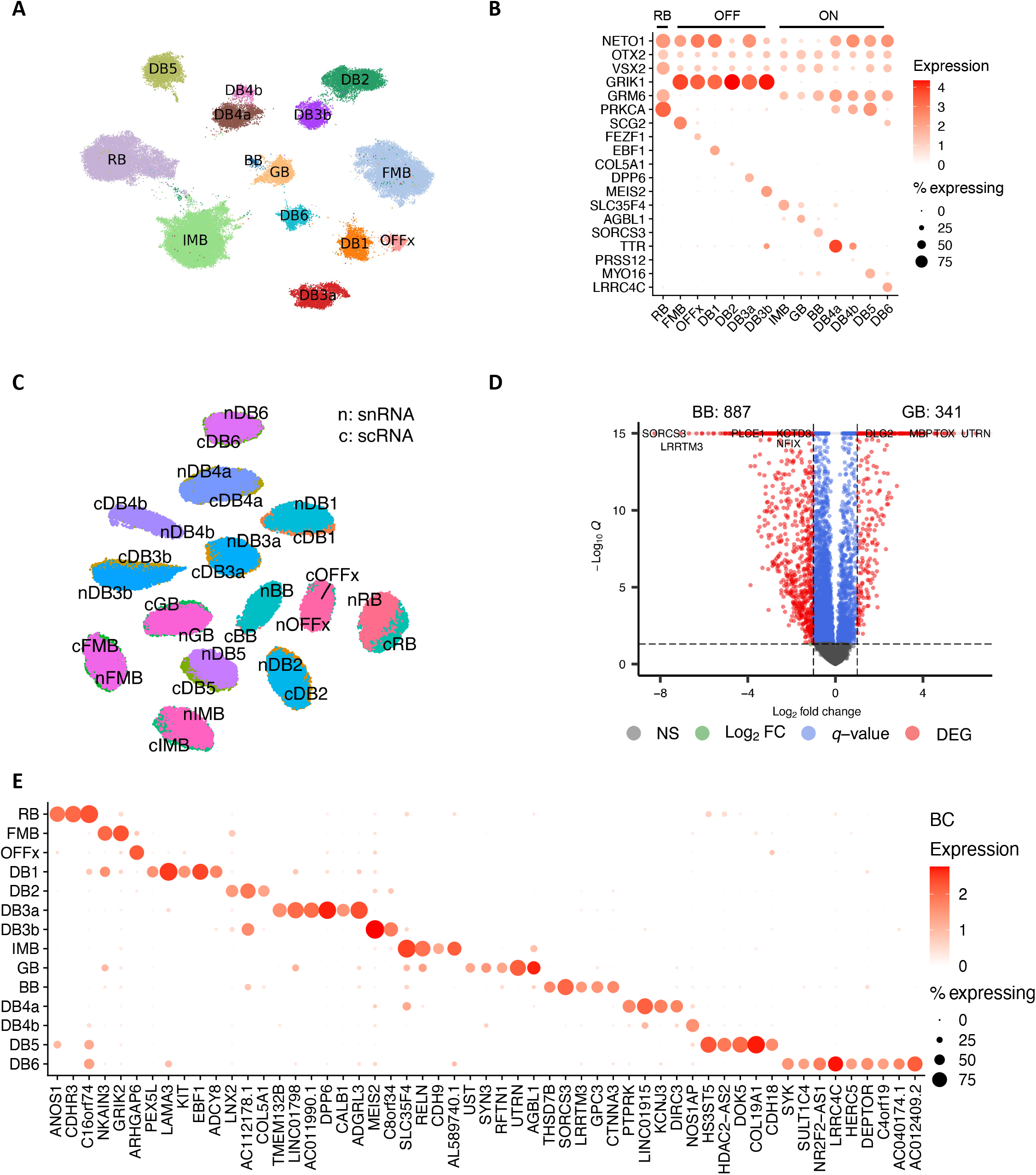
transcriptomic signature of bipolar cells. A. UMAP visualization of BC cells based on single-cell transcriptome data. B. Dot plot of the distribution of marker gene expression by the single-cell measurements. C. Co-embedding between snRNA-seq and scRNA-seq cells. The label names are prefixed by “n” for snRNA and “c” for scRNA. D. Volcano plot of differentially expressed genes between GB and BB of the snRNA-seq datasets. Differentially expressed genes were identified under |log2 fold change|>1 and *q*-value<0.05. E. Predicted markers per BC cell type by the binary classification analysis using snRNA-seq datasets. Rows are BC cell types, and columns represent novel markers.

**Extended Data Figure 4.**
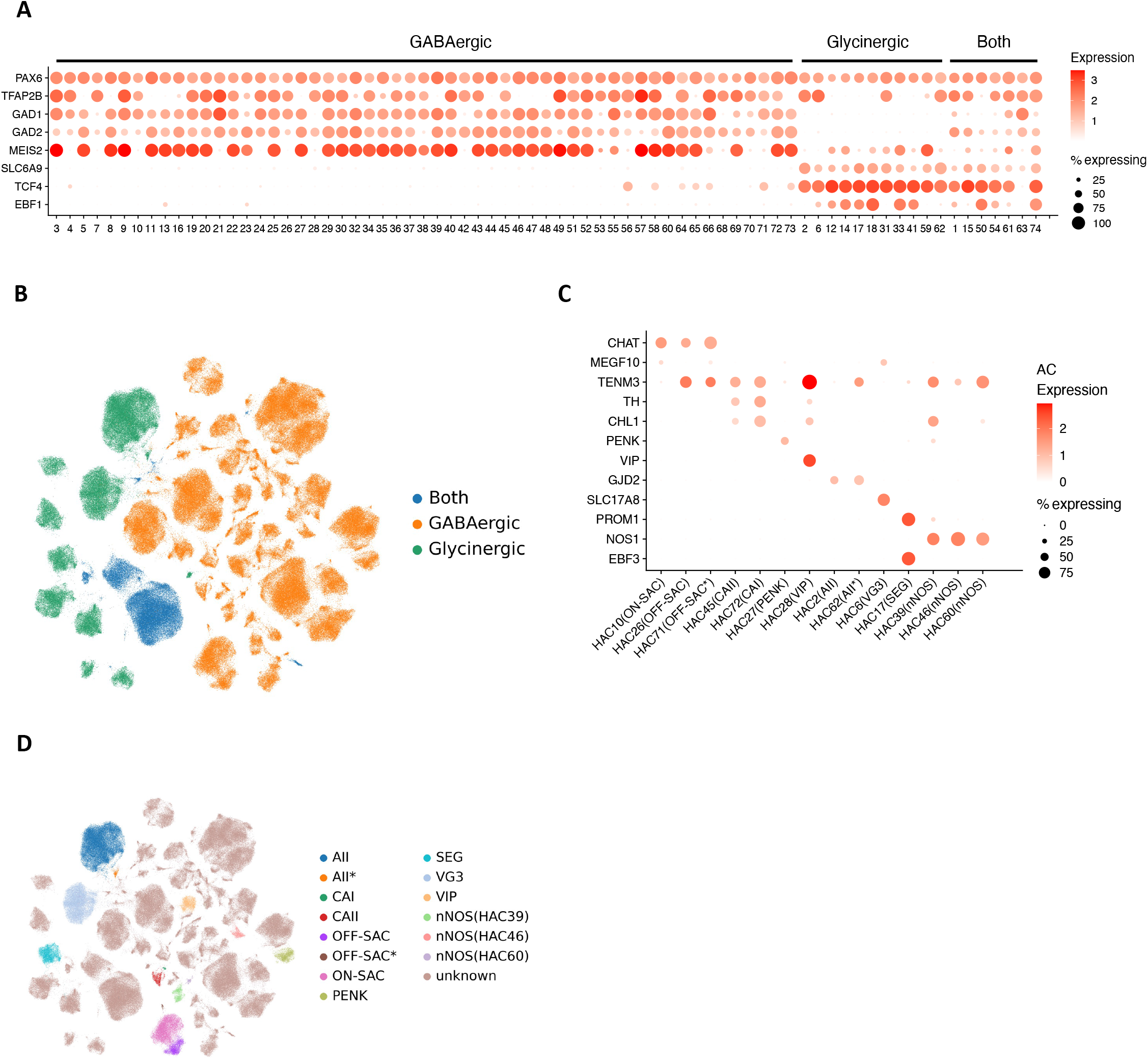
Annotation of amacrine cells. A. Dot plot of AC cell clusters by markers to identify AC subclasses for GABAergic, Glycinergic, and Both. *PAX6* and *TFAP2B* were used as AC pan-markers. *GAD1/GAD2* were used for GABAergic ACs, and *SLC6A9* was used for the Glycinergic ACs. *MEIS2*, *TCF4*, and *EBF1* were also included in the dot plot. B. UMAP of AC cells, colored by the four AC groups. C. Dot plot of 14 AC cell clusters with known markers. The cell type names are indicated in parentheses next to the cluster IDs. D. UMAP visualization of AC cells, colored by the 14 clusters with cell type names. The rest of the clusters are colored as “unknown” without existing names.

**Extended Data Figure 5.**
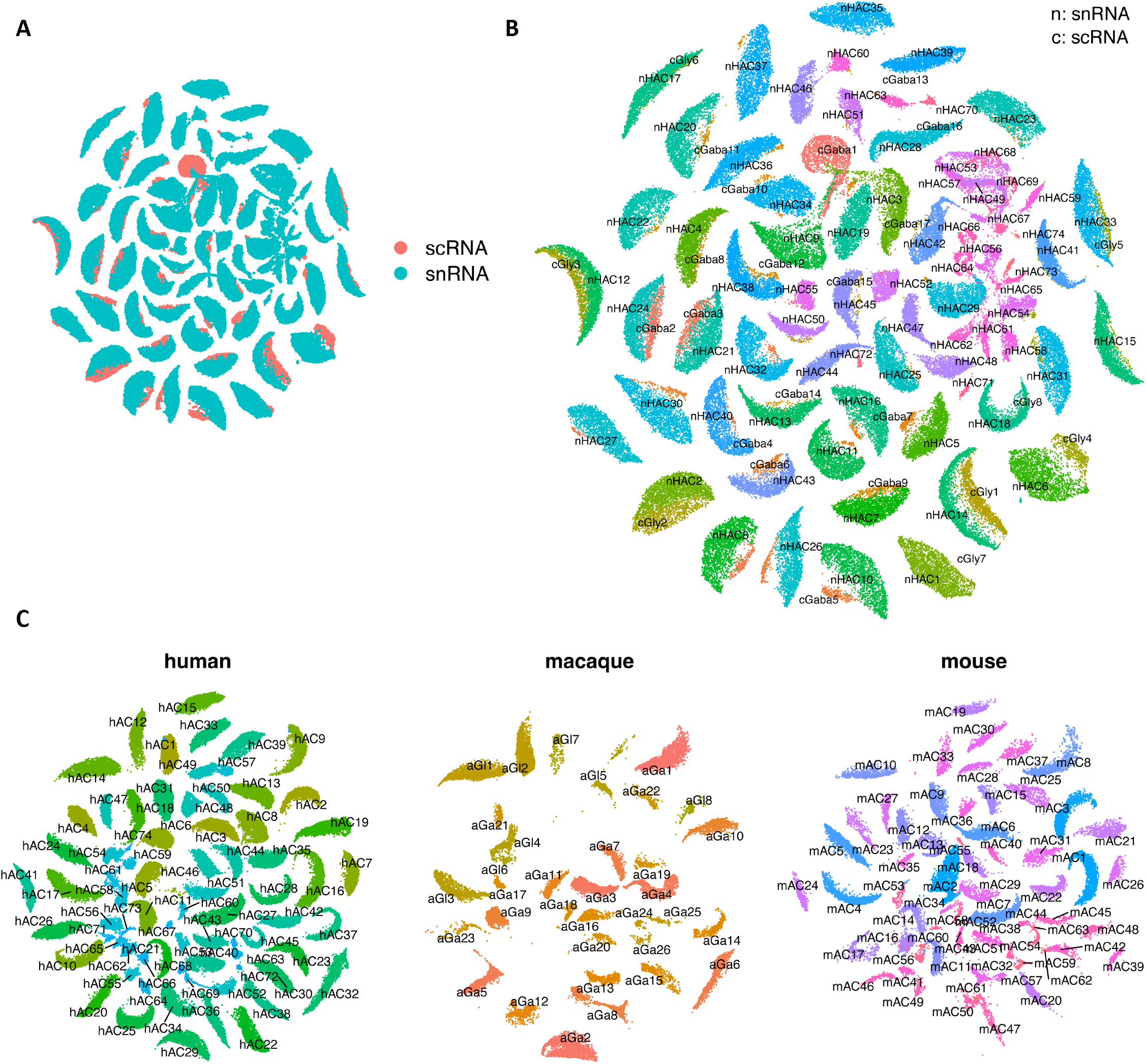
Cross-mapping for human amacrine cells. A. SATURN co-embedding visualization of AC cell types between snRNA-seq and scRNA-seq. AC cells are colored by the two technologies. B. The same SATURN co-embedding with AC type labels color-coded on top of clusters. Labels are prefixed with “n” for snRNA-seq datasets and “c” for scRNA-seq data. C. SATURN co-embedding visualization of AC types across human, macaque and mouse species. AC cell labels for the three species are overlaid on clusters. Labels are prefixed with “h” for human, “a” for macaque, and “m” for mouse.

**Extended Data Figure 6.**
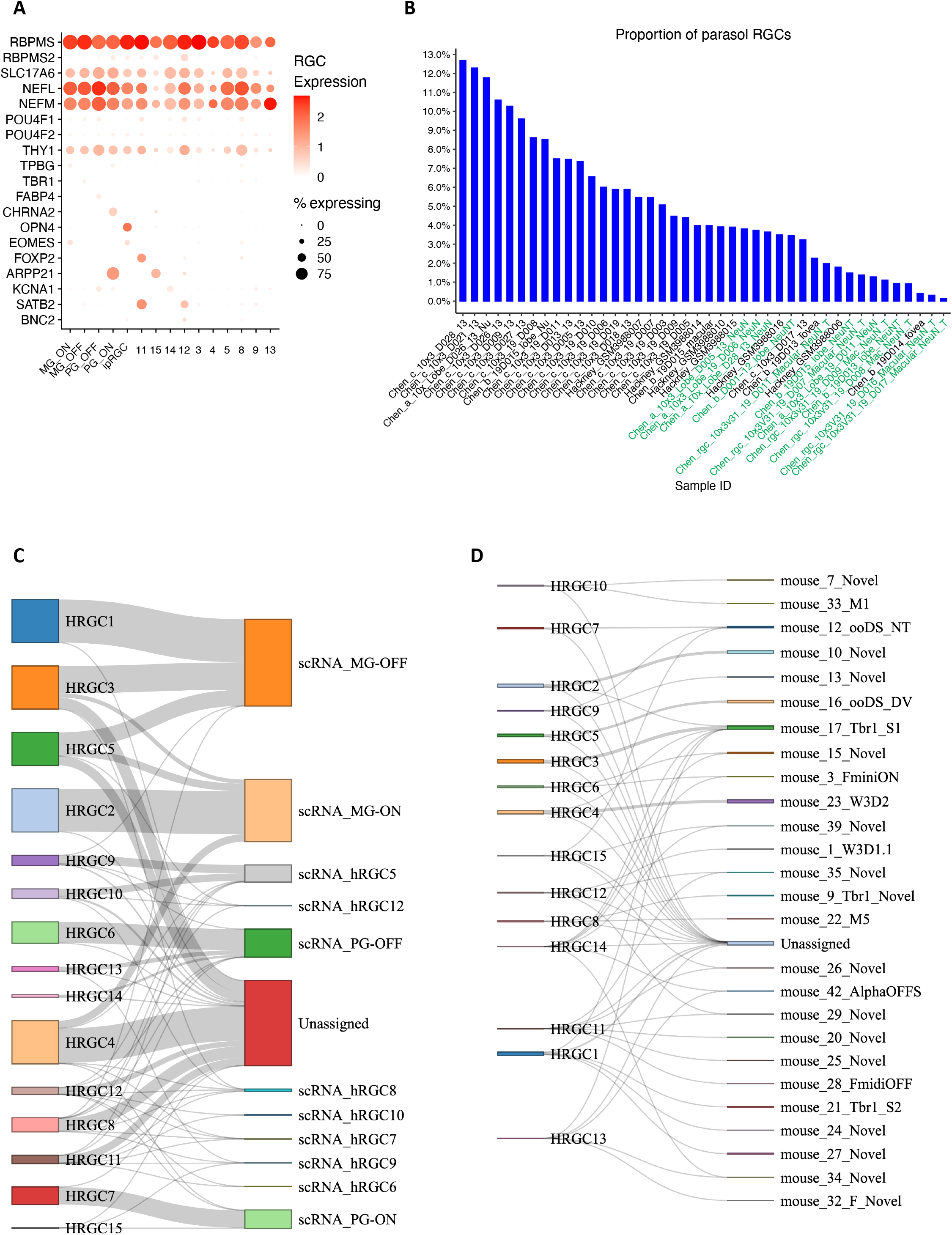
Annotation of retinal ganglion cells. A. Dot plot of RGC cell clusters with existing markers. B. The proportion of parasol RGCs within the RGC population in the samples. Samples enriched by NeuN experiments are highlighted in green. C. Sankey diagram depicting the relationship between RGC clusters from snRNA-seq datasets and the public labeling of RGC types from scRNA-seq datasets. The width of the lines is proportional to the number of cells in the mapping. D. Sankey diagram illustrating RGC types alignment between humans (left column) and mice (right column).

**Extended Data Figure 7.**
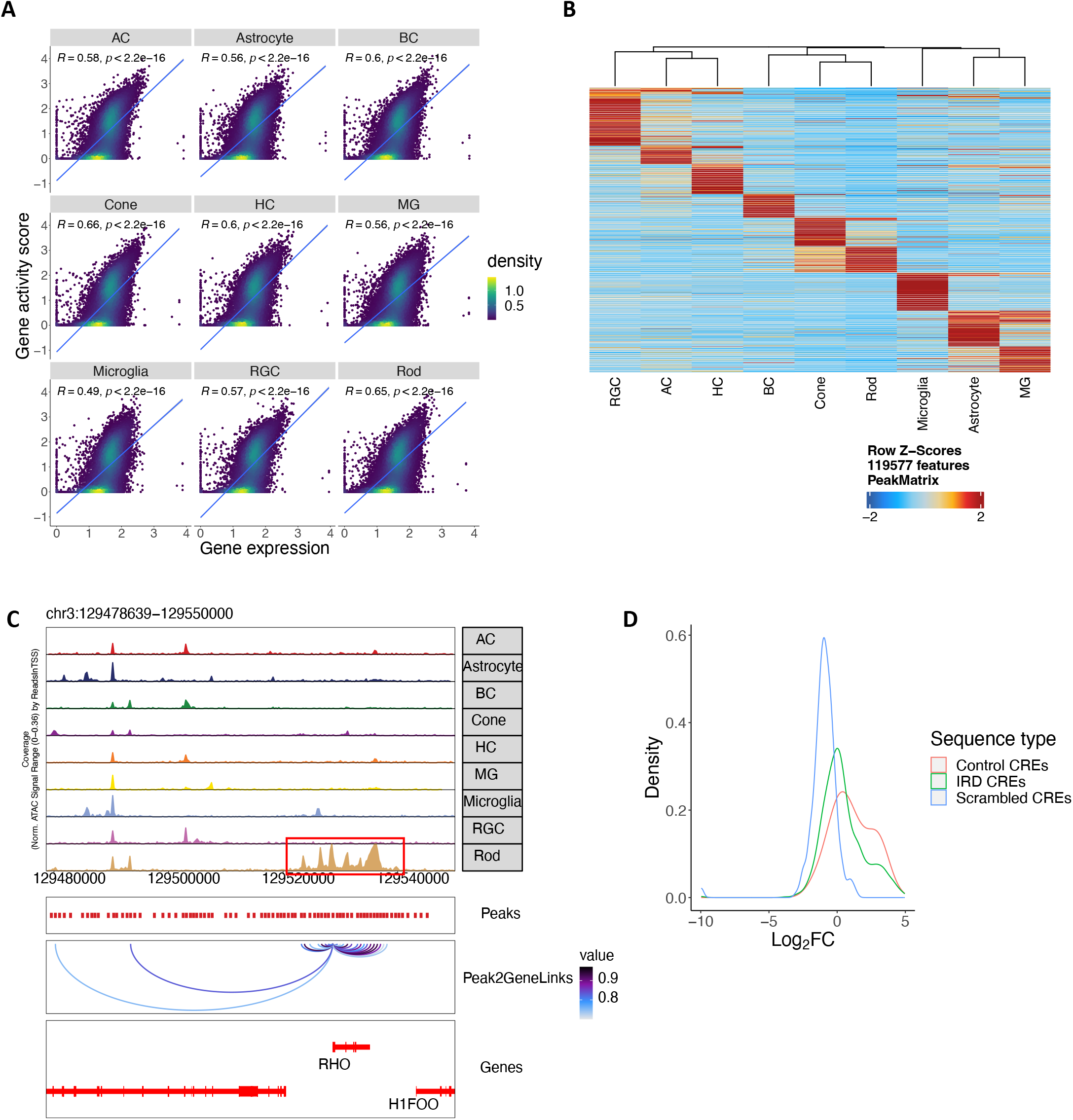
A high resolution snATAC-seq cell atlas of the human retina. A. Scatter plot showing the correlation between gene expression derived from snRNA-seq (X axis) and gene activity score derived from snATAC-seq (Y axis) from major retinal cell classes. B. Heatmap showing the chromatin accessibility of differential accessible regions (DARs) identified in major retinal cell classes. Rows represented chromatin regions specific to certain major classes, and columns corresponded to major classes. C. Genome track of the *RHO* locus showing the cell type specific chromatin accessibility in the promoter and linked cis-regulatory elements of this gene. D. Density plot showing the activity (log2*FC* value of comparison between activity of each tested sequence and the activity of a basal CRX promoter) distribution of the tested sequences by MPRAs. IRD CREs n=1,820 (green), control CREs with a variety of activities n=20 (red), Scrambled CREs n=300 (blue).

**Extended Data Figure 8.**
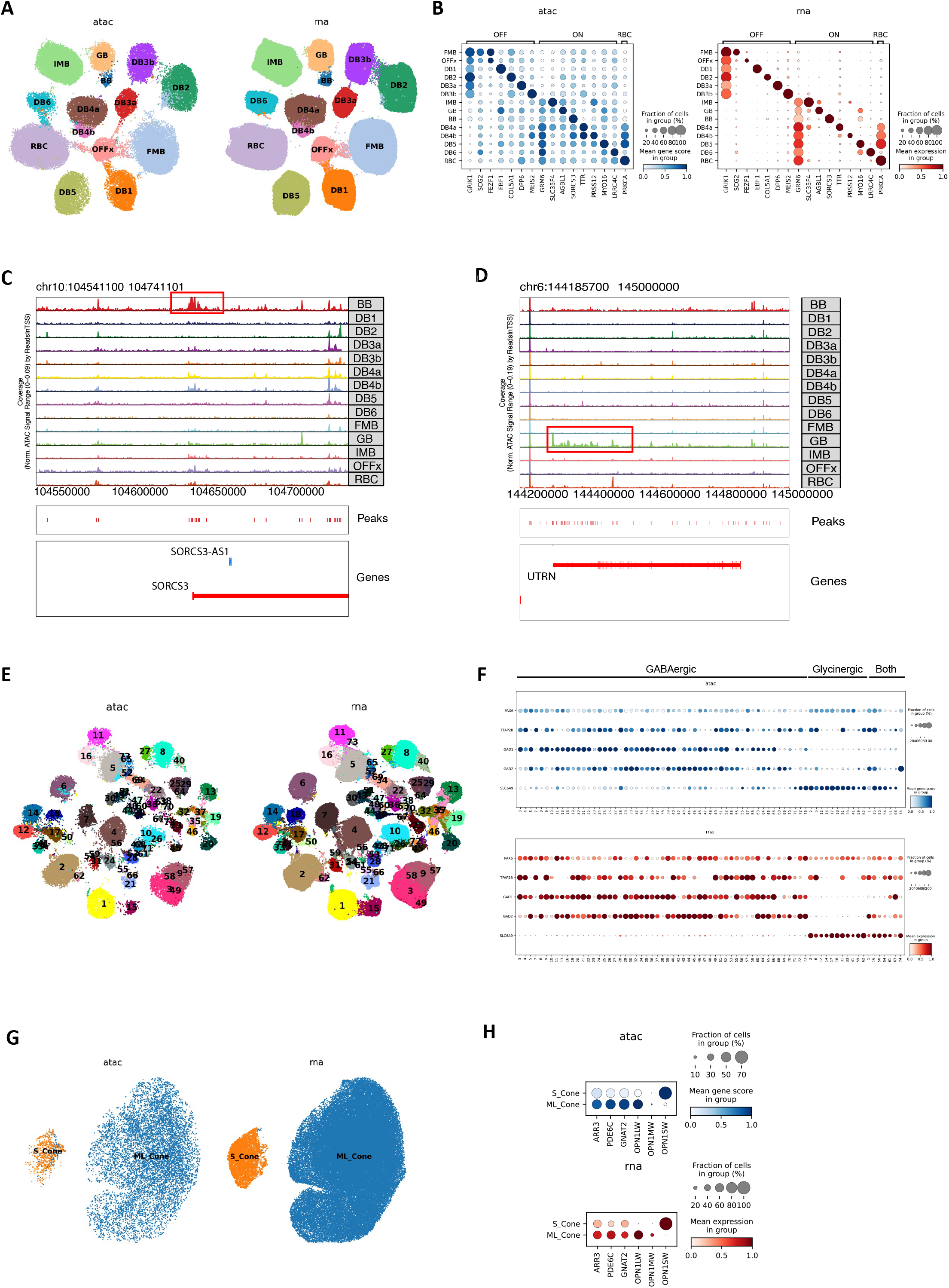
Multi-omics atlas of the human retinal subclass cell types. A. UMAP showing the co-embedding of bipolar cells (BC) from snRNA-seq and snATAC-seq were clustered into BC cell types. B. Dot plot showing marker gene expression measured by snRNA-seq and marker gene activity score derived from snATAC-seq are specific in the corresponding BC cell types. C. Genome track of SORCS3 showing the promoter of SORCS3 is specifically open in BB. D. Genome track of UTRN showing the local chromatin of UTRN is specifically open in GB. E. UMAP showing the co-embedding of amacrine cells (AC) from snRNA-seq and snATAC-seq were clustered into AC cell types. F. Dot plot showing marker gene expression measured by snRNA-seq and marker gene activity score derived from snATAC-seq are specific in the corresponding sub classes of AC types. G. UMAP showing the co-embedding of cone cells (Cone) from snRNA-seq and snATAC-seq were clustered in Cone cell types. H. Dot plot showing marker gene expression measured by snRNA-seq and marker gene activity score derived from snATAC-seq are specific in the corresponding Cone cell types.

**Extended Data Figure 9.**
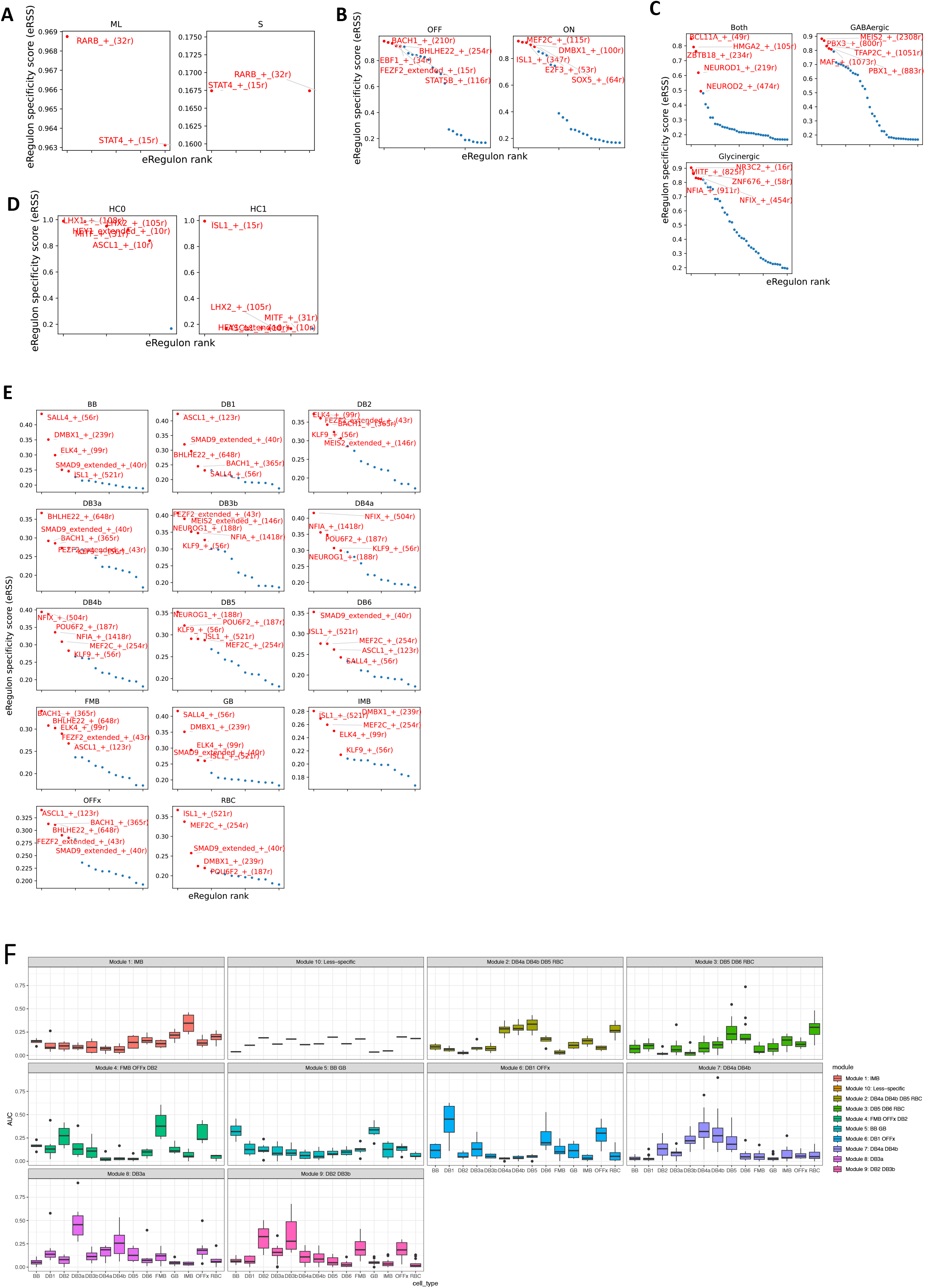
Regulon of the human retinal subclass cell types. A. Dot plot showing the distribution of regulon specificity score of regulons identified in ML- and S-Cone. B. Dot plot showing the distribution of regulon specificity score of regulons identified in OFF- and ON-BC (ON-BC include ON-Cone BC and Rod BC). C. Dot plot showing the distribution of regulon specificity score of regulons identified in GABAergic-, Glycinergic- and Both-AC. D. Dot plot showing the distribution of regulon specificity score of regulons identified in HC0 and HC1. E. Dot plot showing the distribution of regulon specificity score of regulons identified in 14 BC cell types. F. Boxplot showing the average AUC values of the regulon modules identified in BC cell types. The BC cell types with the highest AUC values were labeled in the title of each regulon module.

**Extended Data Figure 10.**
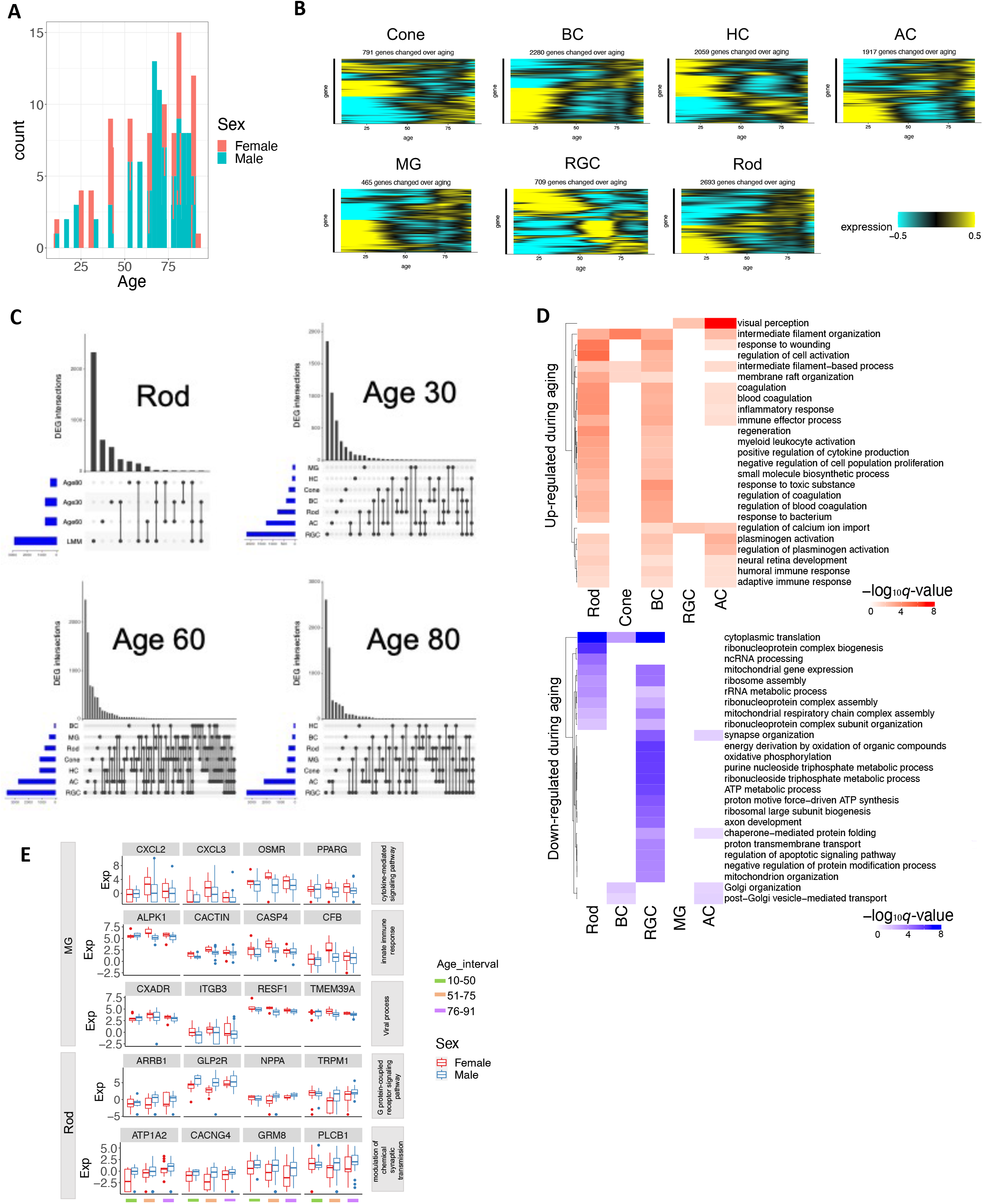
Differential gene expression during aging and associated with sex. A. Age and sex distribution of the analyzed samples. B. Heatmap showing gene expression level of differentially expressed genes (DEGs) during aging in major retinal cell classes identified with linear mixed effect model (LMM). C. UpSet plot showing the overlap of DEGs identified by LMM and sliding-window analysis at the age of 30, 60 and 80 in Rod. UpSet plot showing the number of DEGs across major retinal cell classes at the age of 30, 60 and 80, respectively. D. The GO terms significantly enriched (FDR <0.1) of DEGs during aging identified by LMM across retinal cell classes. E. The examples of DEGs between male and female associated with the enriched GO terms.

## References

1 Regev, A. et al. The Human Cell Atlas. Elife 6 (2017). 10.7554/eLife.27041

2 Rozenblatt-Rosen, O., Stubbington, M. J. T., Regev, A. & Teichmann, S. A. The Human Cell Atlas: from vision to reality. Nature 550, 451–453 (2017). 10.1038/550451a

3 Sikkema, L. et al. An integrated cell atlas of the lung in health and disease. Nat Med 29, 1563–1577 (2023). 10.1038/s41591-023-02327-2

4 Kumar, T. et al. A spatially resolved single-cell genomic atlas of the adult human breast. Nature 620, 181–191 (2023). 10.1038/s41586-023-06252-9

5 van Zyl, T. et al. Cell atlas of the human ocular anterior segment: Tissue-specific and shared cell types. Proc Natl Acad Sci U S A 119, e2200914119 (2022). 10.1073/pnas.2200914119

6 Monavarfeshani, A. et al. Transcriptomic analysis of the ocular posterior segment completes a cell atlas of the human eye. Proc Natl Acad Sci U S A 120, e2306153120 (2023). 10.1073/pnas.2306153120

7 Liang, Q. et al. A multi-omics atlas of the human retina at single-cell resolution. Cell Genomics, 100298 (2023). 10.1016/j.xgen.2023.100298

8 Hahn, J. et al. Evolution of neuronal cell classes and types in the vertebrate retina. bioRxiv (2023). 10.1101/2023.04.07.536039

9 Peng, Y. R. et al. Molecular Classification and Comparative Taxonomics of Foveal and Peripheral Cells in Primate Retina. Cell 176, 1222–1237 e1222 (2019). 10.1016/j.cell.2019.01.004

10 Yamagata, M., Yan, W. & Sanes, J. R. A cell atlas of the chick retina based on single-cell transcriptomics. Elife 10 (2021). 10.7554/eLife.63907

11 Macosko, E. Z. et al. Highly Parallel Genome-wide Expression Profiling of Individual Cells Using Nanoliter Droplets. Cell 161, 1202–1214 (2015). 10.1016/j.cell.2015.05.002

12 Shekhar, K. et al. Comprehensive Classification of Retinal Bipolar Neurons by Single-Cell Transcriptomics. Cell 166, 1308–1323 e1330 (2016). 10.1016/j.cell.2016.07.054

13 Tran, N. M. et al. Single-Cell Profiles of Retinal Ganglion Cells Differing in Resilience to Injury Reveal Neuroprotective Genes. Neuron 104, 1039–1055 e1012 (2019). 10.1016/j.neuron.2019.11.006

14 Yan, W. et al. Mouse Retinal Cell Atlas: Molecular Identification of over Sixty Amacrine Cell Types. J Neurosci 40, 5177–5195 (2020). 10.1523/JNEUROSCI.0471-20.2020

15 Yan, W. et al. Cell Atlas of The Human Fovea and Peripheral Retina. Sci Rep 10, 9802 (2020). 10.1038/s41598-020-66092-9

16 Buenrostro, J. D. et al. Single-cell chromatin accessibility reveals principles of regulatory variation. Nature 523, 486–490 (2015). 10.1038/nature14590

17 Luecken, M. D. et al. Benchmarking atlas-level data integration in single-cell genomics. Nat Methods 19, 41–50 (2022). 10.1038/s41592-021-01336-8

18 Luecken, M. D. et al. in Thirty-fifth conference on neural information processing systems datasets and benchmarks track (Round 2).

19 Cowan, C. S. et al. Cell Types of the Human Retina and Its Organoids at Single-Cell Resolution. Cell 182, 1623–1640 e1634 (2020). 10.1016/j.cell.2020.08.013

20 Orozco, L. D. et al. Integration of eQTL and a Single-Cell Atlas in the Human Eye Identifies Causal Genes for Age-Related Macular Degeneration. Cell Rep 30, 1246–1259 e1246 (2020). 10.1016/j.celrep.2019.12.082

21 Lukowski, S. W. et al. A single-cell transcriptome atlas of the adult human retina. EMBO J 38, e100811 (2019). 10.15252/embj.2018100811

22 Menon, M. et al. Single-cell transcriptomic atlas of the human retina identifies cell types associated with age-related macular degeneration. Nat Commun 10, 4902 (2019). 10.1038/s41467-019-12780-8

23 Voigt, A. P. et al. Molecular characterization of foveal versus peripheral human retina by single-cell RNA sequencing. Exp Eye Res 184, 234–242 (2019). 10.1016/j.exer.2019.05.001

24 Lopez, R., Regier, J., Cole, M. B., Jordan, M. I. & Yosef, N. Deep generative modeling for single-cell transcriptomics. Nat Methods 15, 1053–1058 (2018). 10.1038/s41592-018-0229-2

25 Santiago, C. P. et al. Comparative Analysis of Single-cell and Single-nucleus RNA-sequencing in a Rabbit Model of Retinal Detachment-related Proliferative Vitreoretinopathy. Ophthalmol Sci 3, 100335 (2023). 10.1016/j.xops.2023.100335

26 Rosen, Y. et al. Towards Universal Cell Embeddings: Integrating Single-cell RNA-seq Datasets across Species with SATURN. bioRxiv (2023). 10.1101/2023.02.03.526939

27 Bakken, T. E. et al. Single-cell and single-nucleus RNA-seq uncovers shared and distinct axes of variation in dorsal LGN neurons in mice, non-human primates, and humans. Elife 10 (2021). 10.7554/eLife.64875

28 Kolb, H., Fernandez, E. & Nelson, R. Webvision: the organization of the retina and visual system [Internet]. (1995).

29 Hughes, A. E. O., Myers, C. A. & Corbo, J. C. A massively parallel reporter assay reveals context-dependent activity of homeodomain binding sites in vivo. Genome Res 28, 1520–1531 (2018). 10.1101/gr.231886.117

30 Shepherdson, J. L. et al. Pathogenic variants in CRX have distinct cis - regulatory effects on enhancers and silencers in photoreceptors. bioRxiv (2023). 10.1101/2023.05.27.542576

31 Bravo Gonzalez-Blas, C., et al. SCENIC+: single-cell multiomic inference of enhancers and gene regulatory networks. Nat Methods 20, 1355–1367 (2023). 10.1038/s41592-023-01938-4

32 de Melo, J. et al. Lhx2 Is an Essential Factor for Retinal Gliogenesis and Notch Signaling. J Neurosci 36, 2391–2405 (2016). 10.1523/JNEUROSCI.3145-15.2016

33 Sapkota, D. et al. Onecut1 and Onecut2 redundantly regulate early retinal cell fates during development. Proc Natl Acad Sci U S A 111, E4086–4095 (2014). 10.1073/pnas.1405354111

34 Clark, B. S. et al. Single-Cell RNA-Seq Analysis of Retinal Development Identifies NFI Factors as Regulating Mitotic Exit and Late-Born Cell Specification. Neuron 102, 1111–1126 e1115 (2019). 10.1016/j.neuron.2019.04.010

35 Yamamoto, H., Kon, T., Omori, Y. & Furukawa, T. Functional and Evolutionary Diversification of Otx2 and Crx in Vertebrate Retinal Photoreceptor and Bipolar Cell Development. Cell Rep 30, 658–671 e655 (2020). 10.1016/j.celrep.2019.12.072

36 Remez, L. A. et al. Pax6 is essential for the generation of late-born retinal neurons and for inhibition of photoreceptor-fate during late stages of retinogenesis. Dev Biol 432, 140–150 (2017). 10.1016/j.ydbio.2017.09.030

37 Cheng, H. et al. Photoreceptor-specific nuclear receptor NR2E3 functions as a transcriptional activator in rod photoreceptors. Hum Mol Genet 13, 1563–1575 (2004). 10.1093/hmg/ddh173

38 Kaewkhaw, R. et al. Transcriptome Dynamics of Developing Photoreceptors in Three-Dimensional Retina Cultures Recapitulates Temporal Sequence of Human Cone and Rod Differentiation Revealing Cell Surface Markers and Gene Networks. Stem Cells 33, 3504–3518 (2015). 10.1002/stem.2122

39 Cao, Z. J. & Gao, G. Multi-omics single-cell data integration and regulatory inference with graph-linked embedding. Nat Biotechnol 40, 1458–1466 (2022). 10.1038/s41587-022-01284-4

40 Campello, L. et al. Aging of the Retina: Molecular and Metabolic Turbulences and Potential Interventions. Annu Rev Vis Sci 7, 633–664 (2021). 10.1146/annurev-vision-100419-114940

41 Aninye, I. O. et al. The roles of sex and gender in women’s eye health disparities in the United States. Biol Sex Differ 12, 57 (2021). 10.1186/s13293-021-00401-3

42 Park, D. H., Connor, K. M. & Lambris, J. D. The Challenges and Promise of Complement Therapeutics for Ocular Diseases. Front Immunol 10, 1007 (2019). 10.3389/fimmu.2019.01007

43 Khandhadia, S., Cipriani, V., Yates, J. R. & Lotery, A. J. Age-related macular degeneration and the complement system. Immunobiology 217, 127–146 (2012). 10.1016/j.imbio.2011.07.019

44 Armento, A., Ueffing, M. & Clark, S. J. The complement system in age-related macular degeneration. Cell Mol Life Sci 78, 4487–4505 (2021). 10.1007/s00018-021-03796-9

45 Geerlings, M. J., de Jong, E. K. & den Hollander, A. I. The complement system in age-related macular degeneration: A review of rare genetic variants and implications for personalized treatment. Mol Immunol 84, 65–76 (2017). 10.1016/j.molimm.2016.11.016

46 Ma, H., Yang, F. & Ding, X. Q. Inhibition of thyroid hormone signaling protects retinal pigment epithelium and photoreceptors from cell death in a mouse model of age-related macular degeneration. Cell Death Dis 11, 24 (2020). 10.1038/s41419-019-2216-7

47 Xu, H. & Chen, M. Targeting the complement system for the management of retinal inflammatory and degenerative diseases. Eur J Pharmacol 787, 94–104 (2016). 10.1016/j.ejphar.2016.03.001

48 Nuzzi, R., Scalabrin, S., Becco, A. & Panzica, G. Gonadal Hormones and Retinal Disorders: A Review. Front Endocrinol (Lausanne) 9, 66 (2018). 10.3389/fendo.2018.00066

49 Liu, K. et al. Genetic variation reveals the influence of steroid hormones on the risk of retinal neurodegenerative diseases. Front Endocrinol (Lausanne) 13, 1088557 (2022). 10.3389/fendo.2022.1088557

50 Lu, T. C. et al. Aging Fly Cell Atlas identifies exhaustive aging features at cellular resolution. Science 380, eadg0934 (2023). 10.1126/science.adg0934

51 Whitacre, C. C., Reingold, S. C. & O’Looney, P. A. A gender gap in autoimmunity. Science 283, 1277–1278 (1999). 10.1126/science.283.5406.1277

52 Beeson, P. B. Age and sex associations of 40 autoimmune diseases. Am J Med 96, 457–462 (1994). 10.1016/0002-9343(94)90173-2

53 Oertelt-Prigione, S. The influence of sex and gender on the immune response. Autoimmun Rev 11, A479–485 (2012). 10.1016/j.autrev.2011.11.022

54 Ansar Ahmed, S., Penhale, W. J. & Talal, N. Sex hormones, immune responses, and autoimmune diseases. Mechanisms of sex hormone action. Am J Pathol 121, 531–551 (1985).

55 Huan, T. et al. Identifying Novel Genes and Variants in Immune and Coagulation Pathways Associated with Macular Degeneration. Ophthalmol Sci 3, 100206 (2023). 10.1016/j.xops.2022.100206

56 Rudnicka, A. R. et al. Age and gender variations in age-related macular degeneration prevalence in populations of European ancestry: a meta-analysis. Ophthalmology 119, 571–580 (2012). 10.1016/j.ophtha.2011.09.027

57 Li, M. et al. Autophagy in glaucoma pathogenesis: Therapeutic potential and future perspectives. Front Cell Dev Biol 10, 1068213 (2022). 10.3389/fcell.2022.1068213

58 Dixon, A. et al. Autophagy deficiency protects against ocular hypertension and neurodegeneration in experimental and spontanous glaucoma mouse models. Cell Death Dis 14, 554 (2023). 10.1038/s41419-023-06086-3

59 Kapetanakis, V. V. et al. Global variations and time trends in the prevalence of primary open angle glaucoma (POAG): a systematic review and meta-analysis. Br J Ophthalmol 100, 86–93 (2016). 10.1136/bjophthalmol-2015-307223

60 Zhang, N., Wang, J., Li, Y. & Jiang, B. Prevalence of primary open angle glaucoma in the last 20 years: a meta-analysis and systematic review. Sci Rep 11, 13762 (2021). 10.1038/s41598-021-92971-w

61 Finucane, H. K. et al. Partitioning heritability by functional annotation using genome-wide association summary statistics. Nat Genet 47, 1228–1235 (2015). 10.1038/ng.3404

62. de Leeuw, C. A., Mooij, J. M., Heskes, T. & Posthuma, D. MAGMA: generalized gene-set analysis of GWAS data. PLoS Comput Biol 11, e1004219 (2015). 10.1371/journal.pcbi.1004219

63 Fritsche, L. G. et al. A large genome-wide association study of age-related macular degeneration highlights contributions of rare and common variants. Nat Genet 48, 134–143 (2016). 10.1038/ng.3448

64 Currant, H. et al. Sub-cellular level resolution of common genetic variation in the photoreceptor layer identifies continuum between rare disease and common variation. PLoS Genet 19, e1010587 (2023). 10.1371/journal.pgen.1010587

65 Gharahkhani, P. et al. Genome-wide meta-analysis identifies 127 open-angle glaucoma loci with consistent effect across ancestries. Nat Commun 12, 1258 (2021). 10.1038/s41467-020-20851-4

66 Khawaja, A. P. et al. Genome-wide analyses identify 68 new loci associated with intraocular pressure and improve risk prediction for primary open-angle glaucoma. Nat Genet 50, 778–782 (2018). 10.1038/s41588-018-0126-8

67 Springelkamp, H. et al. New insights into the genetics of primary open-angle glaucoma based on meta-analyses of intraocular pressure and optic disc characteristics. Hum Mol Genet 26, 438–453 (2017). 10.1093/hmg/ddw399

68 Hysi, P. G. et al. Meta-analysis of 542,934 subjects of European ancestry identifies new genes and mechanisms predisposing to refractive error and myopia. Nat Genet 52, 401–407 (2020). 10.1038/s41588-020-0599-0

69 Kemp, J. P. et al. Identification of 153 new loci associated with heel bone mineral density and functional involvement of GPC6 in osteoporosis. Nat Genet 49, 1468–1475 (2017). 10.1038/ng.3949

70 Jiang, L., Zheng, Z., Fang, H. & Yang, J. A generalized linear mixed model association tool for biobank-scale data. Nat Genet 53, 1616–1621 (2021). 10.1038/s41588-021-00954-4

71 Collantes, E. R. A. et al. EFEMP1 rare variants cause familial juvenile-onset open-angle glaucoma. Hum Mutat 43, 240–252 (2022). 10.1002/humu.24320

72 Liu, Y. & Allingham, R. R. Major review: Molecular genetics of primary open-angle glaucoma. Exp Eye Res 160, 62–84 (2017). 10.1016/j.exer.2017.05.002

73 Guo, F. et al. ABCF1 extrinsically regulates retinal pigment epithelial cell phagocytosis. Mol Biol Cell 26, 2311–2320 (2015). 10.1091/mbc.E14-09-1343

74 Youngblood, H., Robinson, R., Sharma, A. & Sharma, S. Proteomic Biomarkers of Retinal Inflammation in Diabetic Retinopathy. Int J Mol Sci 20 (2019). 10.3390/ijms20194755

75 Lu, Y. et al. Single-Cell Analysis of Human Retina Identifies Evolutionarily Conserved and Species-Specific Mechanisms Controlling Development. Dev Cell 53, 473–491 e479 (2020). 10.1016/j.devcel.2020.04.009

76 Muranishi, Y. et al. An essential role for RAX homeoprotein and NOTCH-HES signaling in Otx2 expression in embryonic retinal photoreceptor cell fate determination. J Neurosci 31, 16792–16807 (2011). 10.1523/JNEUROSCI.3109-11.2011

77 Megill, C. et al. cellxgene: a performant, scalable exploration platform for high dimensional sparse matrices. bioRxiv, 2021.2004.2005.438318 (2021). 10.1101/2021.04.05.438318

78 Speir, M. L. et al. UCSC Cell Browser: visualize your single-cell data. Bioinformatics 37, 4578–4580 (2021). 10.1093/bioinformatics/btab503

79 Tarhan, L. et al. Single Cell Portal: an interactive home for single-cell genomics data. bioRxiv (2023). 10.1101/2023.07.13.548886

80 Friedman, R. Z. et al. Information content differentiates enhancers from silencers in mouse photoreceptors. Elife 10 (2021). 10.7554/eLife.67403

81 Lotfollahi, M. et al. Mapping single-cell data to reference atlases by transfer learning. Nat Biotechnol 40, 121–130 (2022). 10.1038/s41587-021-01001-7

82 Owen, L. A. et al. The Utah Protocol for Postmortem Eye Phenotyping and Molecular Biochemical Analysis. Invest Ophthalmol Vis Sci 60, 1204–1212 (2019). 10.1167/iovs.18-24254

83 Zheng, G. X. et al. Massively parallel digital transcriptional profiling of single cells. Nat Commun 8, 14049 (2017). 10.1038/ncomms14049

84 Lun, A. T. L. et al. EmptyDrops: distinguishing cells from empty droplets in droplet-based single-cell RNA sequencing data. Genome Biol 20, 63 (2019). 10.1186/s13059-019-1662-y

85 Heiser, C. N., Wang, V. M., Chen, B., Hughey, J. J. & Lau, K. S. Automated quality control and cell identification of droplet-based single-cell data using dropkick. Genome Res 31, 1742–1752 (2021). 10.1101/gr.271908.120

86 Young, M. D. & Behjati, S. SoupX removes ambient RNA contamination from droplet-based single-cell RNA sequencing data. Gigascience 9 (2020). 10.1093/gigascience/giaa151

87 McGinnis, C. S., Murrow, L. M. & Gartner, Z. J. DoubletFinder: Doublet Detection in Single-Cell RNA Sequencing Data Using Artificial Nearest Neighbors. Cell Syst 8, 329–337 e324 (2019). 10.1016/j.cels.2019.03.003

88 Alquicira-Hernandez, J., Sathe, A., Ji, H. P., Nguyen, Q. & Powell, J. E. scPred: accurate supervised method for cell-type classification from single-cell RNA-seq data. Genome Biol 20, 264 (2019). 10.1186/s13059-019-1862-5

89 Stuart, T. et al. Comprehensive Integration of Single-Cell Data. Cell 177, 1888–1902 e1821 (2019). 10.1016/j.cell.2019.05.031

90 Hao, Y. et al. Integrated analysis of multimodal single-cell data. Cell 184, 3573–3587 e3529 (2021). 10.1016/j.cell.2021.04.048

91 Wolf, F. A., Angerer, P. & Theis, F. J. SCANPY: large-scale single-cell gene expression data analysis. Genome Biol 19, 15 (2018). 10.1186/s13059-017-1382-0

92 Traag, V. A., Waltman, L. & van Eck, N. J. From Louvain to Leiden: guaranteeing well-connected communities. Sci Rep 9, 5233 (2019). 10.1038/s41598-019-41695-z

93 McInnes, L., Healy, J. & Melville, J. Umap: Uniform manifold approximation and projection for dimension reduction. arXiv preprint *arXiv:1802.03426* (2018).

94 Miao, Z. et al. Putative cell type discovery from single-cell gene expression data. Nat Methods 17, 621–628 (2020). 10.1038/s41592-020-0825-9

95 Crow, M., Paul, A., Ballouz, S., Huang, Z. J. & Gillis, J. Characterizing the replicability of cell types defined by single cell RNA-sequencing data using MetaNeighbor. Nat Commun 9, 884 (2018). 10.1038/s41467-018-03282-0

96 Love, M. I., Huber, W. & Anders, S. Moderated estimation of fold change and dispersion for RNA-seq data with DESeq2. Genome Biol 15, 550 (2014). 10.1186/s13059-014-0550-8

97 Benjamini, Y. & Hochberg, Y. Controlling the false discovery rate: a practical and powerful approach to multiple testing. Journal of the Royal statistical society: series B (Methodological) 57, 289–300 (1995).

98 Wu, T. et al. clusterProfiler 4.0: A universal enrichment tool for interpreting omics data. Innovation (Camb) 2, 100141 (2021). 10.1016/j.xinn.2021.100141

99 Conway, J. R., Lex, A. & Gehlenborg, N. UpSetR: an R package for the visualization of intersecting sets and their properties. Bioinformatics 33, 2938–2940 (2017). 10.1093/bioinformatics/btx364

100 Blighe, K., Rana, S. & Lewis, M. EnhancedVolcano: Publication-ready volcano plots with enhanced colouring and labeling. (2022).

101 Chen, H. & Boutros, P. C. VennDiagram: a package for the generation of highly-customizable Venn and Euler diagrams in R. BMC Bioinformatics 12, 35 (2011). 10.1186/1471-2105-12-35

102 Blake, J. A. et al. Mouse Genome Database (MGD): Knowledgebase for mouse-human comparative biology. Nucleic Acids Res 49, D981–D987 (2021). 10.1093/nar/gkaa1083

103 Seal, R. L. et al. Genenames.org: the HGNC resources in 2023. Nucleic Acids Res 51, D1003–D1009 (2023). 10.1093/nar/gkac888

104 Pedregosa, F. et al. Scikit-learn: Machine learning in Python. the Journal of machine Learning research 12, 2825–2830 (2011).

105 Granja, J. M. et al. ArchR is a scalable software package for integrative single-cell chromatin accessibility analysis. Nat Genet 53, 403–411 (2021). 10.1038/s41588-021-00790-6

106 Zhang, Y. et al. Model-based analysis of ChIP-Seq (MACS). Genome Biol 9, R137 (2008). 10.1186/gb-2008-9-9-r137

107 Stuart, T., Srivastava, A., Madad, S., Lareau, C. A. & Satija, R. Single-cell chromatin state analysis with Signac. Nat Methods 18, 1333–1341 (2021). 10.1038/s41592-021-01282-5

108 Robinson, M. D., McCarthy, D. J. & Smyth, G. K. edgeR: a Bioconductor package for differential expression analysis of digital gene expression data. Bioinformatics 26, 139–140 (2010). 10.1093/bioinformatics/btp616

109 Hoffman, G. E. & Schadt, E. E. variancePartition: interpreting drivers of variation in complex gene expression studies. BMC Bioinformatics 17, 483 (2016). 10.1186/s12859-016-1323-z

110 Lehallier, B. et al. Undulating changes in human plasma proteome profiles across the lifespan. Nat Med 25, 1843–1850 (2019). 10.1038/s41591-019-0673-2

111 Murphy, A. E., Schilder, B. M. & Skene, N. G. MungeSumstats: a Bioconductor package for the standardization and quality control of many GWAS summary statistics. Bioinformatics 37, 4593–4596 (2021). 10.1093/bioinformatics/btab665

112 Skene, N. G. & Grant, S. G. Identification of Vulnerable Cell Types in Major Brain Disorders Using Single Cell Transcriptomes and Expression Weighted Cell Type Enrichment. Front Neurosci 10, 16 (2016). 10.3389/fnins.2016.00016

113 Wang, G., Sarkar, A., Carbonetto, P. & Stephens, M. A simple new approach to variable selection in regression, with application to genetic fine mapping. J R Stat Soc Series B Stat Methodol 82, 1273–1300 (2020). 10.1111/rssb.12388

114 Wallace, C. A more accurate method for colocalisation analysis allowing for multiple causal variants. PLoS Genet 17, e1009440 (2021). 10.1371/journal.pgen.1009440

115 Coetzee, S. G., Coetzee, G. A. & Hazelett, D. J. motifbreakR: an R/Bioconductor package for predicting variant effects at transcription factor binding sites. Bioinformatics 31, 3847–3849 (2015). 10.1093/bioinformatics/btv470

116 Grissa, D., Junge, A., Oprea, T. I. & Jensen, L. J. Diseases 2.0: a weekly updated database of disease-gene associations from text mining and data integration. Database (Oxford) 2022 (2022). 10.1093/database/baac019

