## Supplementary Note for "Integrated multi-omics single cell atlas of the human retina"

**Supplementary Tables**

Supplementary Table S1. Summary of collected sc/snRNA-seq data for the human retinal tissues

Supplementary Table S2. Metadata information of sc/snRNA-seq samples

Supplementary Table S3. Summary of donor information for the atlas

Supplementary Table S4. Cell proportion distribution of the major classes

Supplementary Table S5. Differential gene expression analysis of snRNA-seq samples compared to scRNA-seq samples

Supplementary Table S6. Differential gene expression analysis of GB compared to BB cells

Supplementary Table S7. Top ranked genes of GB and BB using snRNA-seq and scRNA-seq datasets

Supplementary Table S8. Identified markers for BC, AC, and RGC by binary classifications

Supplementary Table S9. Annotaton of BC, AC, and RGC cell types using cross-mapping

Supplementary Table S10. Summary of collected snATAC-seq data for the human retina

Supplementary Table S11. Metadata information of snATAC-seq samples

Supplementary Table S12. Open chromatin regions identified in retinal major cell classes using snATAC-seq and MPRA results

Supplementary Table S13. Identified transcription factors and regulon specificity scores for major classes, subclasses and BC types using snRNA-seq and snATAC-seq

Supplementary Table S14. Metadata information of snRNA-seq samples from young donors

Supplementary Table S15. Results of differential gene expression analysis during aging by LMM

Supplementary Table S16. Results of differential gene expression by sliding window analysis (signed -log10(p-value))

Supplementary Table S17. Gene set enrichment results for age-, gender-, interaction between age and gender- dependent genes

Supplementary Table S18. Results of differential gene expression analysis between genders by LMM

Supplementary Table S19. Results of differential gene expression analysis of interaction between aging and gender by LMM

Supplementary Table S20. Fine-mapped genetic variants for seven GWAS traits

Supplementary Table S21. Fine-mapped eQTL variants

### Supplementary Note

#### Benchmark data integration algorithms using scIB

A major source of batch effects in our data arises from the coexistence of both snRNA-seq and scRNA-seq data (Extended Fig. 1D). Consequently, we conducted a specific benchmark to evaluate the capacity of data integration methods to integrate these assays and represent cell types, comparing this with their integration into separate references (Supplementary Fig. 1). Despite including several recently successful approaches for integrating snRNA-seq and scRNA-seq data (Methods), we observed that maintaining separate single-cell and single-nucleus reference atlases provided a more accurate representation of cellular variation in the human retina compared to their integration. For example, even our top-performing integration method still visually separates single cells and single nuclei after integration (Extended Fig. 1C). Since major cell type variation is well captured in both reference integrations, as demonstrated by marker gene expression (Extended Data Fig. 1G), both single-nucleus and single-cell references can be considered valid retina reference atlases, effectively capturing the variation within these two assay types.

Specifically, to facilitate the data integration, we benchmarked 16 different data integration methods using the scIB approach [1] (Fig. 1C and Supplementary Fig. 1). However, due to the large number of over 2 million cells, 11 methods could not be executed due to limitations in scalability of implementation and computing resources, such as the substantial memory required by many algorithms, which exceeded the limit of the programming language R. Based on an overall score derived from 14 evaluation metrics, scGen [2], scANVI [3], and scVI [4] were the top performers among the five methods that can execute in the full dataset. Although scANVI produced the highest overall scores in the majority of tested scenarios and were also top methods when grouping by technology and fraction of cells used for metric scoring (Supplementary Fig. 1), they required initial cell labels such as major cell class labels, rendering it unsuitable for subclass atlas construction. However, scVI can integrate samples without such labels, and it outperformed other label-agnostic methods in our benchmark. Therefore, we chose scVI to construct the retina atlas.

#### Transcriptome comparison between snRNA-seq and scRNA-seq

To investigate differences in the transcriptome derived from the snRNA-seq and scRNA-seq technologies, we first calculated the cell proportions of major classes in samples (Supplementary Fig. 2) and tissue regions (Extended Data Fig. 2A). A subset of snRNA-seq samples with neuronal cell types, enriched experimentally, showed high proportions of AC and RGC cells (Supplementary Fig. 2). To account for the differences in tissue regions and achieve a fair comparison of cell proportions, we compared 48 snRNA-seq and 21 scRNA-seq samples without enrichment, derived from fovea, macular, and periphery (Extended Data Fig. 2A). The cell proportions of major classes showed distinct distributions in tissue regions. While no significant differences were observed in the peripheral tissues between the two technologies, scRNA-seq captured exceptionally high proportion of muller glia cells (MGs) compared to snRNA-seq in fovea/macular tissues. This finding is consistent with the previous report that MG is overrepresented in scRNA-seq samples [5]. Although the two technologies captured distinct abundances of cell types in tissues, they showed close transcriptomic signatures of major classes (Fig. 1E). The mean area under the receiver operating characteristic (AUROC) scores in the heatmap measured the probabilities of cell types characterized by others, utilizing a neighbor-voting algorithm based on correlations among cells [6]. The high scores (e.g., AUROC > 0.9) indicated close similarities in the transcriptomic signatures for major class cell types derived from the two technologies.

In addition, we closely compared the differences of gene expression levels between the datasets of the two technologies (Methods). We identified 1,387 significantly up-regulated genes in snRNA-seq datasets (log2 fold change > 1, $q$-value < 0.05) and 3,242 up-regulated genes in scRNA-seq datasets (log2 fold change < -1, $q$-value < 0.05) regardless of cell types (Fig. 1F and Supplementary Table 5). These two sets of over-represented genes exhibited distinct enriched gene ontology biological processes (GO BPs). Specifically, the over-represented genes in the snRNA-seq datasets showed enriched BPs related to cell adhesion via plasma-membrane adhesion molecules, while the over-represented genes in scRNA-seq datasets were enriched in BPs related to ribonucleoprotein complex or ribosome biogenesis, mitochondrial gene expression, and ATP synthesis (Extended Data Fig. 2B and 2C). The highly expressed genes were consistent across cell types (Extended Data Fig. 2D and 2E), and the enriched GO BPs remained stable regardless of cell types. These findings were consistent with the previous report on the rabbit retina [5].

**Differential gene expression between giant bipolar and blue bipolar cells for snRNA-seq and scRNA-seq**

To identify novel markers for giant bipolar (GB) and blue bipolar (BB) cells, we conducted a differential gene expression analysis. Our results revealed that 341 genes were highly expressed in GB cells, while 887 genes were highly expressed in BB cells (Supplementary Table 6 and Extended Data Fig. 3D). Further analysis of these highly expressed genes showed enrichment in multiple gene ontology (GO) terms (Supplementary Fig. 3A). Specifically, the enriched biological processes (BPs) associated with GB genes were related to axon generation and development, corroborating the annotation of GB cells [7]. Similarly, the enriched BPs associated with BB genes were related to the regulation of membrane potential and synaptic transmission, aligning with the biological functions of BB cells to contact S-cone [8]. These findings provide additional evidence supporting the annotation of GB and BB cells, and the gene list obtained from our analysis offers novel and potentially valuable markers for these two distinct cell types. Moreover, based on the distribution of *AGBL1* and *SORCS3* expressions among the 14 BC cell types, we propose *AGBL1* as a potential marker gene for GB cells, and *SORCS3* as a potential marker gene for BB cells (Fig. 2A and Supplementary Fig. 3C). Regarding GB and BB in the scRNA-seq datasets, highly expressed genes revealed similar enriched GO BPs as those in the snRNA-seq datasets (Supplementary Fig. 3A and 3B). Furthermore, the expression patterns of the proposed marker genes, *AGBL1* and *SORCS3*, were found to recapitulate the expression distribution of the GB and BB cell types in both the snRNA-seq and scRNA-seq datasets (Supplementary Fig. 3C).

**Novel marker detection of bipolar cells**

The constructed BC atlas enabled the detection of comprehensive markers for human BC cell types. A binary classification approach had previously been applied to identify combinations of markers for mouse RGC cell types [9]. This approach was employed to identify novel markers for human BC cell types using the snRNA-seq data (Methods). A total of 55 markers were identified for the 14 BC cell types (Extended Data Fig. 3E and Supplementary Table 8). The dot plot depicted a cell type-specific distribution of markers across cell types. Six out of the 55 markers had previously been curated as known markers for the six cell types (Fig. 2A). The rest of the list provided refined markers for cell types. For example, *FEZF1* was used as an OFFx marker (Fig. 2A). Although it was specifically expressed in the OFFx cell type, it was relatively lowly expressed and observed in a relatively low fraction of OFFx cells. In comparison, a new marker *ARHGAP6* showed high expression across a high fraction of OFFx cells (Extended Data Fig. 3E). Therefore, the predicted novel markers provided valuable candidates for characterizing BC cell types.

**Annotation of amacrine cell clusters by known markers**

Utilizing previously characterized markers [10-12], 14 out of 73 AC clusters were annotated as known AC cell types (Extended Data Fig. 4C and Supplementary Fig. 4A). Starburst AC (SAC) is a GABAergic type, and *CHAT*, recognized as a SAC marker, was observed to be highly expressed in three clusters (HAC10, HAC26, and HAC71). In mice, *Megf10* and *Tenm3* were utilized as markers for ON-SAC and OFF-SAC, respectively[13]. This information enabled the classification of HAC10 as ON-SAC and HAC26 as OFF-SAC. Although HAC71 also exhibited a high expression of *TENM3*, it was extremely less abundant than HAC26 and was distant from it in UMAP (Fig. 3A). Therefore, HAC71 was designated as a potential OFF-SAC type (referred to as OFF-SAC*). Similarly, *GJD2* served as a marker for A2 amacrine cell (AII) and exhibited high expression in both HAC2 and HAC62. HAC2 was annotated as AII, while HAC62 was denoted as a potential AII type (AII*) due to its significantly lower abundance. *TH* was used as a marker for catecholaminergic ACs (CAI/CAII), and both HAC45 and HAC72 exhibited high *TH* expressions. Since *Chl1* positivity indicated CAI in mice, HAC45 was annotated as CAII due to its *CHL1* negativity, while HAC72 was identified as CAI because of its *CHL1* positivity. Similarities to mouse and macaque types also allowed us to tentatively assign HAC27 as a PENK AC, HAC28 as a VIP AC, HAC6 as a VG3 AC, HAC17 as a SEG AC, and HAC39, HAC46, and HAC60 as nNOS ACs (Extended Data Fig. 4D and Supplementary Fig. 4A).

**Annotation of amacrine cell clusters by cross-mapping**

To annotate AC types using public labeling, SATURN co-embedding was utilized to map 73 AC cell clusters based on snRNA-seq data to 25 AC cell types, which was based on scRNA-seq datasets[10]. Among these 25 public cell types, 17 were GABAergic ACs, and 8 were Glycinergic ACs. The cross-mapping demonstrated a confident alignment between these two sources (Extended Data Fig. 5A), and 73 snRNA-seq clusters were confidently mapped to 25 scRNA-seq types (Extended Data Fig. 5B and Supplementary Fig. 5A). A snRNA-seq cluster was annotated with an scRNA-seq type if over 90% of the cells in the cluster were mapped to that type. As a result, 42 snRNA-seq clusters were successfully mapped to 23 scRNA-seq types (Supplementary Table 9). For example, three SAC clusters (HAC10, HAC26, and HAC71) were exclusively aligned with Gaba5, which was a SAC type, and two AII clusters (HAC2 and HAC62) were exclusively mapped to Gly2, which was AII. Additionally, two “Both” clusters (HAC1 and HAC15) were mapped to Gly7, which was also previously categorized as a “Both” type. These agreements provided verification of cross-mapping between snRNA-seq clusters and scRNA-seq types by SATURN and offered additional evidence for annotating cell types in snRNA-seq clusters through novel mappings. Similarly, AC cell type labels from macaque [11] and mouse [14] species can also be used to annotate human AC clusters through SATURN co-embedding alignment. The co-embedding confidently overlaid AC types across the three species (Extended Data Fig. 5C). Specifically, 38 human ACs can be mapped to 32 out of 34 macaque AC types (Supplementary Fig. 5B and Supplementary Table 9), while 55 human ACs can be mapped to 52 out of 63 mouse AC types (Supplementary Fig. 5C and Supplementary Table 9). These cross-species mapping provided potential evidence for annotating human AC types based on known cell types in macaques and mice.

**Annotation of retinal ganglion cell clusters by cross-mapping**

Similar to the cell type annotation for ACs, public labeling of twelve RGC cell types [10] can also be utilized to annotate RGC cell types through SATURN co-embedding (Supplementary Fig. 6A). Specifically, seven RGC clusters were annotated to four RGC types via cross-mapping (Supplementary Fig. 6B and Extended Data Fig. 6B). Through cross-species analysis, human RGCs confidently overlaid with macaque and mouse RGCs (Supplementary Fig. 6C), resulting in the annotation of three human RGCs mapped to three out of 18 macaque RGCs [11] (Fig. 3D) and annotation of six human RGCs mapped to six out of 45 mouse RGCs [9] (Extended Data Fig. 6C). A recent study reported 21 orthologous types (OTs) for RGC cells [15], and five human RGC clusters can confidently be mapped to five RGC OTs (Supplementary Table 9).

From the cross-mapping results, the four most abundant primate RGC types, namely, MG_OFF, MG_ON, PG_OFF, and PG_ON, were confidently mapped to other sources. This includes four corresponding human RGC types, the four OTs, and the four macaque RGC types (Supplementary Table 9). These results confirmed the validity of the cross-mapping analysis for RGC cell types conducted via SATURN. When comparing the primate RGC types (around 18 types) [16] to the diversity in mice RGCs (45 molecularly distinct types) [9], it became evident that a one-to-one mapping between human RGC types and mice ones was uncommon. Indeed, such a one-to-one mapping was rarely observed for these four abundant types.

**Annotation of subclasses and cell types in snATAC-seq cells**

For BC class, we identified 14 BC types in 66,104 snATAC-seq cells, corresponding to the 14 cell types in snRNA-seq cells (Extended Data Fig. 8A-B). For AC, 44,544 snATAC-seq cells were clustered into 73 cell types through co-embedding, which matched with the 73 cell types of AC annotated by snRNA-seq (Extended Data Fig. 8E-F). Notably, for RGC, 29,329 snATAC-seq cells were classified as 15 cell types corresponding to the ones annotated by snRNA-seq (Supplementary Fig. 7A). Consistently, the snATAC-seq cell types matched with snRNA-seq cell types for 9,301 HC snATAC-seq cells and 9,109 Cone snATAC-seq cells as well through co-embedding (Extended Data Fig. 8G-H, Supplementary Fig. 7B). Additionally, 29,003 non-neuronal cell types that were mixed based on snATAC-seq data alone were well separated into MG, astrocyte, and microglia cell clusters when co-embedded with snRNA-seq (Fig. 4A). Overall, our study provides the most comprehensive multimodal reference cell atlas of the human retina to date.

**Annotation of subclass regulon of snATAC-seq cells**

We identified the retinal subclasses and cell type regulons. Specifically, the *RARB* regulon is highly specific for ML-Cone (Extended Data Fig. 9A). For BC subclasses, the regulons of *BACH1*, *FEZF2*, *STAT5B*, *EBF1*, and *BHLHE22* exhibit high specificity for OFF-BC, while the regulons of *MEF2C*, *ISL1*, *E2F3*, *DMBX1*, *SOX5* are specific for ON-BC (including ON CBC and RBC) (Extended Data Fig. 9B). In the case of AC subclasses, the *PBX3*, *MEIS2*, *TFAP2C*, *MAF*, and *PBX1* regulons show high specificity for GABAergic AC; Glycinergic AC is specifically governed by the regulons of *NR3C2*, *ZNF676*, *MITF*, *NFIA* and *NFIX*; and Both-AC relies on the regulons of *BCL11A*, *HMGA2*, *ZBTB18*, *NEUROD1*, *NEUROD2* (Extended Data Fig. 9C). In addition, the regulons of *LHX1*, *LHX2*, *HEY1*, *ASCL1*, and *MITF* specifically regulate HC0, whereas *ISL1* regulon takes on this role for HC1 (Extended Data Fig. 9D).

**Differential gene expression during aging**

Difference in retinal function and disease risks have been associated with individual traits such as age and sex. However, the molecular and cellular hallmarks along with the underlying mechanisms remain elusive. Furthermore, it is unclear whether aging and sex dimorphism vary across different cell types within the same tissue type. To address these questions, we investigated the impact of aging and sex dimorphism of major retinal cell classes with 135 samples from 57 donors spanning the ages from 10 to 91 years old Extended Data Fig. 10A). As a result, we identified 465 – 2,693 differentially expressed genes (DEGs) per cell class during aging utilizing a linear mixed effect model (LMM) ($q$-value $\leq$ 0.05, Fig. 6A, Extended Data Fig. 10B). The majority of the DEGs are different among cell classes (Fig. 6B). A sliding window analysis over aging further showed that most of the cell classes exhibited surges of DEGs at similar age stages, typically around the ages of 30, 60, and 80, with BC being an exception which shows a surge at around age 50 instead of 60 (Fig. 6C). For the same cell class, the majority of DEGs at the three surging ages differ and are also distinct from those identified by LMM above (Extended Data Fig. 10C). Consistently, at each of the surging age, most of the DEGs vary between cell classes (Extended Data Fig. 10C). These results suggested that although most cell classes exhibit similar dynamics pattern, the majority of DEGs during aging may be highly associated with cell type identity.

In addition, based on the DEGs, we observed the changes of both cell class-specific and shared pathways during aging. For example, the commonly activated pathways across cell classes during aging include complement and coagulation cascades, steroid hormone biosynthesis, and adaptive immune response, detected in Rod, BC, AC, and RGC (Fig. 6D-E, FDR ≤ 0.1, Supplementary Table 17). The complement pathways have been shown to play important roles in the pathogenesis of age-related macular degeneration (AMD) [17-22]. Similarly, steroid hormone receptors were found throughout the eye with these steroids being synthesized locally in ocular tissues [23], and alterations in steroid hormone homeostasis have been linked to retinal neurodegenerative disease, such as glaucoma [24]. Interestingly, the genes involved in the neuroactive ligand-receptor interaction pathway also show up-regulation over aging, which may be attributed to dysregulation of gene expression in neuronal plasticity and cellular stress resistance due to an aberrant elevation of cytoplasmic $Ca^{2+}$ levels during aging [25] (Fig. 6D-E, Extended Fig. 10D). In contrast, the common suppressed pathways include ribosome, cytoplasmic translation, mitochondrial gene expression, and ribonucleoprotein complex assembly, aligns with the finding in the fly aging study [26] (Fig. 6D). Furthermore, the suppression of oxidative phosphorylation, protein folding and modification process, ATP metabolic process, and several pathways involved in multiple neurodegeneration diseases were observed in RGC (Fig. 6D-E). These results suggest that during aging, the risks of age-related diseases may rise which may be associated with age-related expression changes of crucial genes. This offers insights into the molecular alterations associated with aging in individual cell type contexts and their potential link to diseases.

**Differential gene expression between sexes**

In addition, we also observed transcriptomic and pathway differences between male and female in individual cell class contexts. Similar to the DEGs associated with aging, the majority of DEGs between male and female are also cell class specific (Extended Data Fig. 10E). Additionally, the differential gene expression between male and female was enriched in both cell class specific and shared GO terms (Fig. 6G). For example, in photoreceptor cells, genes involved in the G protein-coupled receptor signaling pathway, regulation of membrane potential, modulation of chemical synaptic transmission, and regulation of ion transport exhibit up-regulation in males (Extended Data Fig. 10E, Fig. 6 G). This result is consistent with a synaptic sex dimorphism and might provide insight to better understand gender difference in neurological disorders[27]. Conversely, besides the up-regulation of genes associated with dosage compensation and epigenetic regulation across multiple cell types in females, immune-related genes such as those involved in cytokine-mediated signaling pathways, viral processes, and innate immune responses are also up-regulated in females specifically in MG (Extended Data Fig. 10E, Fig. 6G). This finding aligns with the observed sexual dimorphism in the mammalian immune system, where females have higher levels of immune responsiveness than males [28,29] [30,31].

More importantly, we identified DEGs and pathways exhibiting sex-dependent aging changes. For examples, genes involved in complement and coagulation cascades show more significant activation during aging in females compared to males in Cone and AC, (e.g., *A2M* and *F2RL2* in Cone, Fig. 6H-I). This result aligns with the previous studies suggesting that *F2RL2* play a role in the progression to advanced macular disease with neovascularization [32] and the higher prevalence of neovascular age-related macular degeneration in females than males [33]. Conversely, genes involved in autophagy exhibit more significant up-regulation during aging in males compared to females in RGC and AC (e.g., *ATG4A*, *CTSD*, *PRKCD*, *ULK1* in RGC, Fig. 6H-I). Consistently, autophagy has been found to play a crucial role in glaucoma [34,35], and male has higher glaucoma burden than women [36,37]. Additionally, genes involved in ribosome biogenesis show more significant down-regulation during aging in females than males across multiple cell classes (Fig. 6H-I). Overall, our results may provide insight to partially explain gender difference in certain age-related diseases.

**Example loci of fine mapping at selected GWAS loci**

Through an integrated analysis of multi-omic cell atlas, fine-mapping results, and bulk retinal eQTLs, we investigated the regulatory mechanism underlying GWAS loci. As an example, a variant (rs12531825) associated with ONL thickness was fine-mapped (PIP=0.65) to a LCRE of *GLCCI1* (Supplementary Fig. 9B). This LCRE was found to be open in multiple cell classes but most accessible in BC (i.e., a DAR of BC), consistent with the expression pattern of *GLCCI1* across retinal cell classes. Moreover, the GWAS signal is colocalized with retinal eQTL signal of

*GLCCI1* through this variant (colocalization, H4=1.00). Importantly, this variant was predicted to disrupt the binding of *RORA*, which is a transcript factor highly expressed in multiple retinal cell classes.

**Leveraging the retina atlas to study disease-specific cell types using scArches**

One of the applications of the retina atlas is the discovery of transcriptomic and genetic variation associated with disease [38]. To assess the potential of our reference atlas to infer cell types relevant in disease, we performed label transfer using scArches to learn annotations from our reference into disease datasets [39], specifically, 17 age-related macular degeneration donors sequenced with snRNA-seq (AMD). The observed distribution of uncertainties per cell type displays a bias for specific major cell classes, suggesting that specific AMD-related cells have transcriptomics deviations versus the reference (Supplementary Fig. 10A). To discard technology-specific biases, we trained models for sn- and sc-healthy cells. This allowed us to associate top-percentile (97.5%) uncertainty values with astrocyte (14% of mapped cells), and microglia (12%) and AC (7%). Few mapped RPE cells (n=142) have high uncertainties only in sc- (>50%), and not in sn- trained models, and were also considered for downstream interpretation. To interpret the transcriptomic profile of high-uncertainty cells, we stratified cells by uncertainty groups, and calculated differentially expressed genes in each of those. Top marker genes in these comparisons were enriched for a subset of 279 genes reported as AMD-biomarkers [40] (8 out of 240 genes, OR=4.4, $P < 0.01$, Fisher's exact test, one-tailed), and found to have differential expression levels in RPEs (*ACTG1, COL4A3*, *RDH5*), Astrocyte (*TRPM3*), MG (*CDH23*), Microglia (*CLU, EYS*), and AC *(PDE6A*) (Supplementary Fig. 10B). Retinal glial cells (MG, astrocytes and microglia) can contribute to AMD due to age-associated cellular changes [41,42]. This and the separation of donors associated with the fraction of high-uncertainty MG and microglia cells (Supplementary Fig. 10C), demonstrating how HRCA and label transfer allows identifying cell types and biomarkers in disease.

**References:**


1 Luecken, M. D. *et al.* Benchmarking atlas-level data integration in single-cell genomics. *Nat Methods* **19**, 41-50 (2022). https://doi.org:10.1038/s41592-021-01336-8
2 Lotfollahi, M., Wolf, F. A. & Theis, F. J. scGen predicts single-cell perturbation responses. *Nat Methods* **16**, 715-721 (2019). https://doi.org:10.1038/s41592-019-0494-8
3 Xu, C. *et al.* Probabilistic harmonization and annotation of single-cell transcriptomics data with deep generative models. *Mol Syst Biol* **17**, e9620 (2021). https://doi.org:10.15252/msb.20209620
4 Lopez, R., Regier, J., Cole, M. B., Jordan, M. I. & Yosef, N. Deep generative modeling for single-cell transcriptomics. *Nat Methods* **15**, 1053-1058 (2018). https://doi.org:10.1038/s41592-018-0229-2
5 Santiago, C. P. *et al.* Comparative Analysis of Single-cell and Single-nucleus RNA-sequencing in a Rabbit Model of Retinal Detachment-related Proliferative Vitreoretinopathy. *Ophthalmol Sci* **3**, 100335 (2023). https://doi.org:10.1016/j.xops.2023.100335
6 Crow, M., Paul, A., Ballouz, S., Huang, Z. J. & Gillis, J. Characterizing the replicability of cell types defined by single cell RNA-sequencing data using MetaNeighbor. *Nat Commun* **9**, 884 (2018). https://doi.org:10.1038/s41467-018-03282-0
7 Joo, H. R., Peterson, B. B., Haun, T. J. & Dacey, D. M. Characterization of a novel large-field cone bipolar cell type in the primate retina: evidence for selective cone connections. *Vis Neurosci* **28**, 29-37 (2011). https://doi.org:10.1017/S0952523810000374
8 Behrens, C., Schubert, T., Haverkamp, S., Euler, T. & Berens, P. Connectivity map of bipolar cells and photoreceptors in the mouse retina. *Elife* **5** (2016). https://doi.org:10.7554/eLife.20041
9 Tran, N. M. *et al.* Single-Cell Profiles of Retinal Ganglion Cells Differing in Resilience to Injury Reveal Neuroprotective Genes. *Neuron* **104**, 1039-1055 e1012 (2019). https://doi.org:10.1016/j.neuron.2019.11.006
10 Yan, W. *et al.* Cell Atlas of The Human Fovea and Peripheral Retina. *Sci Rep* **10**, 9802 (2020). https://doi.org:10.1038/s41598-020-66092-9
11 Peng, Y. R. *et al.* Molecular Classification and Comparative Taxonomics of Foveal and Peripheral Cells in Primate Retina. *Cell* **176**, 1222-1237 e1222 (2019). https://doi.org:10.1016/j.cell.2019.01.004
12 Bakken, T. E. *et al.* Single-cell and single-nucleus RNA-seq uncovers shared and distinct axes of variation in dorsal LGN neurons in mice, non-human primates, and humans. *Elife* **10** (2021). https://doi.org:10.7554/eLife.64875
13 Peng, Y. R. *et al.* Binary Fate Choice between Closely Related Interneuronal Types Is Determined by a Fezf1-Dependent Postmitotic Transcriptional Switch. *Neuron* **105**, 464-474 e466 (2020). https://doi.org:10.1016/j.neuron.2019.11.002
14 Yan, W. *et al.* Mouse Retinal Cell Atlas: Molecular Identification of over Sixty Amacrine Cell Types. *J Neurosci* **40**, 5177-5195 (2020). https://doi.org:10.1523/JNEUROSCI.0471-20.2020
15 Hahn, J. *et al.* Evolution of neuronal cell classes and types in the vertebrate retina. *bioRxiv* (2023). https://doi.org:10.1101/2023.04.07.536039
16 Kolb, H., Fernandez, E. & Nelson, R. Webvision: the organization of the retina and visual system [Internet]. (1995).
17 Park, D. H., Connor, K. M. & Lambris, J. D. The Challenges and Promise of Complement Therapeutics for Ocular Diseases. *Front Immunol* **10**, 1007 (2019). https://doi.org:10.3389/fimmu.2019.01007

18 Khandhadia, S., Cipriani, V., Yates, J. R. & Lotery, A. J. Age-related macular degeneration and the complement system. *Immunobiology* **217**, 127-146 (2012). https://doi.org:10.1016/j.imbio.2011.07.019
19 Armento, A., Ueffing, M. & Clark, S. J. The complement system in age-related macular degeneration. *Cell Mol Life Sci* **78**, 4487-4505 (2021). https://doi.org:10.1007/s00018-021-03796-9
20 Geerlings, M. J., de Jong, E. K. & den Hollander, A. I. The complement system in age-related macular degeneration: A review of rare genetic variants and implications for personalized treatment. *Mol Immunol* **84**, 65-76 (2017). https://doi.org:10.1016/j.molimm.2016.11.016
21 Ma, H., Yang, F. & Ding, X. Q. Inhibition of thyroid hormone signaling protects retinal pigment epithelium and photoreceptors from cell death in a mouse model of age-related macular degeneration. *Cell Death Dis* **11**, 24 (2020). https://doi.org:10.1038/s41419-019-2216-7
22 Xu, H. & Chen, M. Targeting the complement system for the management of retinal inflammatory and degenerative diseases. *Eur J Pharmacol* **787**, 94-104 (2016). https://doi.org:10.1016/j.ejphar.2016.03.001
23 Nuzzi, R., Scalabrin, S., Becco, A. & Panzica, G. Gonadal Hormones and Retinal Disorders: A Review. *Front Endocrinol (Lausanne)* **9**, 66 (2018). https://doi.org:10.3389/fendo.2018.00066
24 Liu, K. *et al.* Genetic variation reveals the influence of steroid hormones on the risk of retinal neurodegenerative diseases. *Front Endocrinol (Lausanne)* **13**, 1088557 (2022). https://doi.org:10.3389/fendo.2022.1088557
25 Mattson, M. P. & Arumugam, T. V. Hallmarks of Brain Aging: Adaptive and Pathological Modification by Metabolic States. *Cell Metab* **27**, 1176-1199 (2018). https://doi.org:10.1016/j.cmet.2018.05.011
26 Lu, T. C. *et al.* Aging Fly Cell Atlas identifies exhaustive aging features at cellular resolution. *Science* **380**, eadg0934 (2023). https://doi.org:10.1126/science.adg0934
27 Uhl, M., Schmeisser, M. J. & Schumann, S. The Sexual Dimorphic Synapse: From Spine Density to Molecular Composition. *Front Mol Neurosci* **15**, 818390 (2022). https://doi.org:10.3389/fnmol.2022.818390
28 Whitacre, C. C., Reingold, S. C. & O'Looney, P. A. A gender gap in autoimmunity. *Science* **283**, 1277-1278 (1999). https://doi.org:10.1126/science.283.5406.1277
29 Beeson, P. B. Age and sex associations of 40 autoimmune diseases. *Am J Med* **96**, 457-462 (1994). https://doi.org:10.1016/0002-9343(94)90173-2
30 Oertelt-Prigione, S. The influence of sex and gender on the immune response. *Autoimmun Rev* **11**, A479-485 (2012). https://doi.org:10.1016/j.autrev.2011.11.022
31 Ansar Ahmed, S., Penhale, W. J. & Talal, N. Sex hormones, immune responses, and autoimmune diseases. Mechanisms of sex hormone action. *Am J Pathol* **121**, 531-551 (1985).
32 Huan, T. *et al.* Identifying Novel Genes and Variants in Immune and Coagulation Pathways Associated with Macular Degeneration. *Ophthalmol Sci* **3**, 100206 (2023). https://doi.org:10.1016/j.xops.2022.100206
33 Rudnicka, A. R. *et al.* Age and gender variations in age-related macular degeneration prevalence in populations of European ancestry: a meta-analysis. *Ophthalmology* **119**, 571-580 (2012). https://doi.org:10.1016/j.ophtha.2011.09.027
34 Li, M. *et al.* Autophagy in glaucoma pathogenesis: Therapeutic potential and future perspectives. *Front Cell Dev Biol* **10**, 1068213 (2022). https://doi.org:10.3389/fcell.2022.1068213

35 Dixon, A. *et al.* Autophagy deficiency protects against ocular hypertension and neurodegeneration in experimental and spontanous glaucoma mouse models. *Cell Death Dis* **14**, 554 (2023). https://doi.org:10.1038/s41419-023-06086-3

36 Kapetanakis, V. V. *et al.* Global variations and time trends in the prevalence of primary open angle glaucoma (POAG): a systematic review and meta-analysis. *Br J Ophthalmol* **100**, 86-93 (2016). https://doi.org:10.1136/bjophthalmol-2015-307223

37 Zhang, N., Wang, J., Li, Y. & Jiang, B. Prevalence of primary open angle glaucoma in the last 20 years: a meta-analysis and systematic review. *Sci Rep* **11**, 13762 (2021). https://doi.org:10.1038/s41598-021-92971-w

38 Sikkema, L. *et al.* An integrated cell atlas of the lung in health and disease. *Nat Med* **29**, 1563-1577 (2023). https://doi.org:10.1038/s41591-023-02327-2

39 Lotfollahi, M. *et al.* Mapping single-cell data to reference atlases by transfer learning. *Nat Biotechnol* **40**, 121-130 (2022). https://doi.org:10.1038/s41587-021-01001-7

40 Grissa, D., Junge, A., Oprea, T. I. & Jensen, L. J. Diseases 2.0: a weekly updated database of disease-gene associations from text mining and data integration. *Database (Oxford)* **2022** (2022). https://doi.org:10.1093/database/baac019

41 Telegina, D. V., Kozhevnikova, O. S. & Kolosova, N. G. Changes in Retinal Glial Cells with Age and during Development of Age-Related Macular Degeneration. *Biochemistry (Mosc)* **83**, 1009-1017 (2018). https://doi.org:10.1134/S000629791809002X

42 Hata, M. *et al.* Early-life peripheral infections reprogram retinal microglia and aggravate neovascular age-related macular degeneration in later life. *J Clin Invest* **133** (2023). https://doi.org:10.1172/JCI159757

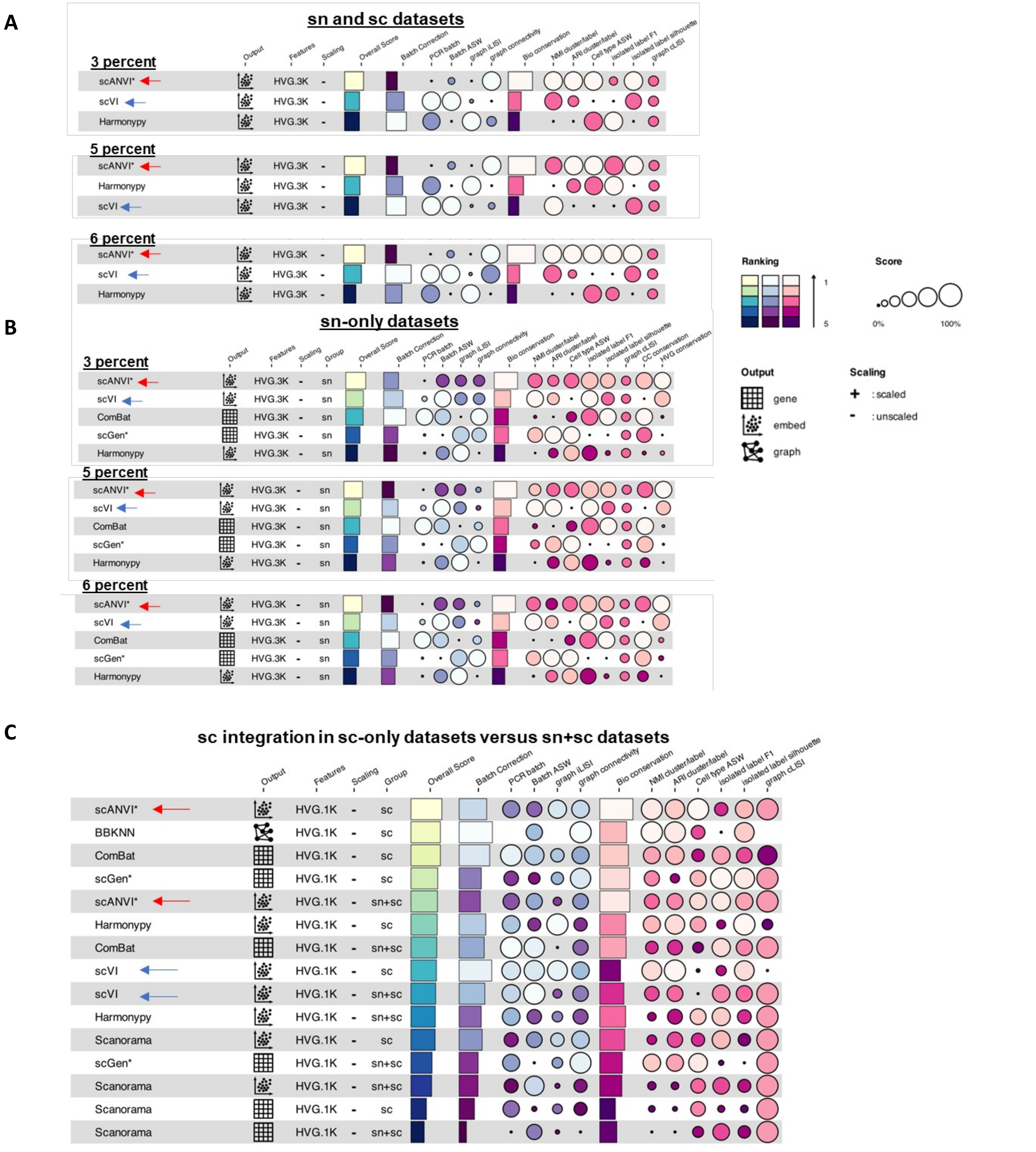


**Supplementary Figure 1. Integration benchmarking detailed results.**

A. Single-cell integration benchmark (scIB) for all sn and sc datasets. Metrics comparison for scANVI, scVI and Harmonpy are shown, using three thousand highly variable features (HVG.3K), and matched samples of 3, 5, and 6 percent of cells from the full integrated objects, for matrics calculations. scANVI and scVI are highlighted with red and blue arrows on each table, respectively. Table notations on right. B. same as A, but only using sn datasets. Legend and arrows as other panels. C. Comparison of metrics between methods for only sc-cells, after integration of sc-only datasets and sn+sc datasets, using one thousand highly-variable genes. All methods have higher overall scores for sc-cells when integrating sc-datasets alone (group=sc), versus sc+sn datasets (group=sc+sn).

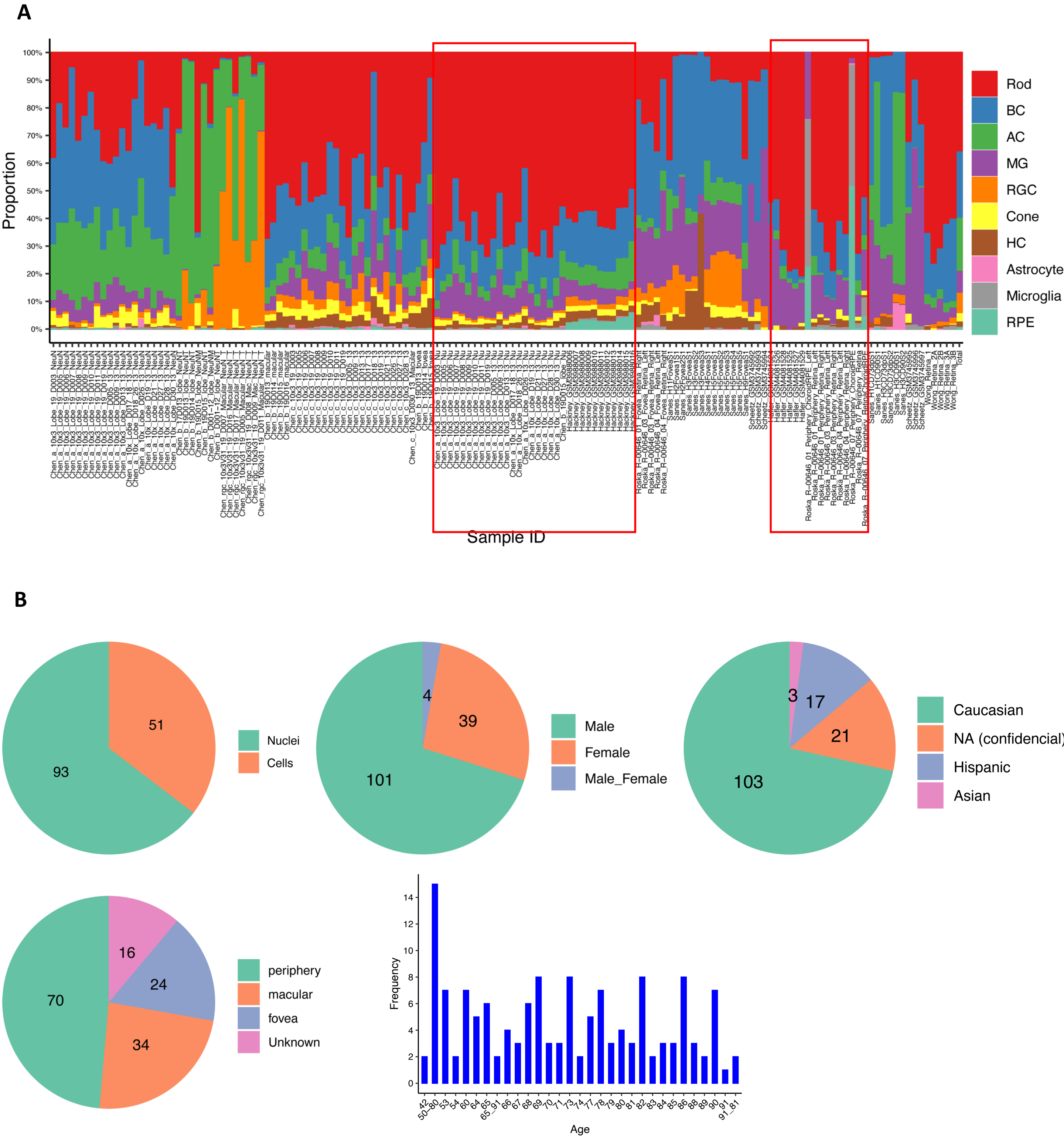


**Supplementary Figure 2. Distribution of samples**

A. Cell proportion distribution of major classes among 144 sample IDs. A subset of snRNA-seq samples without experimental enrichment was highlighted in the red box. B. The metadata distribution for the 144 samples includes information on dissociation technology, gender, race, tissue region, and age.

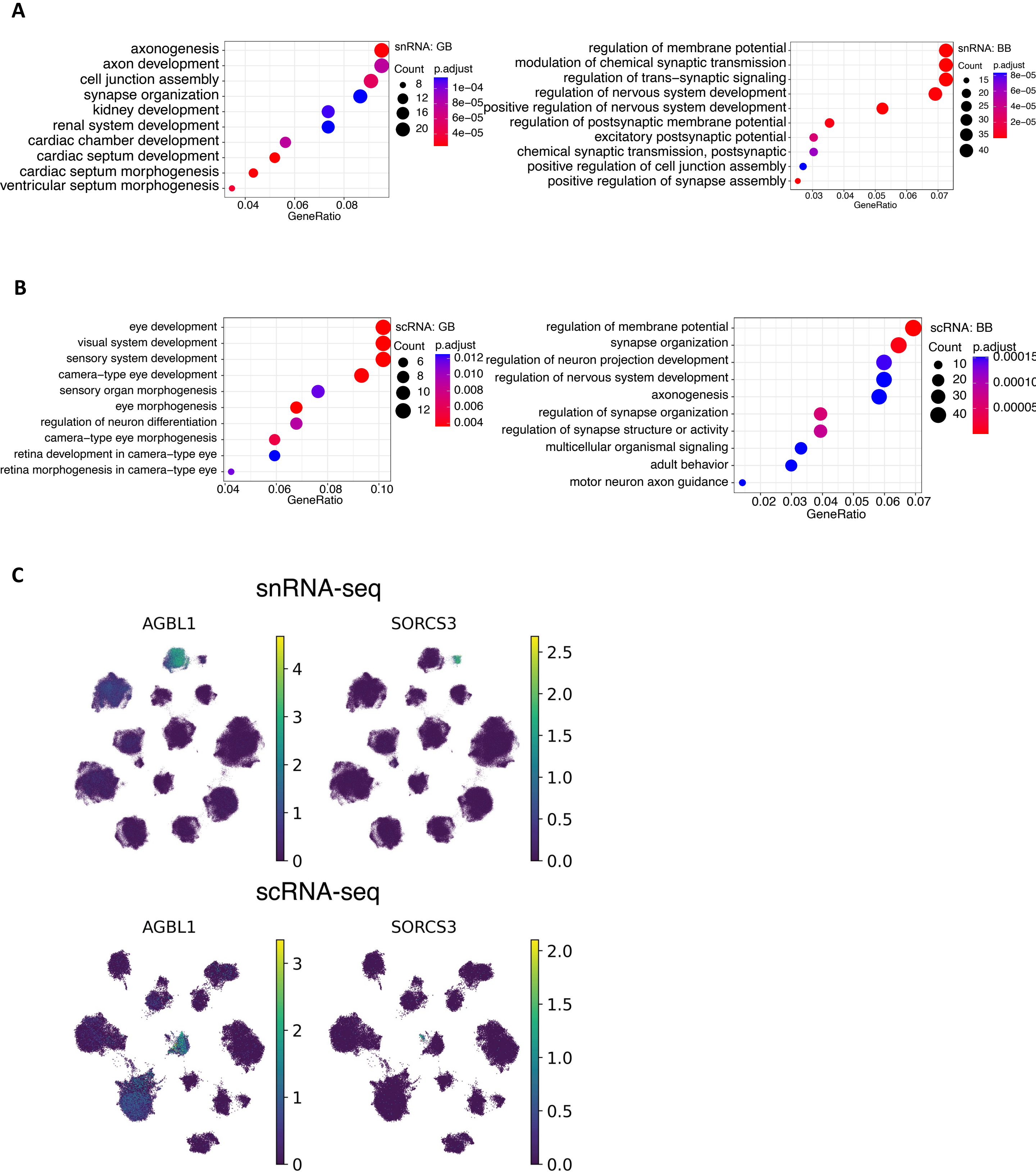


**Supplementary Figure 3. Transcriptomic signature of bipolar cells**

A. Enriched gene ontology biological processes (GO BPs) of highly expressed genes in GB and BB using the single-nuclei transcriptome data. B. Enriched GO BPs of highly expressed genes in GB and BB using single-cell measurements. C. Feature plot depicting AGBL1 and SORCS3 expression in BC clusters for snRNA-seq and scRNA-seq datasets. The color key utilized normalized and log1p transformed values.

**A**

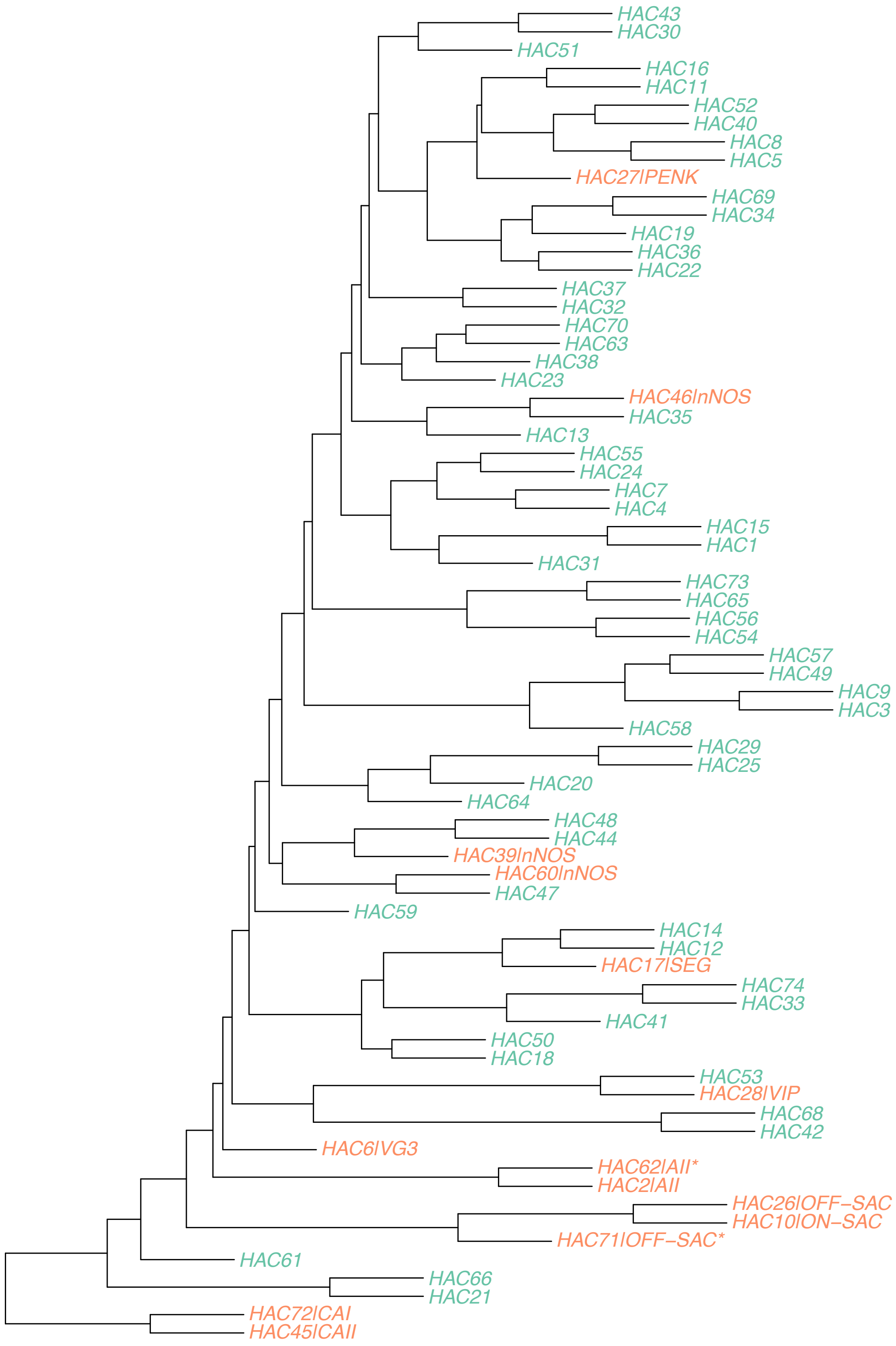


**Supplementary Figure 4. Cross-mapping of human amacrine cells.**

A. AC types in hierarchical clustering using transcriptome data. Each node represents an AC type. A node label is colored in red if its AC type has a name; otherwise, the node label is colored in green.

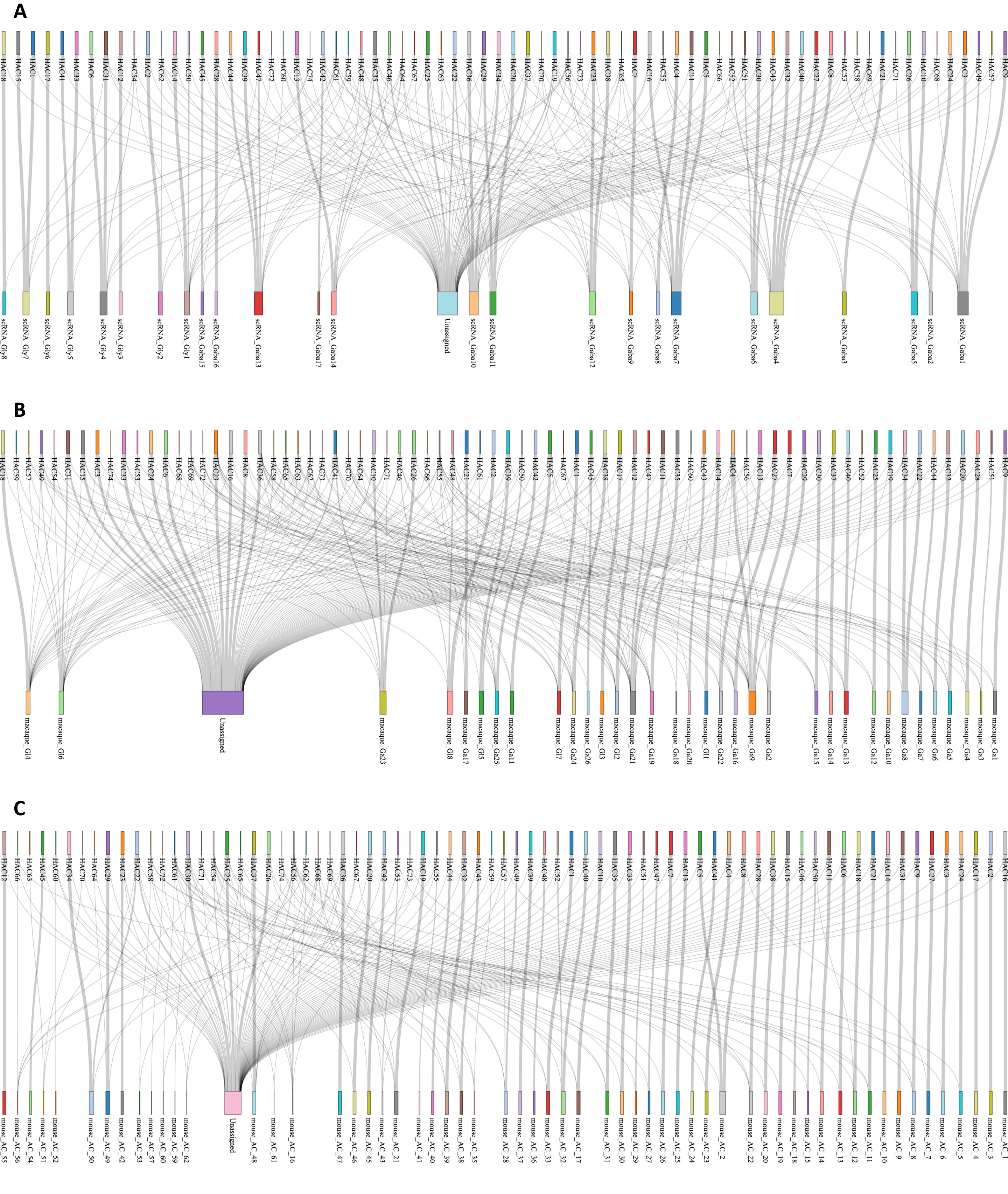


**Supplementary Figure 5. Cross-mapping of human amacrine cells.**

A. Sankey diagram depicting the relationship between AC clusters from snRNA-seq datasets and the public labeling of AC types from scRNA-seq datasets. The width of the lines is proportional to the number of cells in the mapping. B. Sankey diagram illustrating AC types alignment between humans (top row) and macaques (bottom row). C. Sankey diagram illustrating AC types alignment between humans (top row) and mice (bottom row).

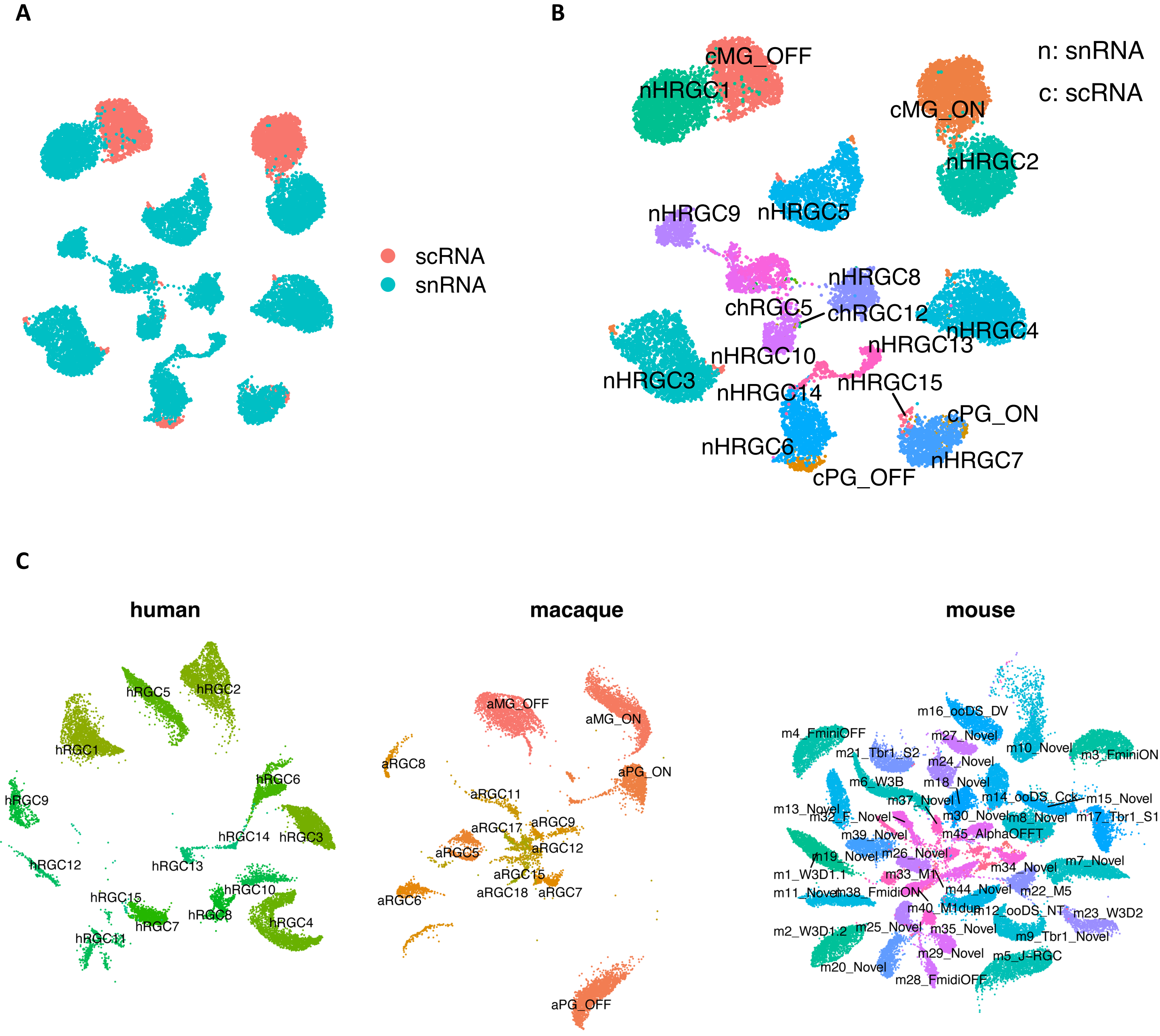


**Supplementary Figure 6. Cross-mapping of human retinal ganglion cells.**

A. SATURN co-embedding visualization of RGC cell types between snRNA-seq and scRNA-seq. RGC cells are colored by the two technologies. B. The same SATURN co-embedding with RGC type labels color-coded on top of clusters. Labels are prefixed with "n" for snRNA-seq datasets and "c" for scRNA-seq data. C. SATURN co-embedding visualization of RGC types across human, macaque and mouse species. RGC cell labels for the three species are overlaid on clusters. Labels are prefixed with "h" for human, "a" for macaque, and "m" for mouse.

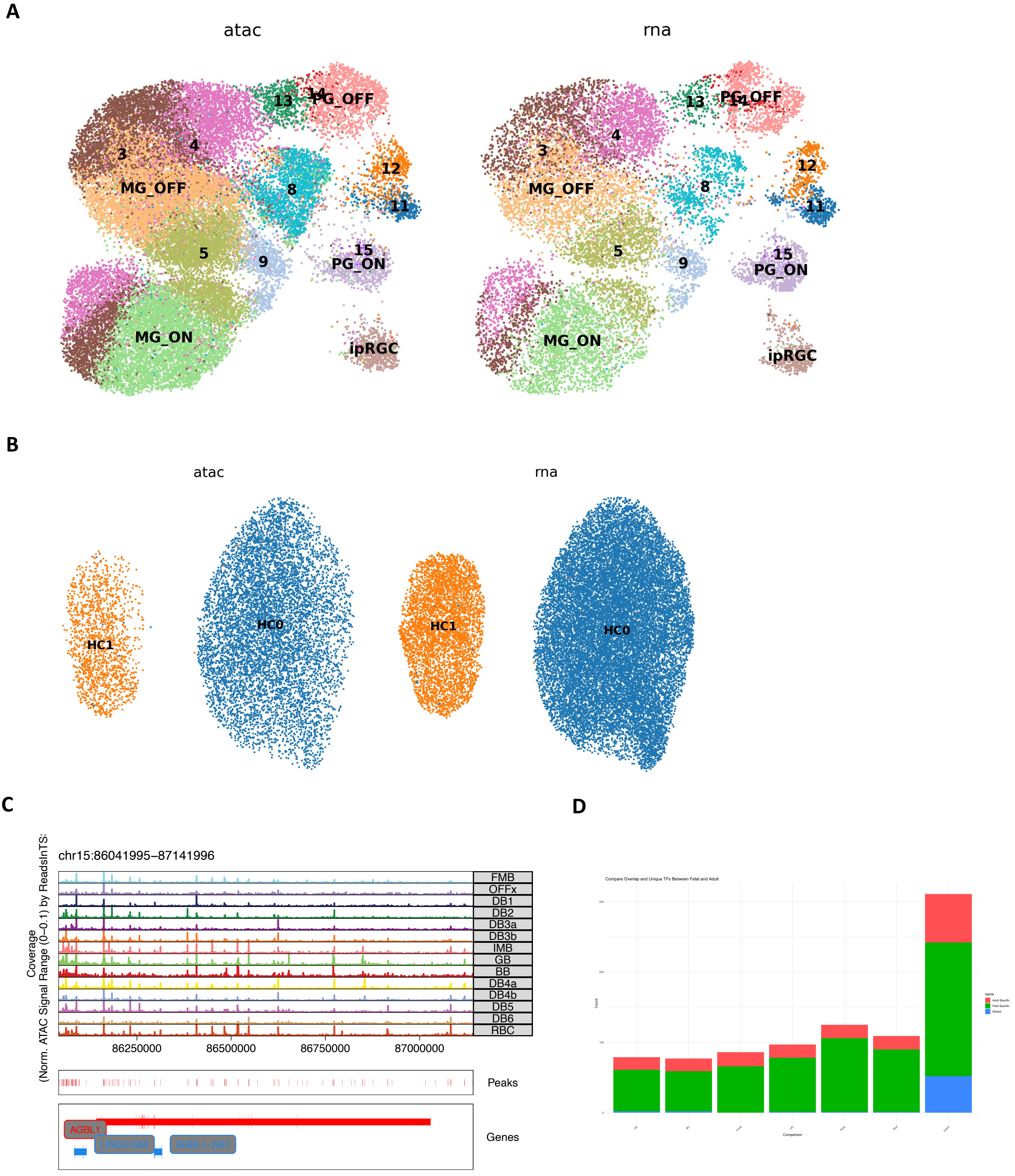


**Supplementary Figure 7. Co-embedding between snRNA-seq and snATAC-seq.**

A. UMAP showing the co-embedding of horizontal cells (RGC) from snRNA-seq and snATAC-seq were clustered in HC sub types. B. UMAP showing the co-embedding of ganglion cells (HC) from snRNA-seq and snATAC-seq were clustered in HC sub types. C. Genome track of ABGL1 of the local chromatin profile. D. Comparison of TFs identified in fetus and adult retinal cell classes.

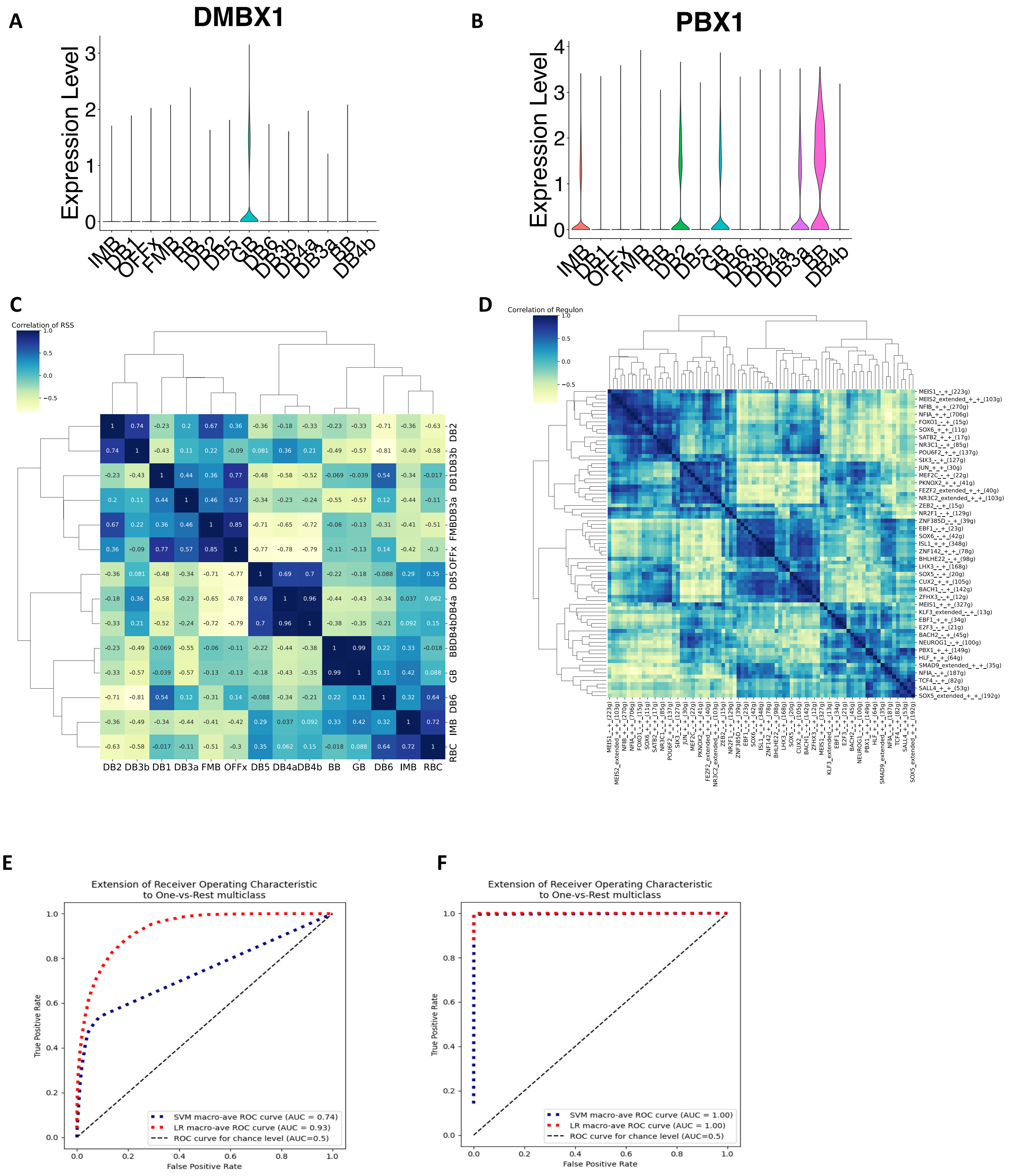


**Supplementary Figure 8. Regulons.**

A. Violin plot showing the gene expression of DMBX1 across BC cell types. B. Violin plot showing the gene expression of PBX1 across BC cell types. C. Correlation of BC cell types in terms of regulon specificity score. D. Heatmap showing the correlation of regulons based on target-gene AUC values associated with cell type identities. E. ROC-AUC of the BC cell type prediction based on TF gene expression with a logistic regression model and a SVM model. F. ROC-AUC of the BC cell type prediction based on target-gene expression with a logistic model and a SVM model

A

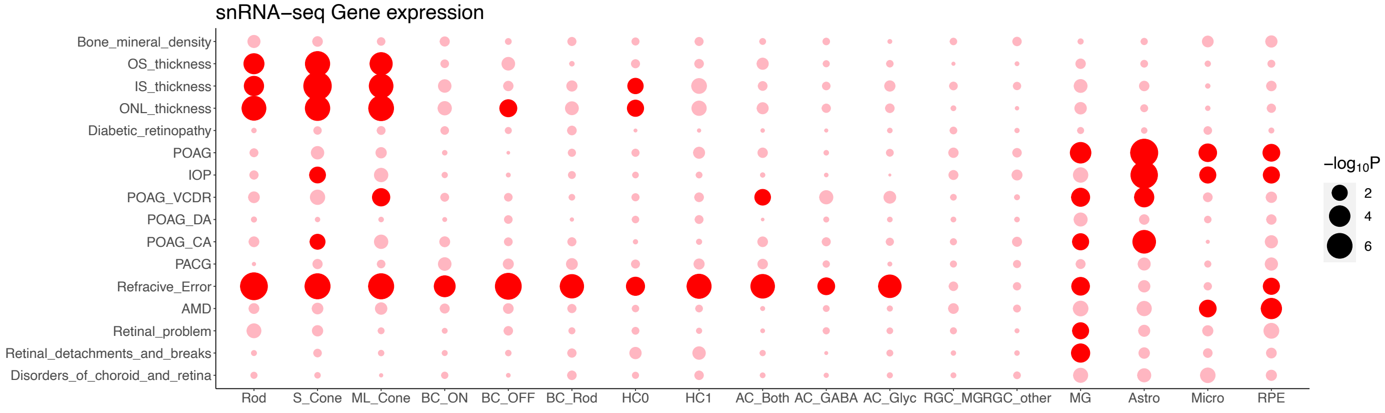


B

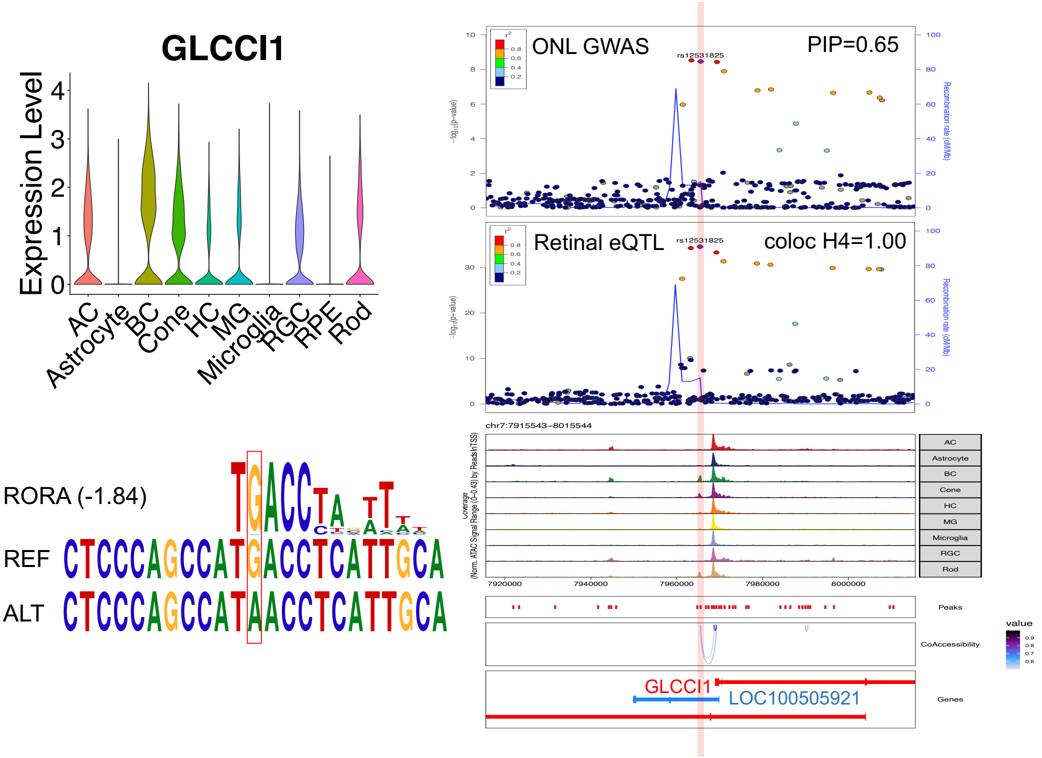


**Supplementary Figure 9. Analysis of GWAS loci**

A. Cell class/subclass enrichment of GWAS loci based on gene expression with MAGMA. Rows represent enriched GWAS traits, and columns represent retinal cell classes or subclasses. The highlight dot indicates the enrichment *q*-value < 0.05. B. Visualization of fine-mapped loci for *GLCCI1*.

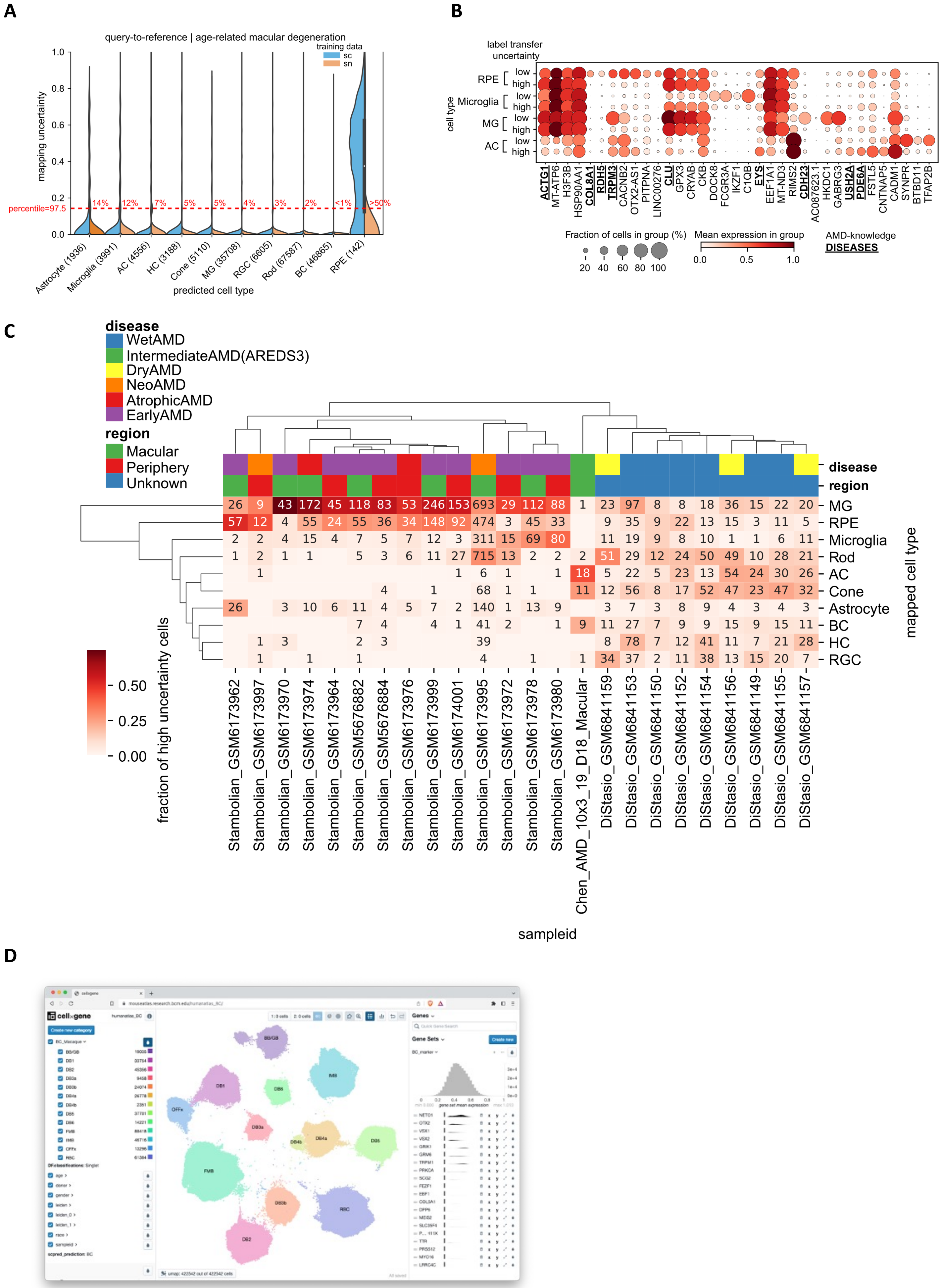
A
query-to-reference | age-related macular degeneration
training data
sc
sn
mapping uncertainty
percentile=97.5
14%
12%
7%
5%
5%
4%
3%
2%
<1%
>50%
Astrocyte (1936)
Microglia (3991)
AC (4556)
HC (3188)
Cone (5110)
MG (35708)
RGC (6605)
Rod (67587)
BC (46865)
RPE (142)
predicted cell type
B
label transfer uncertainty
cell type
RPE
Microglia
MG
AC
low
high
ACTG1
MT-ATP6
H3F3B
HSP90AA1
COL8A1
RDH5
TRPM3
CACNB2
OTX2-AS1
PITPNA
LINC00276
CLU
GPX3
CRYAB
CKB
DOCK8
FCGR3A
IKZF1
C1QB
EYS
EEF1A1
MT-ND3
RIMS2
CDH23
AC087623.1
HKDC1
GABRG3
USH2A
PDE6A
FSTL5
CNTNAP5
CADM1
SYNPR
BTBD11
TFAP2B
Fraction of cells in group (%)
Mean expression in group
AMD-knowledge
DISEASES
C
disease
WetAMD
IntermediateAMD(AREDS3)
DryAMD
NeoAMD
AtrophicAMD
EarlyAMD
region
Macular
Periphery
Unknown
MG
RPE
Microglia
Rod
AC
Cone
Astrocyte
BC
HC
RGC
mapped cell type
fraction of high uncertainty cells
Stambolian_GSM6173962
Stambolian_GSM6173997
Stambolian_GSM6173970
Stambolian_GSM6173974
Stambolian_GSM6173964
Stambolian_GSM5676882
Stambolian_GSM5676884
Stambolian_GSM6173976
Stambolian_GSM6173999
Stambolian_GSM6174001
Stambolian_GSM6173995
Stambolian_GSM6173972
Stambolian_GSM6173978
Stambolian_GSM6173980
Chen_AMD_10x3_19_D18_Macular
DiStasio_GSM6841159
DiStasio_GSM6841153
DiStasio_GSM6841150
DiStasio_GSM6841152
DiStasio_GSM6841154
DiStasio_GSM6841156
DiStasio_GSM6841149
DiStasio_GSM6841155
DiStasio_GSM6841157
sampleid
D
cellxgene

**Supplementary Figure 10. Application of HRCA.**

A. Uncertainties in AMD donor cells (n=17) per major class using *scArches* and healthy sn- and scRNA-seq datasets as training data. Predicted labels and number of cells per class are summarized on the *x*-axis. Red-dashed line indicates global percentile threshold for selection of high-uncertainty cells in each category.
B. Dot plot showing gene expression of markers genes obtained from comparisons between high/low uncertainty groups, in each cell-type, using Scanpy. Genes associated with AMD according to the DISEASES database are bold-highlighted. C. Inference and clustering of high uncertainty cell types in AMD donors. Counts per cell type for inferred cell type labels uncertainty values above percentile threshold (97.5%), clustered using fractional counts (colors) per AMD donor. Hierarchical clustering is made using average linkage. Annotations per sample as harmonized from per-study metadata. D. Screenshot of the CELLxGENE visualization for the HRCA.
